# *In vivo* Treatment of a Severe Vascular Disease via a Bespoke CRISPR-Cas9 Base Editor

**DOI:** 10.1101/2024.11.11.621817

**Authors:** Christiano R. R. Alves, Sabyasachi Das, Vijai Krishnan, Leillani L. Ha, Lauren R. Fox, Hannah E. Stutzman, Claire E. Shamber, Pazhanichamy Kalailingam, Siobhan McCarthy, Christian L. Lino Cardenas, Claire E. Fong, Takahiko Imai, Sunayana Mitra, Shuqi Yun, Rachael K Wood, Friederike M. C. Benning, Joseph Lawton, Nahye Kim, Rachel A. Silverstein, Joana Ferreira da Silva, Demitri de la Cruz, Rashmi Richa, Rajeev Malhotra, David Y. Chung, Luke H. Chao, Shengdar Q. Tsai, Casey A. Maguire, Mark E. Lindsay, Benjamin P. Kleinstiver, Patricia L. Musolino

**Author notes:** Co-first authors (Christiano R. R. Alves, Sabyasachi Das, Vijai Krishnan). Co-second authors (Leillani L. Ha, Lauren R. Fox, Hannah E. Stutzman, Claire E. Shamber, Pazhanichamy Kalailingam, Siobhan McCarthy, Christian L. Lino Cardenas).

## Abstract

Genetic vascular disorders are prevalent diseases that have diverse etiologies and few treatment options. Pathogenic missense mutations in the alpha actin isotype 2 gene (*ACTA2*) primarily affect smooth muscle cell (SMC) function and cause multisystemic smooth muscle dysfunction syndrome (MSMDS), a genetic vasculopathy that is associated with stroke, aortic dissection, and death in childhood. Here, we explored genome editing to correct the most common MSMDS-causative mutation *ACTA2* R179H. In a first-in-kind approach, we performed mutation-specific protein engineering to develop a bespoke CRISPR-Cas9 enzyme with enhanced on-target activity against the R179H sequence. To directly correct the R179H mutation, we screened dozens of configurations of base editors (comprised of Cas9 enzymes, deaminases, and gRNAs) to develop a highly precise corrective A-to-G edit with minimal deleterious bystander editing that is otherwise prevalent when using wild-type SpCas9 base editors. We then created a murine model of MSMDS that exhibits phenotypes consistent with human patients, including vasculopathy and premature death, to explore the *in vivo* therapeutic potential of this base editing strategy. Delivery of the customized base editor via an engineered SMC-tropic adeno-associated virus (AAV-PR) vector substantially prolonged survival and rescued systemic phenotypes across the lifespan of MSMDS mice, including in the vasculature, aorta, and brain. Together, our optimization of a customized base editor highlights how bespoke CRISPR-Cas enzymes can enhance on-target correction while minimizing bystander edits, culminating in a precise editing approach that may enable a long-lasting treatment for patients with MSMDS.

## Introduction

Vascular disorders underlie ∼90% of cardiovascular diseases, which are the leading cause of death and disability in the world^1^. Genetic vascular diseases are accelerated disorders that cause early severe disability and up to 10% of childhood strokes^2–6^. For example, multisystemic smooth muscle dysfunction syndrome (MSMDS) is a smooth muscle cell (SMC) myopathy characterized by severe symptoms in organs enriched with SMCs including vessels, lungs, gut, eye, and bladder, ultimately resulting in death in childhood^7,8^. Life threatening manifestations include cerebral artery stenosis, aortic enlargement^9^, and impairment of cerebrovascular autoregulation and blood-brain barrier function, which together substantially increase risk for aortic or peripheral vascular dissections and strokes in the first decade of life^6,10^. MSMDS is caused by heterozygous various missense variants at arginine 179 of the alpha actin isotype 2 (*ACTA2*) gene^7^, the most common of which is c.536G>A that causes p.Arg179His (R179H)^9^ (**Fig. 1a**), although R179C and R179S have also been observed^9^. Mutant ACTA2 R179H protein acts in a dominant negative manner to destabilize the cytoskeleton of SMCs, pericytes, and other smooth muscle actin-expressing cells, undermining their contractile function. There are no medical or surgical therapies available for patients with MSMDS that can prevent the life-threatening manifestations of stroke or vascular dissections, and, despite extensive symptomatic care, they typically die before adulthood.

**Figure 1.**
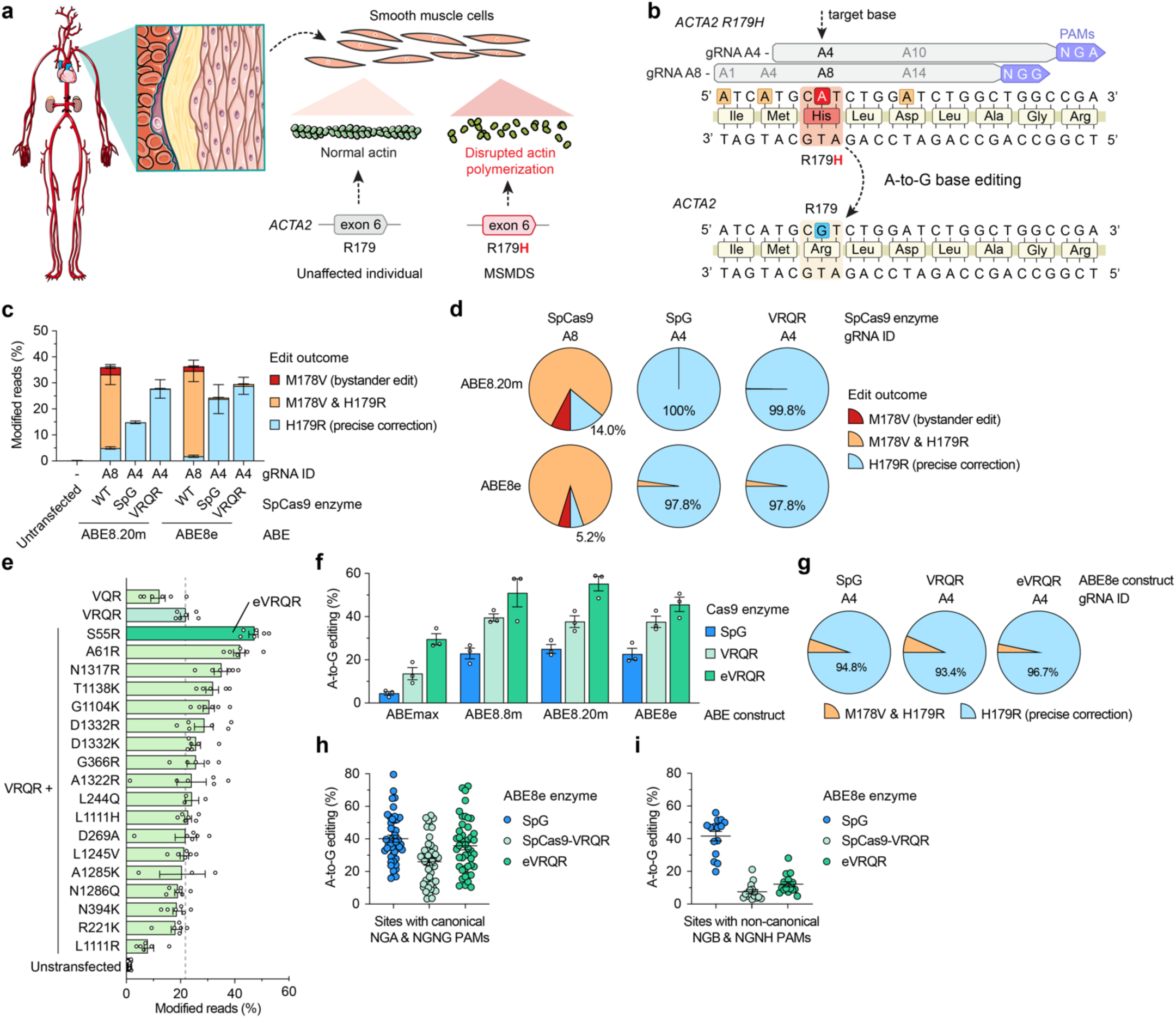
Development of a bespoke adenine base editor to correct *ACTA2* R179H. **a**, Schematic of multisystemic smooth muscle dysfunction syndrome (MSMDS) caused by an *ACTA2* R179H mutation. **b**, Schematic of the genomic region surrounding *ACTA2* R179H with base editor guide RNA (gRNA) target sites shown; potential bystander edits are shown in orange boxes. **c**, A-to-G base editing to correct *ACTA2* R179H in homozygous HEK 293T cells when using ABEs comprised of deaminase domains ABE8.20m^15^ and ABE8e^16^ fused to WT SpCas9 (with gRNA A8), or PAM variant SpCas9 enzymes SpG^18^ or SpCas9-VRQR^39^ (with gRNA A4). Base edited alleles assessed by targeted sequencing and analyzed via CRISPResso2^40^. **d**, Fraction of reads with precise *ACTA2* H179R correction with or without the M178V bystander edit, analyzed from data in **panel c**. **e**, Modified reads at the *ACTA2* R179H target site in homozygous HEK 293T cells when using SpCas9-VRQR nuclease variant enzymes harboring amino acid substitutions to potentiate on-target activity, assessed by targeted sequencing. All conditions utilized *ACTA2* R179H gRNA A4; eVRQR, enhanced SpCas9-VRQR. **f**, A-to-G base editing to correct *ACTA2* R179H in homozygous HEK 293T cells when using ABEs including ABEmax^11,38^, ABE8.8m^15^, ABE8.20m, or ABE8e fused to SpG, SpCas9-VRQR, or eVRQR when paired with *ACTA2* R179H gRNA A4. **g**, Fraction of reads with precise *ACTA2* H179R correction with or without the M178V bystander edit, analyzed from data in **panel f**. **h,i**, Summary of A-to-G base editing in HEK 293T cells with SpG, SpCas9-VRQR, or eVRQR ABE8e constructs when using gRNAs targeting sites with canonical PAMs (NGAN or NGNG; **panel h**) or non-canonical PAMs (NGBN or NGNH; **panel I**) for SpCas9-VRQR or eVRQR. B = C, G, or T; H = A, C, or T. For data in **panels c-h,** mean, s.e.m., and individual datapoints shown from experiments with between n = 3 to 6 independent biological replicates.

Given this tremendous unmet medical need, we sought to develop a genome editing approach to correct the *ACTA2* R179H mutation to provide therapeutic benefit for MSMDS patients, while also establishing a framework to treat the broader class of genetic vascular diseases. The development of CRISPR-Cas9-based base editors (BEs) has enabled the installation of nucleotide-level genetic changes^11–14^. BEs are typically comprised of a fusion of *Streptococcus pyogenes* Cas9 (SpCas9) to a nucleotide deaminase domain, and when directed by a guide RNA (gRNA) to a target genomic site can initiate specific DNA base edits. Two main classes of BEs include cytosine base editors (CBEs) that catalyze C-to-T edits^12,13^ and adenine base editors (ABEs) that catalyze A-to-G edits^11,15,16^. The deaminase domains of CBEs and ABEs typically function in a relatively narrow ‘edit window’ of approximately 6-8 bases^17^ in a portion of the Cas9 target site distal from the protospacer-adjacent motif (PAM). The narrow edit window motivates the use of engineered SpCas9 enzymes with modified or relaxed PAM requirements with increased flexibility to precisely position the deaminase over the edit-of-interest^16,18–23^.

Various types of BEs have shown promise to correct various genetic disorders caused by single nucleotide mutations^24–26^. Because *ACTA2* R179H is caused by mutation of a C•G base pair to T•A, ABEs should in principle be applicable as a therapeutic approach to correct the predominant MSMDS-causative mutation (**Fig. 1a**). Recently developed PAM-relaxed SpCas9 variant enzymes enable the targeting of non-canonical PAMs, thereby expanding the number of pathogenic mutations accessible to base editors^18–20,22,23,27–31^. Although PAM-relaxed BEs offer simplicity and ease-of-use since a small set of enzymes can be utilized to correct a wide range of disease-causing mutations, their minimal PAM requirement can also increase unwanted off-target editing due to expanded access to genomic sequences^18,23,27,32^. Instead, target-customized PAM-selective enzymes require optimization and engineering to be effective and safe, rendering them less widely developed or used despite their propensity to minimize unwanted genome-wide off-targets or to limit target-proximal bystander edits (by shifting the edit window of a base editor). For severe genetic diseases without current treatments, concerns about off-target and bystander editing are considerable, but are also balanced by the tremendous unmet need to develop safe and effective therapeutic approaches that can prolong patient life. To capitalize on the advantageous properties of precise base editors, we therefore wondered whether we could engineer a bespoke PAM-specific Cas9 enzyme using the mutant sequence as an engineering substrate, leading to an enzyme tailored to and optimized for the R179H sequence with minimized genome-wide and bystander edits.

Here we investigated the feasibility of developing a genetic treatment for MSMDS. Correction of *ACTA2* R179H using wild-type (WT) SpCas9 ABEs was efficient, but unfortunately accompanied by high nonsynonymous and deleterious bystander editing, motivating us to pursue alternate editing approaches. Protein engineering against the R179H mutant allele permitted optimization of a bespoke PAM variant base editor to precisely position the ABE edit window to maximize R179H correction while minimizing bystander editing. Comprehensive evaluation of off-target editing via multiple unbiased methods identified few off-target sites and none in genomic areas of known concern. To assess the *in vivo* effectiveness of ABE-mediated R179H correction, we developed a conditional SMC-specific mouse model of MSMDS that exhibits multiple phenotypes consistent with human symptoms including neurovascular dysfunction, aortic enlargement, intestinal dysmotility, and premature death. Delivery of the customized ABE via dual-adeno associated virus (AAV) vectors with an engineered SMC-specific AAV-PR capsid prolonged survival and rescued a range of phenotypes across the lifespan of MSMDS mice. Our results demonstrate how engineering a mutant-specific customized ABE for the R179H sequence can effectively and precisely correct the mutation *in vivo* while minimizing bystander edits, leading to substantial phenotypic recovery, and thus advancing a potential therapeutic strategy for MSMDS.

## Results

### Optimization of adenine base editing to correct the *ACTA2* R179H mutation

We first established a homozygous HEK 293T cell line bearing the *ACTA2* R179H mutation via prime editing^33^ (**Sup. Figs. 1a-e** and **Sup. Note 1**) and then investigated ABE-mediated correction of the mutation. Using this cell line, we tested ABEs paired with two different gRNAs targeted to sites harboring NGA or NGG PAMs that position the target adenine in positions A4 and A8 of the spacer, respectively (**Fig. 1b**). The ABEs were comprised of TadA8.20m^15^ or TadA8e^16^ deaminase domains fused to an engineered SpCas9 PAM variant enzyme SpCas9-VRQR that can target NGA PAMs^21,34^ (for gRNA A4), an engineered PAM-relaxed enzyme SpG that can target NGN PAMs^18^ (for gRNAs A4 and A7), or WT SpCas9 (for gRNA A8) (**Fig. 1c** and **Sup. Figs. 2a,b**). We observed effective on-target correction of R179H with various ABE and gRNA combinations, including either ABE with WT SpCas9 and gRNA A8 or SpCas9-VRQR and gRNA A4, and weaker editing with SpG and gRNA A4 (**Fig. 1c**). Analysis of bystander editing at a nearby adenine revealed high levels of A-to-G editing resulting in a nonsynonymous M178V substitution when using ABE8e-WT or ABE8.20m-WT with gRNA A8 (**Figs. 1c,d**). Use of either ABE8e-or ABE8.20m-SpCas9-VRQR and gRNA A4 resulted mainly in precise R179H correction editing with minimal bystander editing (**Figs. 1c,d**). Together, these data demonstrate how the use of SpCas9 PAM variant enzymes can achieve high levels of correction while effectively shifting the ABE edit window to minimize unwanted bystander edits.

Next, to achieve enhanced on-target R179H correction we performed R179H target-specific protein engineering of SpCas9-VRQR. We identified potential novel and previously described amino acid substitutions^18,20,21,34–36^ in SpCas9 that can augment on-target editing via energetic supplementation with non-specific protein:nucleic acid contacts^20,32,36^ (which can presumably improve editing via increased PAM interaction energetics and/or stabilization of enzyme transition states). Introduction of 18 different single amino acid substitutions into the SpCas9-VRQR nuclease led to a set of derivative enzymes that we assayed for on-target editing against the *ACTA2* R179H sequence in our homozygous HEK 293T cell model (**Fig. 1e**). Several derivative enzymes led to enhanced on-target editing efficiencies on the *ACTA2* R179H target site compared to SpCas9-VRQR, including enzymes with S55R, A61R, N1317R, T1138K, G1104K, and D1332R substitutions (**Fig. 1e**). Characterization of the PAM requirements of these enzymes via a high-throughput PAM determination assay (HD-PAMDA^18,37^) revealed generally comparable PAM preferences between SpCas9-VRQR and most derivative enzymes (**Sup. Fig. 3**), suggesting that for some enzymes the improved editing efficiencies did not negatively impact PAM selectivity.

Given the enhanced on-target editing efficiency that we observed with SpCas9-VRQR nuclease additionally encoding the S55R amino acid substitution (hereafter named enhanced SpCas9-VRQR; eVRQR), we assessed this activity-enhanced enzyme in base editing experiments. We compared SpCas9-VRQR and eVRQR to control enzymes SpG and WT as ABEs including ABEmax^11,38^, ABE8.8m^15^, ABE8.20m^15^, or ABE8e^16^ when paired with gRNA A4. Consistent with our nuclease-based results, eVRQR ABEs resulted in higher *ACTA2* R179H correction compared to SpG or VRQR (**Fig. 1f,g**). Comparison of ABE8e constructs for SpCas9-VRQR, eVRQR, and SpG across other unrelated genomic loci bearing NGN PAMs revealed that eVRQR consistently resulted in the highest levels of on-target editing at sites within its PAM specificity (NGA or NGNG PAMs; **Fig. 1h**), while also minimizing unwanted editing at other sites encoding non-canonical NGBH PAMs compared to SpG (where B is C, G, or T, and H is A, C, or T) (**Fig. 1i** and **Sup. Fig. 4a**).

In addition to the improved on-target R179H correction with eVRQR-based ABEs, we also analyzed bystander editing. ABEs paired with *ACTA2* R179H gRNA A4 again resulted in lower levels of bystander editing compared to R179H correction when WT SpCas9 ABEs and using gRNA A8 (**Sup. Figs. 4b-h**). ABEs with gRNA A4 induced low level A-1 (M178V) and C6-to-A/T/G bystander editing (**Sup. Fig. 4d**), the latter observation consistent with previous results demonstrating that ABEs can also induce cytosine bystander editing if a C of a TC motif is in position 5 or 6 of the spacer^41–44^.

Together, these results support that eVRQR can improve on-target correction of *ACTA2* R179H, enhanced base editing with eVRQR may be generalizable to other target sites with NGA or NGNG PAMs, eVRQR preserves PAM selectivity unlike PAM relaxed enzymes, and the use of customized PAM variant enzymes may minimize bystander editing by shifting the edit window to target sites not accessible with WT SpCas9-based BEs.

### Assessment of ABE-mediated *ACTA2* R179H correction specificity

Next, we sought to characterize the potential physiological impacts resulting from bystander edits co-installed by ABEs when correcting the *ACTA2* R179H mutation. We identified several low or medium level bystander edits installed by ABEs when paired with gRNAs A4 or A8, including *ACTA2* M178V, L180M, L180V, L180L and D181G (**Fig. 2a**). Leveraging a previously validated cellular assay^45^, we generated lentiviral vectors encoding mutant cDNAs bearing *ACTA2* missense mutations and transduced primary human vascular SMCs to assess the impact of mutant cDNA overexpression (**Sup. Fig. 5a**). As a marker of cytoskeletal disfunction, we evaluated expression of the histone deacetylase HDAC9 that has been shown to be elevated within SMCs in animal and cellular model systems of genetically-triggered vascular disease as well as in patient samples^45–48^. We observed that expression of the pathogenic *ACTA2* R179H mutation resulted in elevated HDAC9 transcript levels (**Fig. 2b**) and disruption of cytoskeletal actin filaments (**Sup. Fig. 5b**), indicative of cellular stress in SMCs similar to as previously described^45–47^. Of the bystander mutants that we assayed, only M178V led to similar activation of HDAC9 and cytoskeleton disruption in primary human SMCs compared to the pathogenic R179H mutation (**Fig. 2b** and **Sup. Fig. 5b**). These results suggest that the high co-occurrence of the M178V-causing bystander edit along with R179H correction when using our initial ABE8e-WT and gRNA A8 approach would result in counterproductive editing leading to a similarly pathogenic mutation being introduced (**Figs. 1c,d** and **Sup. Fig. 4f**). Importantly, this problematic M178V bystander edit was minimized by shifting the edit window via the use of eVRQR ABEs paired with gRNA A4 (**Fig. 1g** and **Sup. Fig. 4b**). Other low level bystander edits when using eVRQR ABEs and gRNA A4 appear to be tolerated by cells (**Fig. 2b** and **Sup. Figs. 4c,d and 5b**).

**Figure 2.**
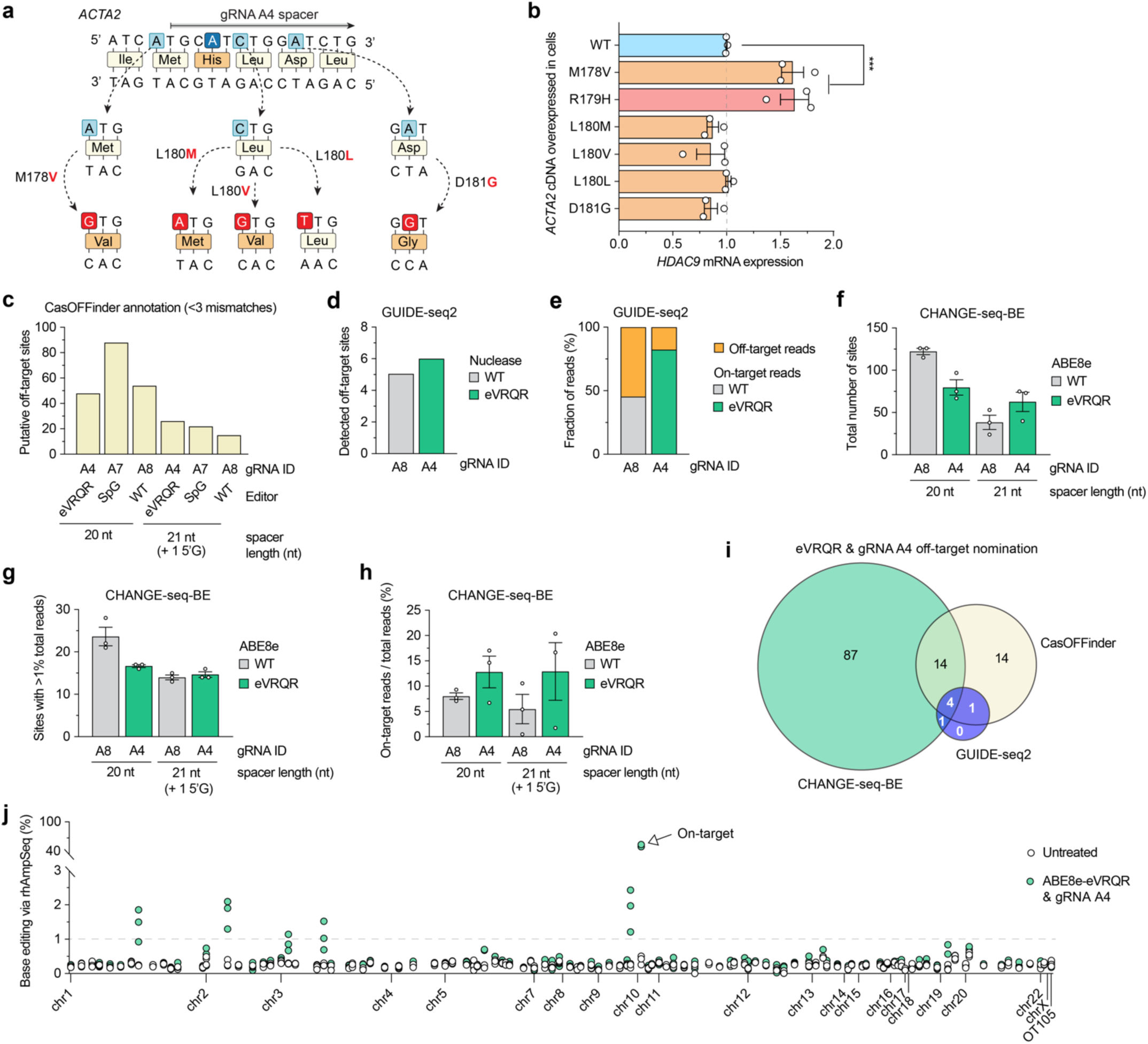
Analysis base editing specificity to correct *ACTA2* R179H. **a**, Schematic of potential *ACTA2* bystander edits induced by adenine base editors (ABEs) paired with *ACTA2* R179H gRNA A4. **b**, *HDAC9* transcript levels assessed by RT-qPCR in human smooth muscle cells (SMCs) following transduction with lentiviral vectors that express *ACTA2* variant cDNAs harboring each mutation indicated in **panel A** (see also **Sup.** Fig. 5a); wild-type (WT) *ACTA2* cDNA in blue, MSMDS-causative *ACTA2* R179H cDNA in red, and cDNAs encoding bystander edits in orange. **c,** Number of putative off-target sites in the human genome with up to 3 mismatches for the spacers of gRNAs A4 with NGAN and NGNG PAMs, A7 with NGNN PAMs, and A8 with NGGN, NAGN and NGAN PAMs, annotated by CasOFFinder^50^. **d,** Total number of GUIDE-seq2-detected off-target sites when using wild-type (WT) SpCas9 nuclease with gRNA A8 or eVRQR with gRNA A4. **e,** Percentage of total GUIDE-seq reads attributable to the on-target site or cumulative off-target sites. **f**, Total number of CHANGE-seq-BE^51^ detected off-target sites with ABE8e-WT with gRNA A8 or ABE8e-eVRQR with gRNA A4, performed using genomic DNA from patients with MSMDS. **g**, Number of CHANGE-seq-BE identified off-target sites that account for greater than 1% of total reads and are common across experiments performed using genomic DNA extracted from fibroblasts from 3 independent patients with MSMDS. **h**, Percentage of CHANGE-seq-BE reads detected at the on-target site relative to the total number of reads in each experiment. **i**, Venn diagram of nominated off-target sites with eVRQR and gRNA A4, between *in silico* CasOFFinder nomination, GUIDE-seq2 (performed via plasmid expression of nucleases in cells), or CHANGE-seq-BE (performed *in vitro* using ABE8e-VRQR protein). **j**, Summary of the levels of on-and off-target base editing in homozygous HEK 293T *ACTA2* R179H cells that were untreated (naïve) or treated with ABE8e-eVRQR and gRNA A4. Genomic DNA was subjected to rhAmpSeq for the on-target site and 121 off-target sites (nominated by CasOFFinder, GUIDE-seq2, or CHANGE-seq-BE assays) with data analysis via CRISPResso2^40^ for n = 3 independent biological replicates; base editing efficiencies plotted for only the most edited base for each target site, typically an adenine in the middle of the edit window. For panels **b** and **f-h**, mean, s.e.m, and individual datapoints shown for n = 3 independent biological replicates.

In addition to bystander edits, BEs can introduce unwanted genome-wide off-target edits. We performed both comprehensive *in silico* prediction and unbiased cell-based and biochemical assays to nominate putative off-target sites. Analysis of putative off-target sites with 3 or fewer mismatches using Cas-OFFinder^49^ for eVRQR with gRNA A4 (NGA & NGNG PAMs), SpG with A7 (NGN), and WT SpCas9 with A8 (NRG, NGA) revealed 48, 88, and 54 sites, respectively, when using gRNAs with 20 nt spacers (**Fig. 2c** and **Sup. Figs. 6a-c**).

We performed two unbiased experimental assays to nominate off-target sites. First, GUIDE-seq2 is an updated version of GUIDE-seq^52,53^ with a simpler workflow that utilizes nucleases and a bait DNA molecule to identify the location of off-target DNA breaks in living cells (Lazzarotto & Li et al, *in preparation*). We performed GUIDE-seq2 in HEK 293T *ACTA2* R179H cells using eVRQR nuclease with gRNA A4 and WT SpCas9 with gRNA A8. Results from GUIDE-seq2 experiments revealed robust on-target editing and capture of the GUIDE-seq dsODN tag (**Sup. Figs. 7a,b**), with off-target analysis revealing 5 or 6 off-target sites detected WT SpCas9/A8 or for eVRQR/A4, respectively (**Fig. 2d** and **Sup. Figs. 7c,d**). With WT SpCas9 and gRNA A8, >50% of GUIDE-seq reads were attributable to off-target sites (**Fig. 2e**) and an off-target site located in an exon of the *ACTC1* gene was detected that was edited more efficiently than the R179H on-target site (**Fig. 2e** and **Sup. Fig. 7c**). Conversely, GUIDE-seq2 analysis using eVRQR with gRNA A4 resulted in >80% of reads at the on-target site that was edited ∼8.5-fold more efficiently than any off-target site (**Fig. 2e** and **Sup. Fig. 7d**). These results suggest increased on-target precision for *ACTA2* R179H targeting with eVRQR compared to WT SpCas9.

Next, we performed an unbiased biochemical base editor-specific off-target nomination assay. CHANGE-seq-BE^54^ is an adapted version of the CHANGE-seq method^55^ that utilizes purified BE proteins instead of nucleases for *in vitro* reactions on purified genomic DNA (gDNA). We performed CHANGE-seq-BE using purified ABE8e-WT^17^ or ABE8e-eVRQR (**Sup. Figs. 8a,b**), synthetic gRNAs, and gDNA from 3 independent MSMDS patient-derived fibroblast cell lines, which led to more sensitive nomination of a range of off-target sites for each ABE and gRNA combination compared to GUIDE-seq2 (**Figs. 2f-h** and **Sup. Figs. 9a-d,10-13**). The *ACTA2* on-target site was more abundant amongst the total CHANGE-seq-BE reads for ABE8e-eVRQR/A4 than ABE8e-WT/A8 (**Fig. 2h**). Of the GUIDE-seq2 nominated off-targets, 5/6 and 2/5 were also nominated by CHANGE-seq-BE for eVRQR/A4 and WT SpCas9/A8, respectively (**Fig. 2i**).

Using a list of nominated off-targets comprised of sites from CasOFFinder, GUIDE-seq2, and CHANGE-seq-BE, we performed validation experiments via rhAmpSeq pooled multiplex sequencing using genomic DNA from naïve or ABE-treated homozygous *ACTA2* R179H HEK 293T cells. Amongst gDNA samples from ABE8e-eVRQR/A4 treated cells, we observed approximately 45% R179H correction at the on-target site, and evidence of only low level off-target editing at 5/117 off-target sites (<3%) whereas the rest were at the limit of detection (**Fig. 2j** and **Sup. Fig. 14**). Off-target editing at the 5 sites occurred in likely innocuous regions of the genome including intergenic regions, putative lncRNAs, and within an intron of the *KDM4C* gene. For comparison, we performed a more limited validation analysis of off-target editing with genomic DNA from naïve or ABE8e-WT/A8-treated homozygous *ACTA2* R179H HEK 293T cells. With ABE8e-WT and gRNA A8, we observed ∼10% off-target editing at an off-target site within an exon of the *ACTC1* gene that resulted in a non-synonymous M178V amino acid substitution (**Sup. Figs. 15a-d**). Overall, our results indicate that ABE8e-eVRQR paired with gRNA A4 can effectively correct *ACTA2* R179H, is able to avoid a problematic bystander edit, and is less prone to consequential off-target editing compared to ABE8e-WT/A8.

### Development of an MSMDS animal model harboring *Acta2* R179H

To explore *in vivo* R179H correction, we generated a knock-in murine model of MSMDS that enables Cre recombinase-inducible expression of a mutant *Acta2* R179H mouse allele (*Acta2*^fl/+^). The mutant *Acta2* allele in heterozygosity was activated using Cre expressed from the SMC-specific *Myh11* promoter^56^ (**Fig. 3a**). Aortic SMCs from Myh11-Cre:*Acta2*^fl/+^ mice (MSMDS mice) demonstrate a paucity of total stress fibers in mutant cells and a lack of colocalization of α-SMA with filamentous actin stress fibers (**Sup. Fig. 16**), similar to published reports for other *ACTA2* mutations^57^. MSMDS mice demonstrated systemic phenotypes that reflect observations in human subjects with MSMDS^6,9^, including decreased survival with most mice likely dying from toxic megacolon before 6 weeks of age (**Fig. 3b**) and impaired weight gain (**Fig. 3c**). In physical activity and motor function assays, MSMDS mice have reduced performance compared to *Acta2*^fl/+^ littermates including latency to fall in rotarod assay and are relatively inactive in open field testing (**Figs. 3d,e**). Notably, MSMDS mice exhibited a severe vasculopathy akin to human MSMDS subjects^6,9^, including wider aortic diameters compared to *Acta2*^fl/+^ littermates (**Fig. 3f**), narrower distal internal carotid arteries in the brain (**Sup. Fig. 17**). Additional symptoms included dilated pupils, hydronephrosis, and distended bladder and gut, that together suggest systemic loss of SMC contractility in MSMDS mice (**Sup. Figs. 18a-d**). These findings demonstrate that MSMDS mice recapitulate the systemic, vascular and white matter phenotypic changes observed in human MSMDS patients^6,7,9,10^ and can be used as a disease model to explore *in vivo ACTA2* R179H correction via base editing.

**Figure 3.**
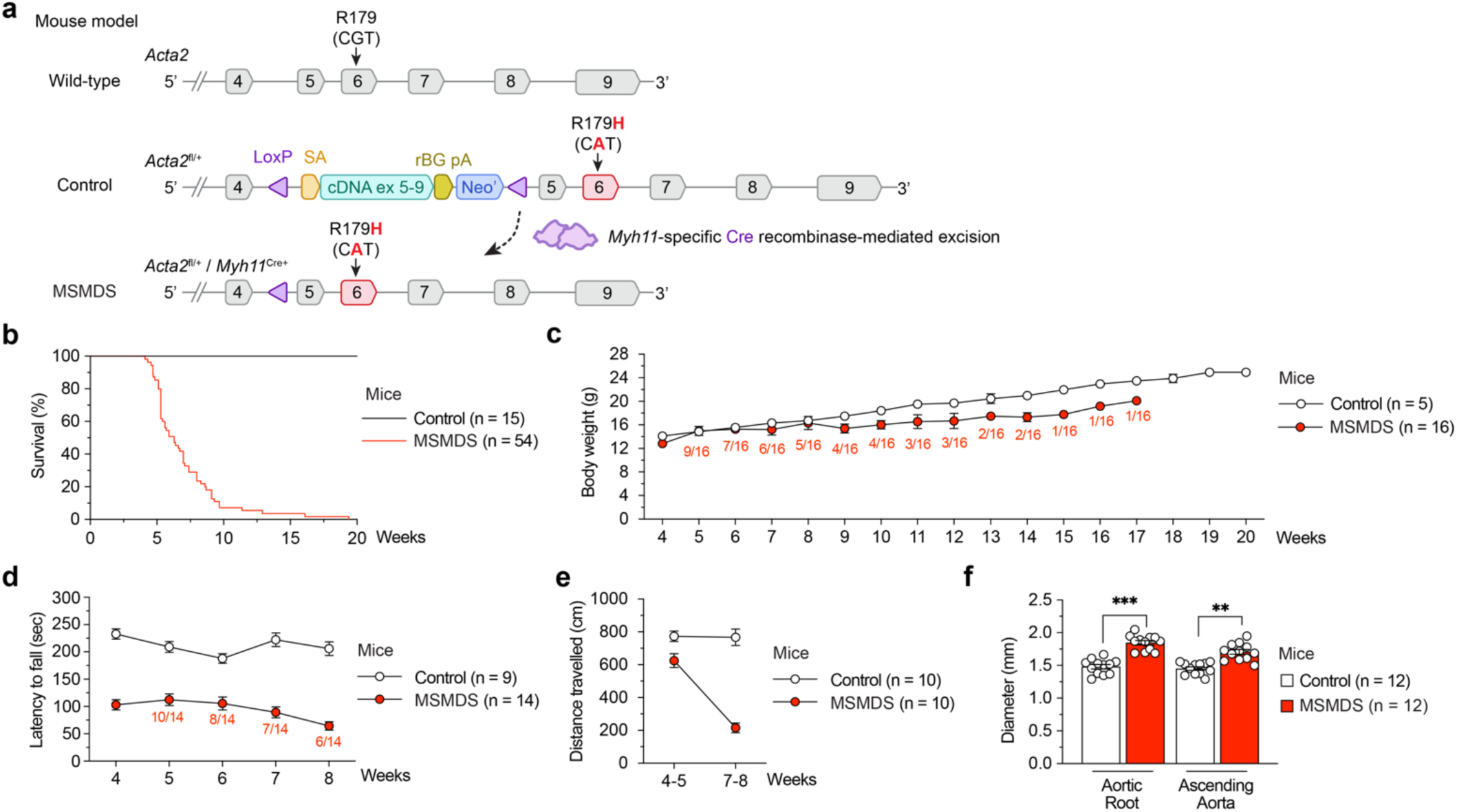
Development and characterization of an *Acta2* R179H mouse model. **a**, Schematic of the *Acta2* locus for wild-type mice, control mice (*Acta2^fl^*^/+^), and MSMDS mice (*Acta2^fl^*^/+^ / *Myh11*^Cre+^). Expression of Cre recombinase in smooth muscle cells of *Acta2^fl^*^/+^ mice crossed with *Myh11*-Cre mice via the *Myh11* promoter excises the exon 5-9 and Neo’ cassette to enable expression of the mutant R179H allele. SA, splice acceptor; ex, exon; rBH pA, Rabbit β-globin polyadenylation signal. **b-e**, Characterization of phenotypes in MSMDS mice (Myh11-Cre:Acta2^fl^) when compared to control mice, including survival (**panel b**), weight gain (**panel c**), exercise performance as demonstrated by latency to fall via rotarod assay (**panel d**), and distance traveled in open field testing (**panel e**). **f**, Comparison of aortic diameter between MSMDS and control mice at 4-5 weeks of age. Log-rank test revealed a significant difference (P < 0.001) between treated and untreated mice in **panel b**. Repeated measurement ANOVA revealed significant difference (P < 0.01) between control and MSMDS mice in **panels c-e**. T-test revealed significant difference (***P <0.001 and ** P <0.01) in **panel f**. Mean and s.e.m. shown in **panels c-f**. Sample size indicated in panels **c-f** with indication of living mice under each time point, and individual datapoints shown in **panel f**.

### *In vivo* base editing in MSMDS mice

Next, we investigated the translatability of our base editing approach from *in vitro* in cells to *in vivo* in MSMDS mice. To simulate *in vivo* conditions where ABE expression might be less optimal from AAV vectors compared to our previous results with plasmid expression in cells, we performed titration experiments in R179H HEK 293T cells to establish dynamic range amongst constructs. Across all plasmid doses, we observed that ABE8e-eVRQR and ABE8.20m-eVRQR with gRNA A4 resulted in the highest levels of on-target *ACTA2* R179H correction (**Sup. Figs. 19a-d**), ∼25-fold lower M178V bystander editing compared to ABE8e-WT paired with gRNA A8 (**Sup. Figs. 19e-h**), and only low levels of D171G bystander editing (**Sup. Figs. 19i-l**). ABE8e-eVRQR and gRNA A4 were therefore selected for *in vivo* studies and cloned into an intein-mediated dual-AAV plasmids similar to as previously described^22,58,59^ (**Fig. 4a**). Comparison of the dual-AAV plasmids encoding the split ABE8e-eVRQR construct to conventional plasmids in R179H HEK 293T cells led to comparable on-target R179H correction (**Figs. 4a,b**).

**Figure 4.**
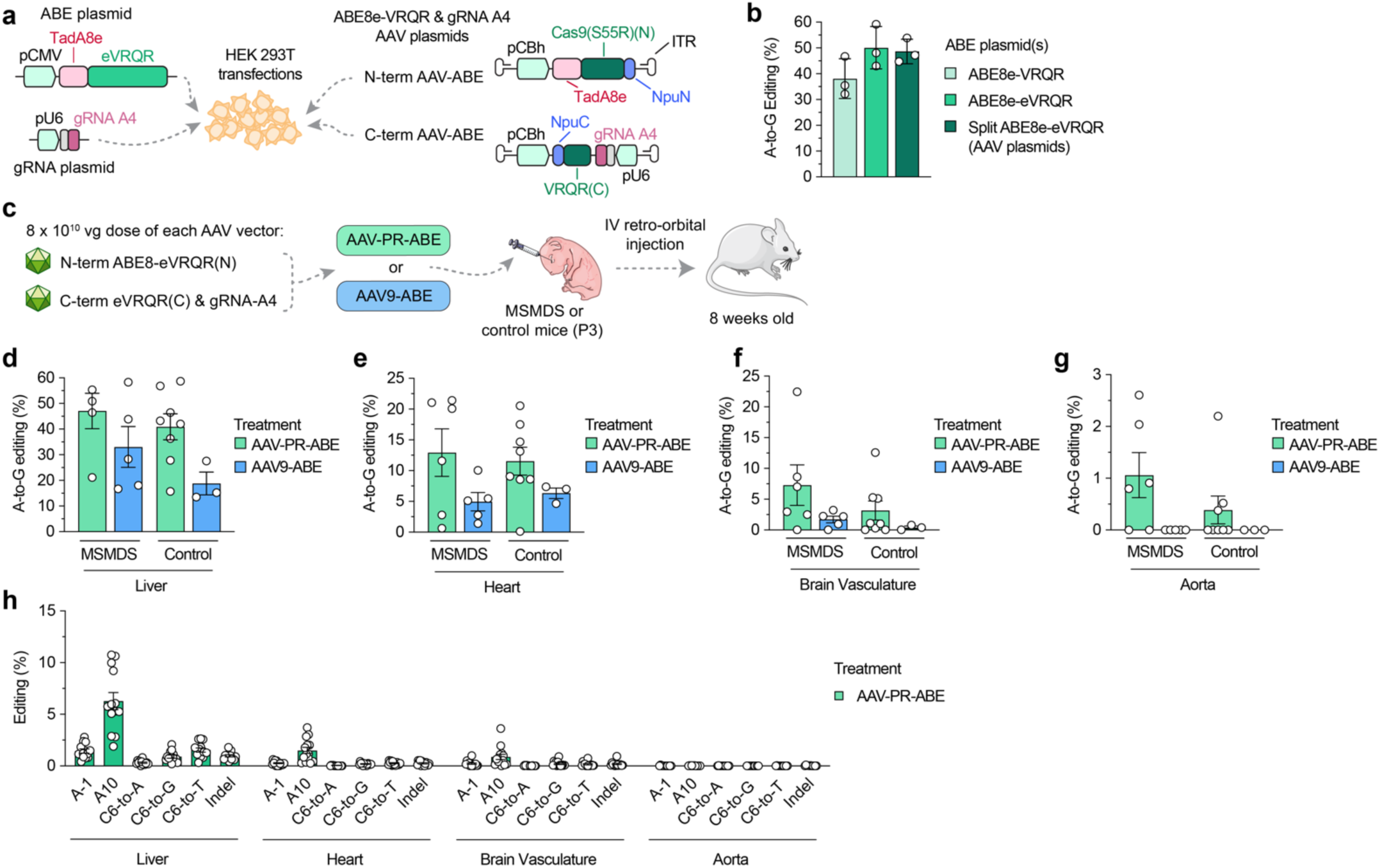
*In vivo* correction of *Acta2* R179H in MSMDS mice. **a**, Schematic of plasmid transfection experiments in HEK 293T *ACTA2* R179H cells to compare conventional ABE and gRNA expression plasmids to ITR-containing intein-split AAV production plasmids for ABE8e-eVRQR and gRNA A4. TadA8e, TadA domain from ABE8e^16^; Npu(N), N-terminal DnaE intein from *Nostoc punctiforme* (Npu)^61^; Npu(C), C-terminal Npu intein; ITR, inverted terminal repeat; Cas9(S55R)(N), residues 1-573 of nSpCas9(D10A/S55R); VRQR(C), residues 574-1,368 of SpCas9-VRQR. **b**, A-to-G base editing to correct the *ACTA2* R179H mutation in HEK 293T R179H cells via plasmid delivery. Editing assessed by targeted sequencing; mean, s.e.m., and individual datapoints shown for n = 3 independent biological replicates. **c**, Schematic of P3 intravenous (IV) injections of dual AAV-PR-ABE or AAV9-ABE vectors that express intein-split ABE8e-VRQR(S55R) and gRNA A4 into in MSMDS and control mice. Mice were sacrificed and tissues were harvested 7-8 weeks after injections. **d-g**, A-to-G editing to correct *Acta2* R179H following IV injections of dual AAV9 or AAV-PR vectors encoding ABE8e-eVRQR with gRNA A4 (AAV-ABE), with analysis of editing in the liver (**panel d**), heart (**panel e**), brain vasculature (**panel f**), or aorta (**panel g**). **h**, Summary of bystander editing or insertion or deletion mutations (indels) following P3 IV injections of AAV-PR encoding ABE8e-eVRQR with gRNA A4 into MSMDS or control mice, with editing evaluated by targeted sequencing of genomic DNA from the liver, heart, brain vasculature, or aorta. Mean, s.e.m. and individual datapoints shown in **panels d-h**.

Prior studies suggested that SMCs are the primary target cell type to treat in MSMDS^6,7,9^, which are enriched within vessels and other tissues with involuntary muscle contractions across the body. Thus, we selected two AAV serotypes to explore for systemic intravenous (IV) injections including AAV9 that mainly transduces neurons and astrocytes, and a recently engineered capsid variant AAV-PR with enhanced transduction of the vasculature including pericytes and SMCs^60^. We produced both AAV9 and AAV-PR vectors for the N-and C-terminal ABE8e-eVRQR/A4 constructs and performed IV injections of 8 x 10^10^ vector genomes (vg) in P3 control and MSMDS mice (**Fig. 4c**). Tissues samples were harvested after 7-8 weeks and gDNA was extracted and analyzed for R179H correction. We observed R179H correction in both control and MSMDS mice (control mice harbor an inactive R179H allele that remains accessible for editing) treated with either AAV9 or AAV-PR vectors across most tissues, with the highest levels of correction in liver (up to ∼60%), heart (up to ∼22%), brain vasculature (up to 23%), and aorta (<3%) (**Figs. 4d-g**, respectively). Notably, AAV-PR-ABE resulted in higher levels of R179H correction compared to AAV9 across all tissues, particularly in brain vasculature (∼4-fold) (**Figs. 4d-g** and **Sup. Figs. 20a-e**). We observed low level bystander editing at positions A-1 causing *ACTA2* M178V (**Sup. Figs. 21a-i)** or C6 causing L180M, L180V or L180L (**Sup. Figs. 22a-i**), and up to ∼12% bystander editing at A10 causing D181G (**Sup. Figs. 23a-i**), though our data suggests that the D181G bystander edit is likely to be innocuous (**Fig. 2b** and **Sup. Fig. 5b**). We observed only low levels of insertion or deletion mutations at the on-target site (**Sup. Figs. 24a-i**).

To investigate biodistribution of AAV vectors, we isolated genomic DNA and AAV genomes from liver, heart, brain vasculature, and aorta. Despite higher levels of on-target R179H correction in these tissues with AAV-PR (**Figs. 4d-g**), we observed lower genome copies in these bulk tissues from mice treated with AAV-PR compared to AAV9 (**Sup. Figs. 25a-d**), suggesting that AAV-PR encoded ABEs mediate higher R179H correction per AAV genome, perhaps via post-entry intracellular mechanisms that potentiate ABE expression (e.g. endosomal escape, capsid uncoating, or epigenetic modification of the AAV genome, as previously described^56–58^).

To gain additional general insight into the off-target profiles of our ABEs across the genome of a model organism, we nominated off-target sites via CHANGE-seq-BE using MSMDS mouse gDNA treated with ABE8e-eVRQR and gRNA A4 or ABE8e-WT and gRNA A8 (**Sup. Figs. 26a-d** and **27-30**). We observed comparable numbers of off-target sites between ABE8e-eVRQR/A4 and ABE8e-WT/A8, similar to our observations when performing CHANGE-seq-BE with human *ACTA2* R179H gDNA.

### Phenotypic rescue in MSMDS mice treated with customized ABEs

To assess potential phenotypic improvements in MSMDS mice from *in vivo* base editing, we analyzed survival in the initial cohort of injected mice. At the prespecified 8-week timepoint, all mice treated with ABE8e-eVRQR and gRNA A4 delivered via AAV-PR or AAV9 (AAV-ABE) were alive and sacrificed for tissue collection, whereas all untreated MSMDS mice were deceased (**Fig. 5a**). We then performed another experiment to analyze long-term survival, and in mice treated with AAV-PR-ABE we observed an improvement in median survival to 22.6 weeks (with survival up to 44 weeks) with AAV-PR-ABE-treated mice compared to only 6 weeks in untreated MSMDS mice (**Fig. 5b**). Death of AAV-ABE-treated mice in both treatment groups was driven by distended gut and bladder that required euthanasia and no aortic dissections or strokes were recorded. These data indicate that a single dose AAV-PR-ABE can substantially expand lifespan nearly 4-fold in this severe mouse model of MSMDS.

**Figure 5.**
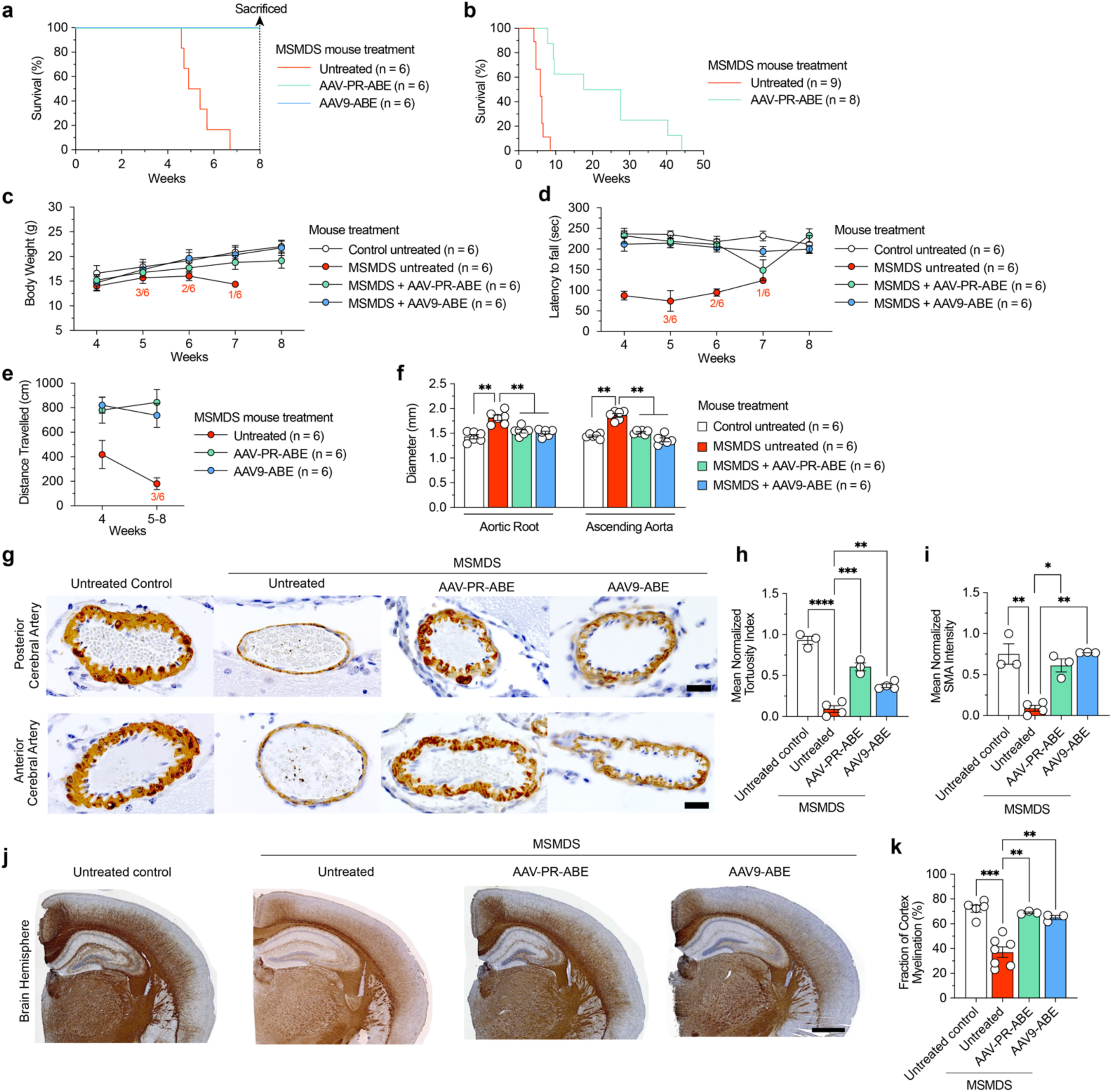
Phenotypic changes in MSMDS mice following AAV-mediated delivery of ABEs. Analyses were performed with untreated control mice (*Acta2^fl^*^/+^), untreated MSMDS mice (*Acta2^fl^*^/+^ / *Myh11*^Cre+^), and MSMDS mice treated with AAV-PR or AAV9 to deliver ABE8e-eVRQR and gRNA-A4 (AAV-ABE) at P3. **a,b**, Survival of mice at 8 weeks of age (**panel a**) or until natural death (**panel b**). **c-f,** Characterization of body mass (**panel c**), rotarod performance (**panel d**), distance traveled in open field testing (**panel e**), and aortic diameter (**panel f**). **g**, Representative images of cerebral arteries from untreated control, MSMDS, and AAV-ABE treated MSMDS mice. Sections were immunostained for smooth muscle actin (SMA) using DAB staining to visualize vascular structures. Fixed tissues were sectioned and imaged with a Zeiss LSM 800 microscope. Scale bar: 20 µm. **h-i,** Calculation of the mean normalized tortuosity index (**panel h**) and SMA intensity (**panel i**) in cerebral arteries. For tortuosity, the inner and outer perimeters of vessels were outlined using ImageJ/FIJI, with the index determined by dividing the inner perimeter by the outer perimeter. SMA intensity was measured by selecting regions of interest (ROIs) and calculating the mean pixel intensity within these regions using ImageJ/FIJI to compare SMA labeling across samples. **j,** Evaluation of neurovascular anatomy via representative coronal sections from untreated control, mutant, and treated MSMDS mice immunostained for myelin basic protein (MBP) using DAB staining and visualized with a Zeiss LSM 800 microscope. Scale bar: 500 µm. **k,** Cortical myelination assessed by measuring the total area of the cortex and the area of myelinated cortex using ImageJ/FIJI. The proportion of myelinated cortex was calculated as the percentage of the myelinated area relative to the total cortical area. Log-rank test revealed a significant difference (P < 0.001) between treated and untreated mice in **panels a-b**. Repeated measures ANOVA revealed significant differences (P < 0.01) between treated and untreated MSMDS mice in **panels c-e**. One-way ANOVA followed by Fisher’s exact test revealed significant difference in **panels f**, **h**, **i** and **k**. *P <0.05, **P <0.01, ***P <0.001, ****P <0.0001. Mean and s.e.m. shown in **panels c-f, h-i,** and **k**. Sample size indicated in **panels c-e**, and individual datapoints shown in **panels f, h-i** and **k**.

Treatment of MSMDS mice with AAV-PR-ABE or AAV9-ABE rescued a range of additional phenotypes, including body weight (**Fig. 5c**), and locomotor and behavioral performance (**Figs. 5d-e**). Furthermore, AAV-ABE treatment reverted cerebral and histological pathology in MSMDS mice including vascular structure with correction of echocardiographic aortic diameters (**Fig. 5f**), improvement in SMC contractility by restoration of the elastin tortuosity of anterior and posterior cerebral arteries (**Figs. 5g-i**), and thalamic and hippocampal arterioles morphology (**Sup. Figs. 31a-b**). MSMDS mice develop cerebral white matter injury evidenced by significant decreased global myelination levels (**Figs. 5j,k**) and altered myelination patterns in the corpus callosum, striatum, and cerebral cortex (**Sup. Figs. 31d-h**). Importantly, AAV-ABE treatment restored normal neurovascular anatomy and myelination of the cerebral cortex (**Figs. 5j,k** and **Sup. Figs. 31d-h**). Moreover, AAV-ABE treatment rescued the morphology of kidney, lung, and liver (**Sup. Fig. 32**). Taken together, these histological analyses demonstrate that *Acta2* R179H correction can elicit a systemic improvement in MSMDS mice to mitigate the pathological changes, with a particular improvement in brain vasculature.

Lastly, to determine whether AAV treatment later in life could also provide some benefit, we performed IV injections of 8 x 10^10^ vg of each of the N-and C-terminal AAV9 vectors in P14 MSMDS mice (**Sup. Fig. 33a**). We harvested tissues at 6 weeks post-injection for analysis of R179H correction and observed detectable but lower editing when compared with the previous cohorts of mice injected at P3 (**Sup. Figs. 32b,c**). Interestingly, AAV9-ABE-treated mice at P14 demonstrated more modest phenotypic improvement, including prolonged survival (**Sup. Fig. 33d**), partially restored body mass and physical performance (**Sup. Figs. 33e-g**) as well as rescued aortic diameter (**Sup. Fig. 33h**). These data indicate that although early intervention led to the greatest therapeutic effect, treatment at a later timepoint can also improve MSMDS pathology.

## Discussion

Here we developed a mutation-specific customized base editor for efficient and precise correction of the most common MSMDS-causative mutation. Compared to experiments using conventional wild-type SpCas9 base editors, the bespoke enzyme minimized a prevalent bystander edit proximal to the target base that counteracts the phenotypic benefits of mutation correction. We demonstrated that systemic IV delivery of the ABE via a dual-AAV vector approach with a recently developed SMC-tropic capsid resulted in efficient and precise *ACTA2* R179H correction in MSMDS mice that diminished vascular, gut, motor, and physiological symptoms while also substantially improving lifespan compared to untreated MSMDS mice. We were motivated to develop a genetic treatment for MSMDS given the tremendous unmet medical need since there are no disease-modifying therapies to prevent the severe manifestations of stroke or vascular dissections in infants and children. Our results support that early genetic correction in MSMDS mice led to the most profound recovery, encouraging early treatment in the therapeutic window before irreversible neurologic damage that occurs during school age and early teenage years. Due to the easily observable external phenotype of congenital mydriasis and expanded access to exome sequencing, children with MSMDS are now frequently diagnosed shortly after birth. These criteria and the tremendous unmet need support the continued development of an AAV-ABE approach as a genetic medicine for MSMDS.

Genetic therapies for autosomal dominant disorders like MSMDS necessitate different approaches when compared to gene or protein replacement therapies that have successfully treated recessive conditions. For instance, precise correction of the pathogenic mutation can restore normal physiology without risks of transgene overexpression. Our approach of directly and precisely correcting the MSMDS-causative dominant-negative mutation was enabled by the optimization of a customized mutation-specific CRISPR-Cas enzyme. By utilizing the mutant allele as a substrate for protein engineering, we developed a bespoke enzyme with favorable on-target activity and minimal off-target edits. The growing catalog of activity-enhancing mutations should be extensible to other engineered SpCas9 enzymes^18,20,21,35,62^ or Cas orthologs^63–71^, and may motivate expanded exploration of engineering mutation-customized enzymes to treat specific disease-causing sequences. Future engineering of efficacious enzymes with more narrow and specific PAM requirements should reduce the reliance on PAM-relaxed enzymes to access genetic targets for base editing, since the engineering of a large catalog of bespoke PAM-specific enzymes will offer comprehensive genome coverage with minimized off-target risk that accompanies PAM-relaxed enzymes^23,27,32^.

In this study, we observed increased survival and phenotypic recovery across relevant tissues following AAV-ABE treatment when using both AAV-PR and AAV9 capsids (**Supplementary Note 2**). In certain tissues we observed lower levels of editing in bulk samples (e.g. ∼5% *ACTA2* R179H correct with up to 20% in the brain vasculature of some mice), yet these organs demonstrated near-WT physiology (e.g. aortic diameters and neurovascular myelination). The discrepancy between editing levels and phenotypic improvement could be the result of editing only the few vessel-proximal cells that are most impactful to disease physiology, a low percentage of total SMCs amongst the bulk tissue that was assayed, or other currently unknown mechanisms (e.g. that the AAV-delivered ABE impacts *ACTA2* expression or splicing, or that MSMDS pathology results from specific subtypes of cells that were targeted and edited by our AAV-ABE approach). Our results are consistent with a recent study that utilized ABEs to modulate splicing in both cellular and mouse models of Duchenne muscular dystrophy, achieving less than 20% on-target editing in bulk tissue but restored mRNA splicing and dystrophin protein levels^72^. Additional studies will be necessary to reconcile these observations.

There are potential limitations of our study. With base editing, mutation-proximal bystander edits and genome-wide off-target edits can occur. However, the biologic implications of these edits are unclear and long-term monitoring of patients will be required to understand potential risks, motivating the development of assays to more carefully understand the impact of off-target edits. Furthermore, the size constraints of the AAV genome necessitate splitting the large base editor coding sequence into two AAV vectors, with the expressed ABE ‘halves’ being reconstituted via intein domains within doubly transduced cells^58,59^. Continued development of BEs with smaller coding sequences that can be packaged into single AAVs^73,74^ may improve *in vivo* base editing efficiency and simplify manufacturing complexity by eliminating the need for doubly-transduced cells^75^. Also, sustained expression of the ABE from the AAV episome creates potential for genomic toxicity by increasing risk for unintended off-target editing and may result in an adaptive immune response to the ABE protein. More optimal methods to express the editor could involve temporal inactivation following a short time frame to permit editing, or alternate more transient modalities including nanoparticle-mediated mRNA delivery or via viral-like particles^76,77^.

Together, the bespoke base editor developed in our study may provide a long-lasting treatment for patients with MSMDS. More broadly, this work establishes a blueprint for how mutation-specific CRISPR-Cas enzymes paired with an engineered vascular SMC-specific AAV capsid may enable efficient and safe *in vivo* base editing to treat other genetic vascular diseases, including potentially more common disorders such as Cerebral Autosomal Dominant Arteriopathy with Subcortical Infarcts and Leukoencephalopathy (CADASIL), amyloid angiopathy, moyamoya disease, Marfan, Loeys-Dietz, or vascular Ehlers-Danlos syndromes.

## Methods

### Plasmids and oligonucleotides

Target site sequences for gRNAs are available in **Supplementary Table 2.** Plasmids used in this study are described in **Supplementary Table 3**; new plasmids related to genome editing technologies generated during this study are available through Addgene: (https://www.addgene.org/Benjamin_Kleinstiver/). Oligonucleotide sequences are available in **Supplementary Table 4**.

ABE plasmids were generated by subcloning different TadA deaminase sequences into the NotI and BglII sites of pCMV-T7-ABEmax(7.10)-VRQR-P2A-EGFP (RTW5025; Addgene plasmid 140003) via isothermal assembly^78^. Additional mutations were cloned into SpCas9-VRQR nuclease or ABE plasmids using isothermal assembly the Q5 Site-Directed Mutagenesis Kit (E0554; New England Biolabs; NEB). Expression plasmids for human U6 promoter-driven gRNAs were generated by annealing and ligating duplexed oligonucleotides corresponding to spacer sequences into BsmBI-digested pUC19-U6-BsmBI_cassette-SpCas9_gRNA (BPK1520; Addgene plasmid 65777). Npu intein-split ABE constructs were cloned into N-and C-terminal AAV plasmids (Addgene plasmids 137177 and 137178, respectively). The N-terminal vector was modified to include the TadA8e domain and optionally include the S55R mutation for SpCas9-eVRQR. The C-terminal vector was modified to encode the spacers for *ACTA2* R179H gRNAs A4 or A8 and optionally to include VRQR mutations for SpCas9-eVRQR.

### Cell culture and transfections

Human HEK 293T cells (American Type Culture Collection; ATCC) and primary fibroblasts (derived from skin biopsies from three patients with MSMDS at MGH; obtained under informed consent under protocol ID 2018P002134) were cultured in Dulbecco’s Modified Eagle Medium (DMEM) supplemented with 10% heat-inactivated FBS (HI-FBS) and 1% penicillin-streptomycin. Transfections were performed 20 hours following seeding of 2x10^4^ HEK 293T cells per well in 96-well plates. Prime editor transfections contained 70 ng prime editor expression plasmid (either PEmax or PEmax-SpG), 40 ng pegRNA plasmid, and 12.5 ng ngRNA plasmid mixed with 0.79 µL of TransIT-X2 (Mirus) in a total volume of 15 µL Opti-MEM (Thermo Fisher Scientific). Base editor transfections contained 70 ng of ABE expression plasmid and 30 ng gRNA expression plasmid (exception for specific experiments testing lower doses, which included a supplemented weight of a stuffer plasmid BPK1098 to reach 100 ng total DNA) mixed with 0.72 µL of TransIT-X2 (Mirus) in a total volume of 15 µL Opti-MEM (Thermo Fisher Scientific). Transfection mixtures were incubated for 15 minutes at room temperature and distributed across the seeded HEK 293T cells. Experiments were halted after 72 hours and genomic DNA (gDNA) was collected by discarding the media, resuspending the cells in 100 µL of quick lysis buffer (20 mM Hepes pH 7.5, 100 mM KCl, 5 mM MgCl_2_, 5% glycerol, 25 mM DTT, 0.1% Triton X-100, and 60 ng/µL Proteinase K (NEB)), heating the lysate for 6 minutes at 65 °C, heating at 98 °C for 2 minutes, and then storing at -20 °C. For experiments to generate clonal cell lines, fluorescence-activated cell sorting was performed to sort for GFP+cells. Samples of supernatant media from cell culture experiments were analyzed monthly for the presence of mycoplasma using MycoAlert PLUS (Lonza).

Primary human aortic SMCs (HAoSMC) from healthy donors were purchased from Cell Applications Inc. (catalog 354K-05a). Primary aneurysm aortic SMCs were isolated from fresh TAA tissue at the moment of surgery by standard explant of the aortic media. Mouse aortic SMCs were isolated by standard explant of the ascending section of the aortas from WT or Fbn1C1039G/+ mice. SMC identity was assessed by immunofluorescence staining of contractile markers, including SM22α, Cnn, smoothelin, and vinculin. In order to preserve cell identity, all experiments were carried out at passages 1–5. Human and murine SMCs were grown with SMC growth medium from Cell Applications Inc. (catalog 311-500).

### Next-generation sequencing and data analysis

The genome modification efficiencies of nucleases, base editors, and prime editors were determined by next-generation sequencing (NGS) using a 2-step PCR-based Illumina library construction method, similar to as previously described^18^. Briefly, genomic loci were amplified from approximately 50 ng of gDNA using Q5 High-fidelity DNA Polymerase (NEB) and the primers (**Supplementary Table 4**). PCR products were purified using paramagnetic beads prepared as previously described^65,79^. Approximately 20 ng of purified PCR-1 products were used as template for a second round of PCR (PCR-2) to add barcodes and Illumina adapter sequences using Q5 and primers (**Supplementary Table 4**) and cycling conditions of 1 cycle at 98 °C for 2 min; 10 cycles at 98 °C for 10 sec, 65 °C for 30 sec, 72 °C 30 sec; and 1 cycle at 72 °C for 5 min. PCR products were purified prior to quantification via capillary electrophoresis (Qiagen QIAxcel), normalization, and pooling. Final libraries were quantified by qPCR using the KAPA Library Quantification Kit (Complete kit; Universal) (Roche) and sequenced on a MiSeq sequencer using a 300-cycle v2 kit (Illumina). On-target genome editing activities were determined from sequencing data using CRISPResso2^80^ using parameters: CRISPResso -r1 READ1 -r2 READ2 --amplicon_seq--amplicon_name --guide_seq GUIDE -w 20 --cleavage_offset -10 for nucleases and CRISPResso -r1 READ1 -r2 READ2 --amplicon_seq --guide_seq GUIDE -w 20 --cleavage_offset -10 --base_editor_output -- conversion_nuc_from A --conversion_nuc_to G --min_frequency_alleles_around_cut_to_plot 0.001. Since amplification of *ACTA2* amplifies both wild-type R179 and mutant R179H alleles in heterozygous HEK 293T cells, MSMDS fibroblasts, and mouse model of MSMDS, final levels of editing were calculated as: ([%CGT in the treated samples] - [%CGT in the control]) / [%CAT in the control]. Otherwise, bystander edits were calculated based on their absolute levels of editing.

### Lentiviral vector production

Lentiviral vectors were produced in HEK 293T cells upon transfection with a packaging plasmid (psPAX2; Addgene plasmid 12260), an envelope plasmid (pCMV-VSVG-G; Addgene plasmid 8454) and the respective lentiviral plasmid encoding variant *ACTA2* cDNAs with different bystander editing-induced amino acid substitutions (**Supplementary Table 3**). To produce each lentiviral vector, 5 million HEK 293T cells were seeded in 10 cm dishes (Corning). 24 hours after seeding, a mix was prepared with 1 µg of pCMV-VSVG-G plasmid, 2 µg of psPAX2 plasmid, and 4 µg of the lentiviral vector expressing the *ACTA2* cDNA variant in a final volume of 200 µL Opti-MEM (Fisher Scientific) and vortexed gently for 10 seconds. 50 µL of PEI MAX (Fisher Scientific) at the stock concentration of 1 mg/mL dissolved in water (pH 7.1) was added to each DNA mix and vortexed gently for 10 seconds. The DNA:PEI mix was incubated at room-temperature for 20 minutes. After incubation, the DNA mixes were each added to 8 mL aliquots of DMEM medium supplemented with 10% FBS and 1% penicillin-streptomycin, and the transfection was performed by replacing the medium on the seeded cells. Approximately 60 hours post-transfection, lentiviral particles were isolated through differential centrifugation, where conditioned medium was collected and centrifuged at 2,000 g for 5 minutes to remove cells and cell debris, and lentiviral particles were then concentrated through ultracentrifugation at 70,000g. The pellet was resuspended in 1 mL of ice-cold PBS.

### Assessment of putative phenotypic impact of bystander edits

Primary human aortic smooth muscle cells (VSMC) from healthy donors were purchased from Cell Applications Inc. (354K-05a), California, USA. Smooth muscle cell identity was assessed by immunofluorescence staining of ACTA2 and F-actin. In order to preserve cell identity, all experiments were carried out at passages 1–5. VSMCs were grown with SMC growth medium from Cell Applications Inc. (catalog 311-500). At passage 1, primary VSMCs were transduced with lentivirus overexpressing each ACTA2 variant described above. Following 72 hours post-infection, cells expressing wild-type or mutant ACTA proteins were detected using fluorescent microscopy. Slides were washed with PBS twice for 2 min and tissues were blocked with donkey serum at 10% for 1 hour followed by incubation overnight at 4°C with ACTA2 monoclonal antibody (ThermoFisher Scientific, Cat# UM870129) at 1:200 dilution and high-affinity F-actin probe (ThermoFisher Scientific, Cat# R37110) at 2 drops per mL. Slides were washed with PBS-tween at 0.1% 3 times for 3 min each followed by incubation with secondary antibody (1:400) for 1hr at room temperature. Then slides were washed with PBS-tween at 0.1%, 4 times for 3 min each and slides were mounted with diamond mounting medium containing DAPI. Slides were visualized with the Leica TCS SP8 confocal microscopy station and micrographs were digitized with the Leica Application SuiteX software. Imaging and analysis were performed using Volocity 5.2 software. Two-dimensional and white light images were analyzed using ImageJ software.

To quantify HDAC9 mRNA expression in VSMC, total RNA isolation was performed using Qiagen RNeasy kit (Qiagen, Hilden, Germany) and genomic DNA was eliminated by DNase (Qiagen, Hilden, Germany) digestion on columns following procedures indicated by the provider. RT-qPCR was performed as previously described^81^. cDNA was generated using High-Capacity cDNA Reverse Transcription kit (Applied Biosystems, Foster City, CA, 4368814) according to manual instruction. cDNA produced from 500 ng of starting RNA was diluted, and 40 ng was used to perform qPCR using Light Cycler 480 Probes Master mix (Roche Diagnostics, Mannheim, Germany, 04707494001). The real-time PCR reaction was run on Light Cycler 480 (Roche Diagnostics, Mannheim, Germany) using TaqMan premade gene expression assays (Applied Biosystems, Foster City, CA) and using the following human probes: HDAC9 (FAM-MGB)-Hs01081558_m1 and GAPDH Hs02786624_g1. Quantitative qRT-PCR was performed and the ΔΔCt method was used to calculate relative gene expression. ΔCt values were calculated as the difference between Ct values from the target gene and the housekeeping gene GAPDH.

### Adenine base editor protein expression and purification

A protein expression vector encoding ABE8e-eVRQR and an N-terminal His_8_-tag (plasmid ID LLH551) was cloned for protein production based on a previous ABE expression plasmid^17^. The ABE was overexpressed in *E. coli* BL21(DE3) cells (NEB) in Terrific Broth medium for 24 hours at 18°C following induction with 0.8% (w/v) l-rhamnose as previously described^17^. The ABE was purified using a modified protocol based on previous studies^82^. Briefly, cells were harvested by centrifugation at 6,000 g for 15 minutes and lysed by sonication in 30 mM HEPES pH 8.0, 1 M KCl, 10 mM imidazole, 10% glycerol, 2 mM TCEP, and 0.1 mg/ml lysozyme. The lysate was cleared by ultracentrifugation at 142,000 g for 45 min and purified by affinity chromatography using an EconoFit Nuvia Ni-charged IMAC column (Bio-Rad). ABE8e was eluted with a linear gradient to 500 mM imidazole and buffer exchanged to 30 mM HEPES pH 8.0, 100 mM KCl, 10% glycerol, and 2 mM TCEP using a HiPrep 26/10 Desalting column (Cytiva). Following purification by cation exchange chromatography using HiTrap SP resin (Cytiva) and elution in a linear gradient to 1.2 M KCl, ABE8e was applied to a Superose 6 Increase 10/300 GL column (Cytiva) for size exclusion chromatography and eluted in 20 mM Tris-HCl pH 7.5, 400 mM KCl, 10% glycerol and 2 mM TCEP. Purified ABE8e was concentrated to 17.7 mg/ml, resulting in a yield of 0.8 mg protein per liter of culture. Protein purity was evaluated by SDS-PAGE analysis and the final size exclusion chromatography trace and SDS-PAGE gel are presented in **Sup. Fig. 8.**

### Off-target analysis via GUIDE-seq2

GUIDE-seq2 (Lazzarotto & Li et al*, in preparation*) is an adapted version of the original GUIDE-seq method^52,53^. Briefly, approximately 20,000 HEK 293T cells were seeded per well in 96-well plates ∼20 hours prior to transfection, performed using 29 ng of nuclease expression plasmid, 12.5 ng of gRNA expression plasmid, 1 pmol of the GUIDE-seq double-stranded oligodeoxynucleotide tag (dsODN; oSQT685/686)^52,53^, and 0.3 µL of TransIT-X2 (Mirus). Genomic DNA was extracted ∼72 hours post transfection using the DNAdvance Kit (Beckman Coulter) according to manufacturer’s instructions, and then quantified by Qubit (Thermo Fisher). On-target dsODN integration was assessed by PCR amplification, library preparation, and next-generation sequencing as described above, with data analysis via CRISPREsso2^80^ run in non-pooled mode by supplying the target site spacer, the reference amplicon, and both the forward and reverse dsODN-containing amplicons as ‘HDR’ alleles with custom parameters: -w 25 -g GUIDE --plot_window_size 50. The fraction of alleles bearing an integrated dsODN was calculated as the number of reads mapped to the forward dsODN amplicon plus the number of reads mapped to the reverse dsODN amplicon divided by the sum of the total reads mapped to all three amplicons.

GUIDE-seq2 reactions were performed essentially as described (Lazzarotto & Li et al*, in preparation*) with minor modifications. Briefly, the Tn5 transposase was prepared by combining 36 µL hyperactive Tn5 (1.85 mg/mL, purified as previously described^94^), 15 µL annealed i5 adapter oligos encoding 8 nucleotide (nt) barcodes and 10-nt unique molecular indexes (UMIs) (**Supplementary Table 4**), with 52 µL 2x Tn5 dialysis buffer (100 mM HEPES-KOH pH 7.2, 200 mM NaCl, 0.2 mM EDTA, 2 mM DTT, 0.2% Triton X-100, and 20% glycerol) for 60 minutes at 24 °C. Tagmentation reactions were performed in 40 µL reactions for 7 minutes at 55 °C, containing approximately 250 ng of genomic DNA, 8 µL of the assembled Tn5/i5 -transposome, and 8 µL of freshly prepared 5x TAPS-DMF buffer (50 mM TAPS-NaOH, 25 mM MgCl_2_, and 50% dimethylformamide (DMF)). Tagmentation reactions were halted using 5 µL of a 50% proteinase K (NEB) solution (mixed with H_2_O) with incubation at 55 °C for 15 minutes, purified using SPRI-guanidine magnetic beads, and analyzed via TapeStation with High Sensitivity D5000 tapes (Agilent). Separate PCR reactions were performed using dsODN sense-and antisense-specific primers (**Supplementary Table 4**) using Platinum Taq (Thermo Fisher), with a thermocycler program of 95 °C for 5 minutes, followed by 15 cycles of temperature cycling (95 °C for 30 s, 70 °C (-1 °C per cycle) for 120 s, and 72 °C for 30 s), 20 constant cycles (95 °C for 30 s, 55 °C for 60 s, and 72 °C for 30 s), an a final extension at 72 °C for 5 minutes. PCR products were purified using SPRI beads and analyzed via QIAxcel (Qiagen) prior to sample pooling to form single sense-and antisense-libraries. Libraries were purified using the Pippin Prep (Sage Science) DNA size selection system to achieve a size range of 250-500 base pairs. Sense-and antisense-libraries were quantified using Qubit (Thermo Fisher) and pooled in equal amounts to achieve a final concentration of 2 nM. The library was sequenced using NextSeq1000/2000 P3 kit (Illumina) with cycle settings of 146, 8, 18, 146. Demultiplexed sequencing reads were down sampled to ensure equal numbers of reads for samples being compared using the same gRNA. Data analysis was performed using an updated version of the open-source GUIDE-seq2 analysis software^95^ (https://github.com/tsailabSJ/guideseq/tree/V2) with the max_mismatches parameter set to 6 (a summary of GUIDE-seq2 data is available in **Supplementary Table 1**).

### Off-target analysis via CHANGE-seq-BE

The circularization for high-throughput analysis of nuclease genome-wide effects by sequencing for base editors (CHANGE-seq-BE) method^54^ was adapted from the original CHANGE-seq protocol^55^ to be applicable for profiling BEs instead of nucleases. We performed CHANGE-seq-BE essentially as previously described^54^. Briefly, library preparation was performed using gDNA extracted from untreated cells of three *ACTA2* R179H MSMDS-fibroblast lines and liver from a mouse model of MSMDS using the Gentra PureGene Tissue Kit (Qiagen). Approximately 800 ng of purified gDNA per CHANGE-seq reaction was tagmented with a custom Tn5-transposome^27^ to an average length of 500 bp, gap repaired with KAPA HiFi HotStart Uracil+ Ready Mix (Roche) and Taq DNA ligase (NEB), and treated with a mixture of USER enzyme and T4 polynucleotide kinase (NEB). Intramolecular circularization was performed at a concentration of 5 ng/µL with T4 DNA ligase (NEB), and treated with a cocktail of exonucleases, Lambda exonuclease (NEB), Exonuclease I (NEB), Exonuclease III (NEB) and Plasmid-Safe ATP-dependent DNase (Lucigen) to enzymatically degrade remaining linear DNA molecules, followed by Quick CIP (NEB) dephosphorylation. gRNAs (Synthego) A8 and A4 for mouse and human sequences (**Supplementary Table 4**) were re-folded prior complexation with ABE8e-WT and ABE8e-eVRQR at a BE:gRNA ratio of 1:3 to ensure ribonucleoprotein complexation. *In vitro* cleavage reactions were performed with 125 ng of exonuclease-treated circularized DNA in 1x Digenome-seq deamination buffer^83^ with 300 nM ABE8e-WT or ABE8e-eVRQR protein, and 900 nM of synthetic gRNA in a 50 μL reaction. Deaminated DNA products were treated with proteinase K (NEB), followed by Endonuclease V (NEB) treatment. The Endonuclease V-treated products were end repaired with Klenow fragment exo-(NEB), A-tailed (NEB), ligated with a hairpin adapter (NEB), treated with USER enzyme (NEB) and amplified by PCR using KAPA HiFi HotStart Uracil+ Ready Mix (Roche). Completed libraries were quantified by qPCR using KAPA Library Quantification kit (Complete kit; Universal) (Roche) and sequenced with 151-bp paired-end reads on an Illumina NextSeq 2000 instrument. Data analysis was conducted using open-source CHANGE-seq-BE analysis software (https://github.com/tsailabSJ/changeseq/tree/dev) and summary data is available in **Supplementary Table 1**.

### High-throughput PAM Determination assay (HT-PAMDA)

The high-throughput PAM determination assay (HT-PAMDA) was performed using linearized randomized PAM-containing plasmid substrates that were subject to *in vitro* cleavage reactions with SpCas9-VRQR nuclease variants essentially as previously described^18,37^, with the exception that non-target controls substrates were added to the pooled PAM libraries. To normalize depletion of PAM-encoding substrates over time in the HT-PAMDA assay against uncleavable molecules (instead of utilizing the least cleaved PAMs for normalization during data analysis as described in the original method^18,37^), we cloned four control target plasmids encoding spacer sequences that do not match the gRNA and combined them with the conventional HT-PAMDA library of substrates. The four control plasmids were cloned via ligation of annealed oligonucleotides (**Supplementary Table 4**) into p11-lacY-wtx1 (Addgene ID 69056) digested with EcoRI-HF, SpeI-HF and SphI-HF (NEB). The HT-PAMDA substrate libraries containing the four control plasmids (combined at a ratio a 1:100 control plasmid pool to HT-PAMDA library) were linearized with PvuI-HF (NEB) before performing HT-PAMDA previously described^18,37^.

Sequencing reads were analyzed using custom Python scripts to determine cleavage rates on each substrate with unique spacers and PAMs similar to as previously described^18,37^, but using modified scripts to enable normalization to the control substrates (available at: https://github.com/RachelSilverstein/HT-PAMDA-2.

### Animal care and generation of the knock-in *Acta2^R179H^* mouse model

All mice were cared for under strict compliance with the Mass General Brigham Institutional Animal Care and Use Committee (IACUC), regulated by the United States Department of Agriculture and the United States Public Health Service and Institutional Animal Care and Use Committee of Massachusetts General Hospital, MA, USA (under IACUC protocols 2012N000196 and 2018N000067). The *Acta2^R179Hfl/+^* mice were generated by editing embryonic stem (ES) cells via homology directed repair (HDR) followed by injection of clonal lines into C57BL/6 albino embryos and implantation in CD-1 pseudo-pregnant females (Cyagen). Homozygous *Acta2^R179Hfl/+^* mice were crossed with Myh11-cre (all smooth muscle) for activation of the mutation. The B6.Cg-Tg(Myh11-cre,-EGFP)2Mik/J (cat# 007742) mice were purchased from Jackson laboratories. Both mouse lines were crossed to induce the expression of the *Acta2* H179 mutant protein. Mice were identified by genotyping the toe tissue sample collected by toe clipping for the purpose of identification at around 8-10 days of age. Forward Primer: 5’-CTCATGTAGGAGGGATCTAGGGA-3’ and Reverse Primer: 5’-CTCCATGGTTTTATGCAATTTGGG-3’ were used which upon PCR amplification results in different band sizes as wild-type amplicon: 201 bp, and targeted amplicon: 251 bp. Safety data, such as body weight, food consumption and clinical signs, were also collected during the study.

### AAV production

Plasmids encoding ABE8e-VRQR split into N-term and C-terminal fragments via an Npu intein and gRNA A4 for packaging into AAV9 and AAV-PR vectors were cloned as previously described^22,60,84^ (see **Supplementary Table 3** for plasmids). An AAV-PR rep/cap plasmid was used for production (Addgene plasmid #197565). AAV9 or AAV-PR vectors encoding ABEs and gRNAs were produced by UMass Chan Medical School Viral Vector Core

### Extraction of gDNA from mouse tissues

Genomic DNA was extracted from mouse tissues using the Agencourt DNAdvance protocol (Beckman Coulter). Briefly, ∼10 to 20 mg frozen tissue samples were incubated at 37 °C for 30 min prior to treatment. Lysis reactions were performed using LBH lysis buffer, 1 M DTT, and proteinase K (40mg/mL) in 200 μL reactions, incubated overnight (18 to 20 hr) at 55 °C with shaking at 100 RPM. gDNA was purified from lysate using Bind BBE solution containing magnetic beads and performing three washes with 70% ethanol. DNA was eluted in 200 μL of Elution buffer EBA, and the approximate concentrations of gDNA were quantified by Nanodrop.

### Quantification of AAV genomes in mouse tissues

Droplet Digital PCR (ddPCR) reactions were performed to quantify the approximate number of AAV genomes in transduced tissues using 20 to 60 ng of gDNA, 250 nM of each primer and 900 nM probe (**Supplementary Table 4**), and ddPCR supermix for probes (no dUTP) (BioRad) in 20 μL reactions. Droplets were generated using a QX200 Automated Droplet Generator (BioRad). Thermal cycling conditions were: 1 cycle of (95 °C for 10 min), 40 cycles of (94 °C for 30 sec, 58 °C for 1 min), 1 cycle of (98 °C for 10 min), hold at 4 °C. PCR products were analyzed using a QX200 Droplet Reader (BioRad) and absolute quantification of inserts was determined using QuantaSoft (v1.7.4).

### Ultrasounds

Nair hair removal cream was used on all mice the day prior to ultrasounds. All ultrasounds were performed on awake, unsedated mice using the Visualsonics Vevo660 imaging system and a 30 MHz transducer. The aorta was imaged and analysed by MicroDicom software using a standard parasternal long axis view. Dimensions from each animal represent averages of measurements made on still frames in systole of the maximal internal diameter of the aortic valve annulus, aortic sinuses, sinotubular junction, or ascending aorta by a researcher blinded to genotype.

### Internal carotid artery measurements

To measure the diameter the internal carotid arteries (ICAs), mice were anesthesia with isoflurane and transcardially perfused with black india ink (Higgins, Chartpak, Inc. MA, USA). Brains were collected and photographed. The diameter of the ICAs in both left and right hemispheres was measured using ImageJ (NIH, Bethesda, MD, USA), and calculated the average of both sides.

### Rotarod and Open Field Test

Motor coordination was assessed using Ugo Basile Rota-Rod for mice (Model# 47600). Mice were placed on the beam of a rotarod revolving at a 4 RPM. default speed, facing in the opposite orientation to rotation. The speed was gradually accelerated to a maximum of 40 RPM over a 5 minute test session. The latency before falling was measured up to a maximum total time on the rod of 5 min for 3 trials. Experiments were performed over two days: the first day mice were trained for the experimental procedure, and on the second consecutive day the test was performed.

The open field task is a sensorimotor test used to determine general activity levels, gross locomotor activity, and exploration habits in mice. Mice were placed at the center of a square, white plexiglass box arena and allowed to freely move for 5 minutes while being recorded by an overhead camera. The footage is then analyzed by activity monitor automated tracking system software for total distance travelled.

### Tissue Isolation and Histological Analysis

For aortas and neurovascular circulation gross organ images, latex was injected into the left ventricular apex under low pressure until it was visible in the femoral artery. Animals were then fixed in Formalin (10%) for 24 h before transfer to 70% ethanol for dissection, photography, and storage. Other organs were then resected.

Mouse aortic SMCs were isolated by standard explant of the ascending section of the aortas from Acta2R179Hfl/+ or Myh11-Cre: Acta2R179Hfl/+ mice. To preserve cell identity, all experiments were carried out at passages 1–5. Human and murine SMCs were grown with SMC growth medium from Cell Applications Inc. (catalog 311-500).

For histological analysis, organs were removed from the animals or dissected in situ for photography prior to paraffinization and sectioning (7 μM). For DAB antibody labelling, the tissue sections were initially deparaffinized in xylene for two cycles of 6 minutes each, followed by hydration in a series of ethanol washes, including 100% ethanol for 6 minutes, and treatment with 0.5% H2O2 in methanol for 10 minutes. Further hydration was achieved with 90% ethanol for 3 minutes and 70% ethanol for 3 minutes, followed by washing with phosphate-buffered saline (PBS) three times for 3 minutes each. Antigen retrieval was performed either by incubating sections in a water bath with sodium citrate for 30 minutes or with EDTA for 15 minutes, depending on the primary antibody used. After cooling the sections on the benchtop for 20 minutes, they were washed in PBS three times for 3 minutes each. Nonspecific binding was blocked by incubating sections in PBS containing 3% bovine serum albumin (BSA) for 1 hour at room temperature, followed by overnight incubation with the primary antibodies anti-smooth muscle actin (SMA, ab5694, 1:1000; abcam) or anti-myelin basic protein (MBP, ab209328, 1:1000; abcam) diluted in 1% BSA/PBS at 4 °C.

On the second day, sections were washed in PBS three times for 3 minutes each before being incubated with DAKO Rabbit/Mouse HRP Kit-provided HRP (Mouse-K4001, Rabbit-K4003; Dako) for 30 minutes at room temperature. After another wash with PBS, the signal was developed using the DAKO DAB Kit (K3468; Dako) for 2 minutes at room temperature. The reaction was terminated with water, followed by counterstaining using Hematoxylin (7231; Epredia). Finally, sections underwent three washes with water for dehydration, were mounted with xylene, and cover slipped for examination.

For DAB staining visualization, the stained slices were imaged using a Zeiss LSM 800 microscope, which provided high-resolution images necessary for detailed analysis.

For the immunolabel intensity calculation, the mean immune labelling intensity for MBP and SMA labeled brain slices was calculated using ImageJ/FIJI software (V.1.54f). The region of interest (ROI) was selected, and the mean pixel intensity within these regions was measured for quantitative comparison of immunolabeling across samples.

The extent of cortical myelination was evaluated by measuring the proportion of myelinated cortex in brain slices. The myelinated cortex area and the total cortex area were measured using ImageJ/FIJI. The proportion of myelinated cortex was calculated using the following formula: Proportion of Myelinated Cortex = (Total Cortex Area\Myelinated Cortex Area) × 100%.

The tortuosity index of blood vessels in the brain slices was calculated to assess vascular changes. Images were processed using NIH’s ImageJ software. The Free Hand tool in ImageJ was used to outline the perimeter of both the outer and inner membranes of the vessels. The tortuosity index was determined by the ratio of the inner perimeter to the outer perimeter: Tortuosity Index = Outer Perimeter \ Inner Perimeter.

## Data availability

Primary datasets are available in **Supplementary Tables 1,5,6,7**. Sequencing datasets will be deposited with the NCBI Sequence Read Archive (SRA) under **PRJNAxxxxxx**. Plasmids from this study will be made available through Addgene.

## Acknowledgements

This work was supported by a Charles A. King Trust Postdoctoral Research Fellowship, Bank of America, N.A., Co-Trustees (C.R.R.A.), a James L. and Elisabeth C. Gamble Endowed Fund for Neuroscience Research / Mass General Neuroscience Transformative Scholar Award (C.R.R.A.), a MGH Physician/Scientist Development Award (C.R.R.A.), a Natural Sciences and Engineering Research Council of Canada (NSERC) Postgraduate Scholarship-Doctoral Postgraduate Scholarship-Doctoral (PGS D – 567791 to R.A.S.), an EMBO Long Term Fellowship (ALTF 750-2022; to J.F.daS), a Swiss National Science Foundation grant (P180777; to F.M.C.B.), St. Jude Children’s Research Hospital and ALSAC and National Institutes of Allergy and Infectious Diseases awards U01AI176470 and U01AI176471 (S.Q.T.), an MGH Howard M. Goodman Fellowship (to B.P.K.), the Kayden-Lambert MGH Research Scholar Award 2023-2028 (B.P.K.), a sponsored research agreement with Angea Biotherapeutics (R.M., D.Y.C., C.A.M., M.E.L., B.P.K., and P.L.M.), and National Institutes of Health (NIH) grants K01NS134784 (C.R.R.A.), R01HL162928 (R.M.), K08NS112601 (D.Y.C), R35GM142553 (L.H.C.), DC017117 (C.A.M.), DP2CA281401 (B.P.K), P01HL142494 (B.P.K.), and R01NS125353 (to P.L.M, M.E.L., & B.P.K.).

## Author contributions

M.E.L., B.P.K., and P.L.M. conceived of and designed the study. All authors designed, performed, or supervised experiments, and/or analyzed data. C.R.R.A., C.L.L.C., L.L.H., H.E.S., P.K., L.R.F., and M.S. performed cell culture experiments. C.R.R.A., S.D., V.K., C.L.L.C., L.L.H., H.E.S., C.E.S., P.K., L.R.F., C.E.F., M.S., and S.Y. performed molecular and biochemical experiments. S.D., V.K., C.L.L.C., C.E.S., C.E.F., T.I., J.L., R.R., and D.Y.C. conducted the *in vivo* experiments with mice and histological analysis. C.R.R.A., L.L.H., H.E.S., L.R.F., S.Y., and J.F.S. performed plasmid cloning and lentivirus production. N.K. and R.A.S. performed HT-PAMDA experiments. D.L.C and C.A.M. performed titrations of AAV preparations and advised on AAV-related experiments. L.H.C. and F.M.C.B. performed protein purification. S.Q.T. and R.K.W. designed and performed CHANGE-seq-BE experiments. C.R.R.A., M.E.L. and B.P.K. wrote the manuscript with contributions or revisions from all authors.

## References

1. Cw, T. et al. Heart Disease and Stroke Statistics-2023 Update: A Report From the American Heart Association. Circulation 147, (2023).

2. Ostrem, B. E. L., Godfrey, D., Caruso, P. A. & Musolino, P. L. Monogenic Causes of Cerebrovascular Disease in Childhood: A Case Series. Pediatr. Neurol. 149, 39–43 (2023).

3. Jankovic, M. et al. The Genetic Basis of Strokes in Pediatric Populations and Insight into New Therapeutic Options. Int. J. Mol. Sci. 23, 1601 (2022).

4. Grossi, A. et al. Targeted re-sequencing in pediatric and perinatal stroke. Eur. J. Med. Genet. 63, 104030 (2020).

5. Ilinca, A. et al. A stroke gene panel for whole-exome sequencing. Eur. J. Hum. Genet. 27, 317–324 (2019).

6. Munot, P. et al. A novel distinctive cerebrovascular phenotype is associated with heterozygous Arg179 ACTA2 mutations. Brain J. Neurol. 135, 2506–2514 (2012).

7. Milewicz, D. M. et al. De novo ACTA2 mutation causes a novel syndrome of multisystemic smooth muscle dysfunction. Am. J. Med. Genet. A. 152A, 2437–2443 (2010).

8. Richer, J. et al. R179H mutation in ACTA2 expanding the phenotype to include prune-belly sequence and skin manifestations. Am. J. Med. Genet. A. 158A, 664–668 (2012).

9. Regalado, E. S. et al. Clinical history and management recommendations of the smooth muscle dysfunction syndrome due to ACTA2 arginine 179 alterations. Genet. Med. Off. J. Am. Coll. Med. Genet. 20, 1206–1215 (2018).

10. Lauer, A. et al. Cerebrovascular Disease Progression in Patients With ACTA2 Arg179 Pathogenic Variants. Neurology 96, e538–e552 (2021).

11. Gaudelli, N. M. et al. Programmable base editing of A•T to G•C in genomic DNA without DNA cleavage. Nature 551, 464–471 (2017).

12. Komor, A. C., Kim, Y. B., Packer, M. S., Zuris, J. A. & Liu, D. R. Programmable editing of a target base in genomic DNA without double-stranded DNA cleavage. Nature 533, 420–424 (2016).

13. Nishida, K. et al. Targeted nucleotide editing using hybrid prokaryotic and vertebrate adaptive immune systems. Science 353, (2016).

14. Rees, H. A. & Liu, D. R. Base editing: precision chemistry on the genome and transcriptome of living cells. Nat. Rev. Genet. 19, 770–788 (2018).

15. Gaudelli, N. M., et al. Directed evolution of adenine base editors with increased activity and therapeutic application. Nat. Biotechnol. 38, 892–900 (2020).

16. Richter, M. F., et al. Phage-assisted evolution of an adenine base editor with improved Cas domain compatibility and activity. Nat. Biotechnol. 38, 883–891 (2020).

17. Huang, T. P., Newby, G. A. & Liu, D. R. Precision genome editing using cytosine and adenine base editors in mammalian cells. Nat. Protoc. 16, 1089–1128 (2021).

18. Walton, R. T., Christie, K. A., Whittaker, M. N. & Kleinstiver, B. P. Unconstrained genome targeting with near-PAMless engineered CRISPR-Cas9 variants. Science 368, 290–296 (2020).

19. Miller, S. M. et al. Continuous evolution of SpCas9 variants compatible with non-G PAMs. Nat. Biotechnol. 38, 471–481 (2020).

20. Nishimasu, H. et al. Engineered CRISPR-Cas9 nuclease with expanded targeting space. Science 361, 1259–1262 (2018).

21. Kleinstiver, B. P. et al. Engineered CRISPR-Cas9 nucleases with altered PAM specificities. Nature 523, 481–485 (2015).

22. Alves, C. R. R., et al. Optimization of base editors for the functional correction of SMN2 as a treatment for spinal muscular atrophy. Nat. Biomed. Eng. 8, 118–131 (2024).

23. Bzhilyanskaya, V. et al. High-fidelity PAMless base editing of hematopoietic stem cells to treat chronic granulomatous disease. Sci. Transl. Med. 16, eadj6779 (2024).

24. Anzalone, A. V., Koblan, L. W. & Liu, D. R. Genome editing with CRISPR-Cas nucleases, base editors, transposases and prime editors. Nat. Biotechnol. 38, 824–844 (2020).

25. Porto, E. M., Komor, A. C., Slaymaker, I. M. & Yeo, G. W. Base editing: advances and therapeutic opportunities. Nat. Rev. Drug Discov. 19, 839–859 (2020).

26. Porto, E. M. & Komor, A. C. In the business of base editors: Evolution from bench to bedside. PLOS Biol. 21, e3002071 (2023).

27. Christie, K. A. & Kleinstiver, B. P. Making the cut with PAMless CRISPR-Cas enzymes. Trends Genet. 37, 1053–1055 (2021).

28. Chatterjee, P., Jakimo, N. & Jacobson, J. M. Minimal PAM specificity of a highly similar SpCas9 ortholog. Sci. Adv. 4, eaau0766 (2018).

29. Zhao, L. et al. PAM-flexible genome editing with an engineered chimeric Cas9. Nat. Commun. 14, 6175 (2023).

30. Chai, A. C. et al. Base editing correction of hypertrophic cardiomyopathy in human cardiomyocytes and humanized mice. Nat. Med. 1–11 (2023) doi:10.1038/s41591-022-02176-5.

31. Lebek, S. et al. Ablation of CaMKIIδ oxidation by CRISPR-Cas9 base editing as a therapy for cardiac disease. Science 379, 179–185 (2023).

32. Hibshman, G. N. et al. Unraveling the mechanisms of PAMless DNA interrogation by SpRY-Cas9. Nat. Commun. 15, 3663 (2024).

33. Anzalone, A. V. et al. Search-and-replace genome editing without double-strand breaks or donor DNA. Nature 576, 149–157 (2019).

34. Kleinstiver, B. P. et al. High-fidelity CRISPR-Cas9 nucleases with no detectable genome-wide off-target effects. Nature 529, 490–495 (2016).

35. Spencer, J. M. & Zhang, X. Deep mutational scanning of S. pyogenes Cas9 reveals important functional domains. Sci. Rep. 7, 16836 (2017).

36. Anders, C., Bargsten, K. & Jinek, M. Structural Plasticity of PAM Recognition by Engineered Variants of the RNA-Guided Endonuclease Cas9. Mol. Cell 61, 895–902 (2016).

37. Walton, R. T., Hsu, J. Y., Joung, J. K. & Kleinstiver, B. P. Scalable characterization of the PAM requirements of CRISPR-Cas enzymes using HT-PAMDA. Nat. Protoc. 16, 1511–1547 (2021).

38. Koblan, L. W., et al. Improving cytidine and adenine base editors by expression optimization and ancestral reconstruction. Nat. Biotechnol. 36, 843–846 (2018).

39. Kleinstiver, B. P. et al. High-fidelity CRISPR-Cas9 nucleases with no detectable genome-wide off-target effects. Nature 529, 490–495 (2016).

40. Clement, K. et al. CRISPResso2 provides accurate and rapid genome editing sequence analysis. Nat. Biotechnol. 37, 224–226 (2019).

41. Kim, H. S., Jeong, Y. K., Hur, J. K., Kim, J.-S. & Bae, S. Adenine base editors catalyze cytosine conversions in human cells. Nat. Biotechnol. 37, 1145–1148 (2019).

42. Jeong, Y. K. et al. Adenine base editor engineering reduces editing of bystander cytosines. Nat. Biotechnol. 1–8 (2021) doi:10.1038/s41587-021-00943-2.

43. Arbab, M. et al. Determinants of Base Editing Outcomes from Target Library Analysis and Machine Learning. Cell 182, 463–480.e30 (2020).

44. Kim, N., et al. Deep learning models to predict the editing efficiencies and outcomes of diverse base editors. Nat. Biotechnol. 42, 484–497 (2024).

45. Lino Cardenas, C. L., et al. An HDAC9-MALAT1-BRG1 complex mediates smooth muscle dysfunction in thoracic aortic aneurysm. Nat. Commun. 9, 1009 (2018).

46. Malhotra, R. et al. HDAC9 is implicated in atherosclerotic aortic calcification and affects vascular smooth muscle cell phenotype. Nat. Genet. 51, 1580–1587 (2019).

47. Lino Cardenas, C. L., et al. HDAC9 complex inhibition improves smooth muscle-dependent stenotic vascular disease. JCI Insight 4, e124706 (2019).

48. Chou, E. L. et al. Aortic Cellular Diversity and Quantitative Genome-Wide Association Study Trait Prioritization Through Single-Nuclear RNA Sequencing of the Aneurysmal Human Aorta. Arterioscler. Thromb. Vasc. Biol. 42, 1355–1374 (2022).

49. Bae, S., Park, J. & Kim, J.-S. Cas-OFFinder: a fast and versatile algorithm that searches for potential off-target sites of Cas9 RNA-guided endonucleases. Bioinformatics 30, 1473–1475 (2014).

50. Cas-OFFinder: a fast and versatile algorithm that searches for potential off-target sites of Cas9 RNA-guided endonucleases -PubMed. https://pubmed.ncbi.nlm.nih.gov/24463181/.

51. Lazzarotto, C. et al. CHANGE-seq-BE enables simultaneously sensitive and unbiased in vitro profiling of base editor genome-wide activity. 2024.03.28.586621 Preprint at 10.1101/2024.03.28.586621 (2024).

52. Tsai, S. Q. et al. GUIDE-seq enables genome-wide profiling of off-target cleavage by CRISPR-Cas nucleases. Nat Biotechnol 33, 187–197 (2015).

53. Malinin, N. L. et al. Defining genome-wide CRISPR-Cas genome-editing nuclease activity with GUIDE-seq. Nat. Protoc. 16, 5592–5615 (2021).

54. Lazzarotto, C. R. et al. CHANGE-seq-BE enables simultaneously sensitive and unbiased in vitro profiling of base editor genome-wide activity. 2024.03.28.586621 Preprint at 10.1101/2024.03.28.586621 (2024).

55. Lazzarotto, C. R. et al. CHANGE-seq reveals genetic and epigenetic effects on CRISPR-Cas9 genome-wide activity. Nat. Biotechnol. 38, 1317–1327 (2020).

56. Xin, H.-B., Deng, K.-Y., Rishniw, M., Ji, G. & Kotlikoff, M. I. Smooth muscle expression of Cre recombinase and eGFP in transgenic mice. Physiol. Genomics 10, 211–215 (2002).

57. Liu, Z. et al. Vascular disease-causing mutation, smooth muscle α-actin R258C, dominantly suppresses functions of α-actin in human patient fibroblasts. Proc. Natl. Acad. Sci. U. S. A. 114, E5569–E5578 (2017).

58. Koblan, L. W. et al. In vivo base editing rescues Hutchinson-Gilford progeria syndrome in mice. Nature 589, 608–614 (2021).

59. Levy, J. M. et al. Cytosine and adenine base editing of the brain, liver, retina, heart and skeletal muscle of mice via adeno-associated viruses. *Nat*. Biomed. Eng. 4, 97–110 (2020).

60. Ramirez, S. H. et al. An Engineered Adeno-Associated Virus Capsid Mediates Efficient Transduction of Pericytes and Smooth Muscle Cells of the Brain Vasculature. Hum. Gene Ther. 34, 682–696 (2023).

61. Zettler, J., Schütz, V. & Mootz, H. D. The naturally split Npu DnaE intein exhibits an extraordinarily high rate in the protein trans-splicing reaction. FEBS Lett. 583, 909–914 (2009).

62. KLEINSTIVER, B. & WALTON, R. T. Crispr-cas enzymes with enhanced on-target activity. (2021).

63. Ran, F. A. et al. In vivo genome editing using Staphylococcus aureus Cas9. Nature 520, 186–191 (2015).

64. Kleinstiver, B. P. et al. Broadening the targeting range of Staphylococcus aureus CRISPR-Cas9 by modifying PAM recognition. Nat. Biotechnol. 33, 1293–1298 (2015).

65. Kleinstiver, B. P. et al. Engineered CRISPR-Cas12a variants with increased activities and improved targeting ranges for gene, epigenetic and base editing. Nat. Biotechnol. 37, 276–282 (2019).

66. Chen, Y. et al. Synergistic engineering of CRISPR-Cas nucleases enables robust mammalian genome editing. The Innovation 3, 100264 (2022).

67. McGaw, C. et al. Engineered Cas12i2 is a versatile high-efficiency platform for therapeutic genome editing. Nat. Commun. 13, 2833 (2022).

68. Zhang, H. et al. An engineered xCas12i with high activity, high specificity, and broad PAM range. Protein Cell 14, 540–545 (2023).

69. Yan, H. et al. Assessing and engineering the IscB–ωRNA system for programmed genome editing. Nat. Chem. Biol. 1–12 (2024) doi:10.1038/s41589-024-01669-3.

70. Wu, T. et al. An engineered hypercompact CRISPR-Cas12f system with boosted gene-editing activity. Nat. Chem. Biol. 1–10 (2023) doi:10.1038/s41589-023-01380-9.

71. Xiao, Q., et al. Engineered IscB–ωRNA system with expanded target range for base editing. Nat. Chem. Biol. 1–9 (2024) doi:10.1038/s41589-024-01706-1.

72. Lin, J., et al. Adenine base editing-mediated exon skipping restores dystrophin in humanized Duchenne mouse model. Nat. Commun. 15, 5927 (2024).

73. Davis, J. R., et al. Efficient in vivo base editing via single adeno-associated viruses with size-optimized genomes encoding compact adenine base editors. Nat. Biomed. Eng. 1–12 (2022) doi:10.1038/s41551-022-00911-4.

74. Zhang, H. et al. Adenine Base Editing in Vivo with a Single Adeno-Associated Virus Vector. 2021.12.13.472434 doi:10.1101/2021.12.13.472434.

75. Liu, Z. et al. An all-in-one AAV vector for cardiac-specific gene silencing by an adenine base editor. 2024.09.30.615742 Preprint at 10.1101/2024.09.30.615742 (2024).

76. Tsuchida, C. A., Wasko, K. M., Hamilton, J. R. & Doudna, J. A. Targeted nonviral delivery of genome editors in vivo. Proc. Natl. Acad. Sci. 121, e2307796121 (2024).

77. Kim, J., Eygeris, Y., Ryals, R. C., Jozić, A. & Sahay, G. Strategies for non-viral vectors targeting organs beyond the liver. Nat. Nanotechnol. 1–20 (2023) doi:10.1038/s41565-023-01563-4.

78. Gibson, D. G., et al. Enzymatic assembly of DNA molecules up to several hundred kilobases. Nat. Methods 6, 343–345 (2009).

79. Rohland, N. & Reich, D. Cost-effective, high-throughput DNA sequencing libraries for multiplexed target capture. Genome Res. 22, 939–946 (2012).

80. Clement, K., et al. CRISPResso2 provides accurate and rapid genome editing sequence analysis. Nat. Biotechnol. 37, 224–226 (2019).

81. DeRosa, S., et al. MCOLN1 gene therapy corrects neurologic dysfunction in the mouse model of mucolipidosis IV. Hum. Mol. Genet. 30, 908–922 (2021).

82. Lapinaite, A., et al. DNA capture by a CRISPR-Cas9-guided adenine base editor. Science 369, 566–571 (2020).

83. Kim, D., Kim, D., Lee, G., Cho, S.-I. & Kim, J.-S. Genome-wide target specificity of CRISPR RNA-guided adenine base editors. Nat. Biotechnol. 37, 430–435 (2019).

84. Hanlon, K. S., et al. Selection of an Efficient AAV Vector for Robust CNS Transgene Expression. Mol. Ther. Methods Clin. Dev. 15, 320–332 (2019).

