## Supplementary Notes and Figures for "*In vivo* Treatment of a Severe Vascular Disease via a Bespoke CRISPR-Cas9 Base Editor"

### Supplementary Materials

Supplementary Tables *Attached separately*

**Supplementary Table 1**: Cas-OFFinder, GUIDE-seq2, and CHANGE-seq-BE results

**Supplementary Table 2**: gRNA target sites

**Supplementary Table 3**: Plasmids

**Supplementary Table 4**: Oligonucleotides and probes

**Supplementary Table 5**: rhAmpSeq results for gRNA A4 off-target analysis

**Supplementary Table 6**: rhAmpSeq results for gRNA A8 off-target analysis

**Supplementary Table 7**: Primary datasets

***NOTE:*** All Supplementary Tables are attached separately as .xlsx files.

Supplementary Notes

**Supplementary Note 1**: Development of HEK 293T cell lines harboring *ACTA2* R179H p.2

**Supplementary Note 2**: Discussion of the use of AAV-PR and AAV9-based ABEs to treat MSMDS p.2

Supplementary Figures and Legends

**Supplementary Figures 1-33** p.3

Supplementary References p.39

### Supplementary Notes

**Supplementary Note 1**: Development of HEK 293T cell lines harboring *ACTA2* R179H

We sought to generate HEK 293T cell lines bearing the *ACTA2* R179H mutation, owing to the ease of cell culture and transfection to then subsequently utilize the cell lines to rapidly screen and prioritize combinations of editors and guide RNAs (gRNAs) that maximize editing efficiencies. To establish cell lines encoding either homozygous and heterozygous mutations, we tested various prime editing[1] strategies (**Sup. Fig. 1a**). We designed prime editing guide RNAs (pegRNAs) near the *ACTA2 R179* locus targeting sites with NGG or NGT protospacer adjacent motifs (PAMs), and for each site we screened different primer binding site (PBS) and reverse transcriptase template (RTT) lengths (**Sup. Fig. 1b**). Transfections in HEK 293T cells were performed to investigate the efficacy of SpCas9-PE2[1] or SpG-PE2[2,3] paired with these pegRNAs and various PE3 nicking gRNAs[1] (ngRNAs) to generate the intended *ACTA2* R179H mutation, with some combinations achieving >10% precise introduction of the mutation (**Sup. Figs. 1c,d**). To create clonal cell lines, plasmids encoding the most optimal editor (SpCas9-PE2 paired with a pegRNA encoding an 11 bp PBS and 13 nt RTT and ngRNA #3; **Sup. Figs. 1b,c**) were transfected into fresh HEK 293T cells were single cell cloned (**Sup. Fig. 1e**). Targeted sequencing of the *ACTA2* locus in genomic DNA extracted from clonal cell lines confirmed that some harbored homozygous or heterozygous *ACTA2* R179H mutations. To maximize sensitivity to detect correction, a homozygous HEK 293T cell line harboring *ACTA2* R179H (hereafter named HEK 293T-*ACTA2*-R179H) was further used in this study to screen base editor strategies to correct the *ACTA2* R179H mutation.

**Supplementary Note 2**: Discussion of the use of AAV-PR and AAV9-based ABEs to treat MSMDS

In addition to the effectiveness of AAV-PR-ABE to treat MSMDS mice, we also observed that AAV9 could effectively rescue survival of mice despite having a lower tropism for smooth muscle cells (SMCs)[4]. In contrast to AAV-PR, AAV9 is not reported to efficiently transduce SMCs in brain using conventional fluorescent reporters. It has been demonstrated that fluorescent reporters only capture a portion of all AAV transduction events, which can be transient in nature, but could lead to permanent modifications in the context of gene editing approaches[5]. This indicates that low levels of transduction in SMCs may support therapeutic effects. However, AAV-PR offers potential advantages over AAV9, including that AAV-PR has higher specificity to MSMDS-related target cells and should therefore with limit off-target effects in the CNS across neurons and astrocytes (that can be transduced with AAV9). Furthermore, systemic based AAV therapies have not directly scaled from rodents to large animals and humans in terms of dose/weight, in that the latter usually requires higher doses to achieve therapeutic benefit[6–8]. Thus more efficient capsids such as AAV-PR may permit clinical translation at doses that are below thresholds (~$10^{14}$ vg/kg) that have caused toxicities such as thrombotic microangiopathy in clinical trials.

### Supplementary Figures and Legends

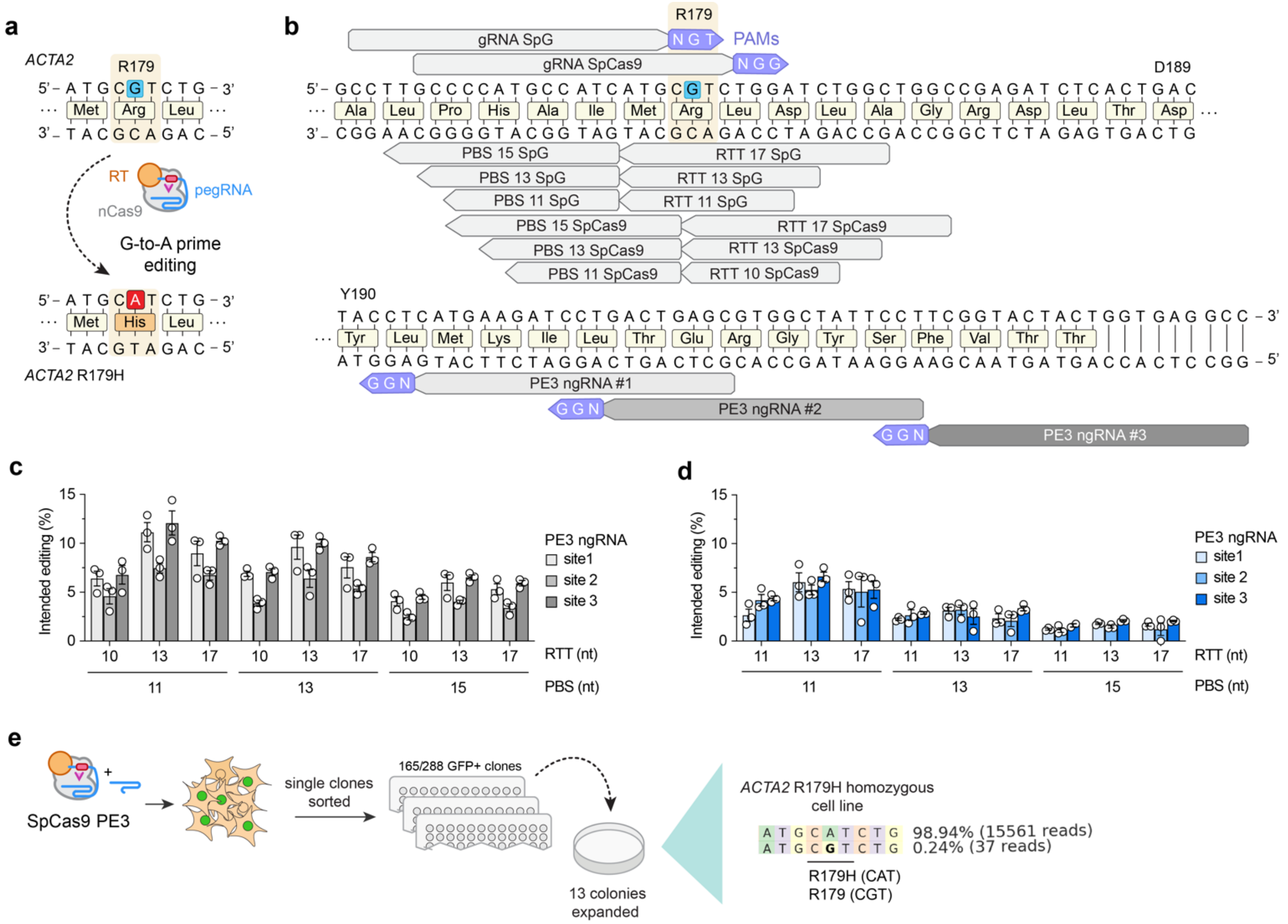


**Supplementary Figure 1. Generation of a HEK 293T cell line harboring the *ACTA2* R179H mutation. a**, Schematic of the *ACTA2 R179H* mutation to be generated via prime editing. nCas9, nickase SpCas9 H840A; RT, reverse transcriptase; pegRNA, prime editing guide RNA. **b**, Schematic of the *ACTA2* gene locus near the site of amino acid R179 with annotations for two target sites (bearing NGT and NGG PAMs), pegRNA designs with various primer binding site (PBS) lengths and reverse transcriptase template (RTT) lengths, and three additional target sites for nicking gRNAs (ngRNAs). **c**,**d**, Precise intended G-to-A editing to introduce the *ACTA2* R179H mutation using a SpCas9-PE2[1] paired with a pegRNA targeting the NGG PAM target site and various ngRNAs (**panel c**), or when using SpG-PE2[2,3] paired with a pegRNA targeting the NGT PAM target site and various ngRNAs (**panel d**). Experiments were performed with several combinations of pegRNAs bearing different length PBSs and RTTs, and three different ngRNAs. Prime editing efficiencies assessed by targeted sequencing and analyzed using CRISPResso2; mean, s.e.m., and individual datapoints shown for n = 3 independent biological replicates. **e**, Schematic of experimental approach to sort and expand clonal HEK 293T cells harboring the *ACTA2 R179H* mutation when using the most efficiency PE option from **panel c**.

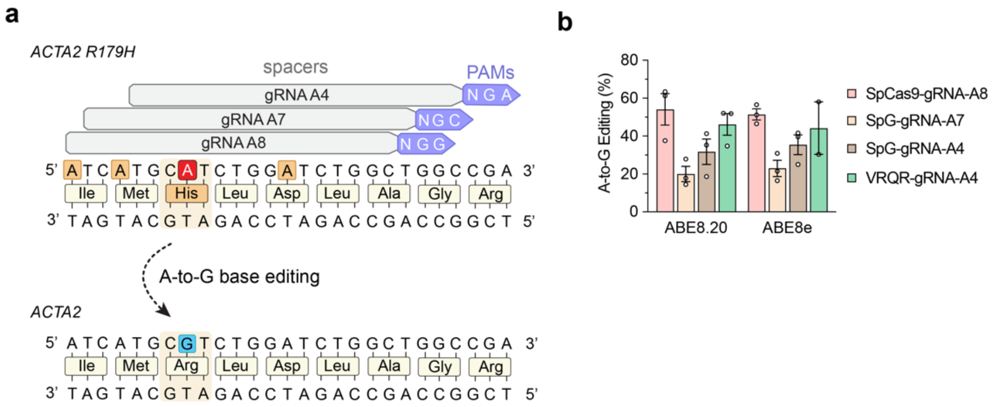


**Supplementary Figure 2. Base editing to correct *ACTA2* R179H when using gRNAs A4, A7, or A8. a**, Schematic of target site spacers and PAMs for *ACTA2* R179H correction. **b**, On-target A-to-G base editing to correct *ACTA2* R179H in homozygous HEK 293T cells when using ABEs comprised of deaminase domains ABE8.20m[9] and ABE8e[10] fused to WT SpCas9 (with gRNA A8), the SpCas9-derived PAM variant enzyme SpG[2] (with gRNAs A4 or A7), or SpCas9-VRQR[11,12] (with gRNA A4). Mean, s.e.m., and individual datapoints shown from experiments with n = 3 independent biological replicates.

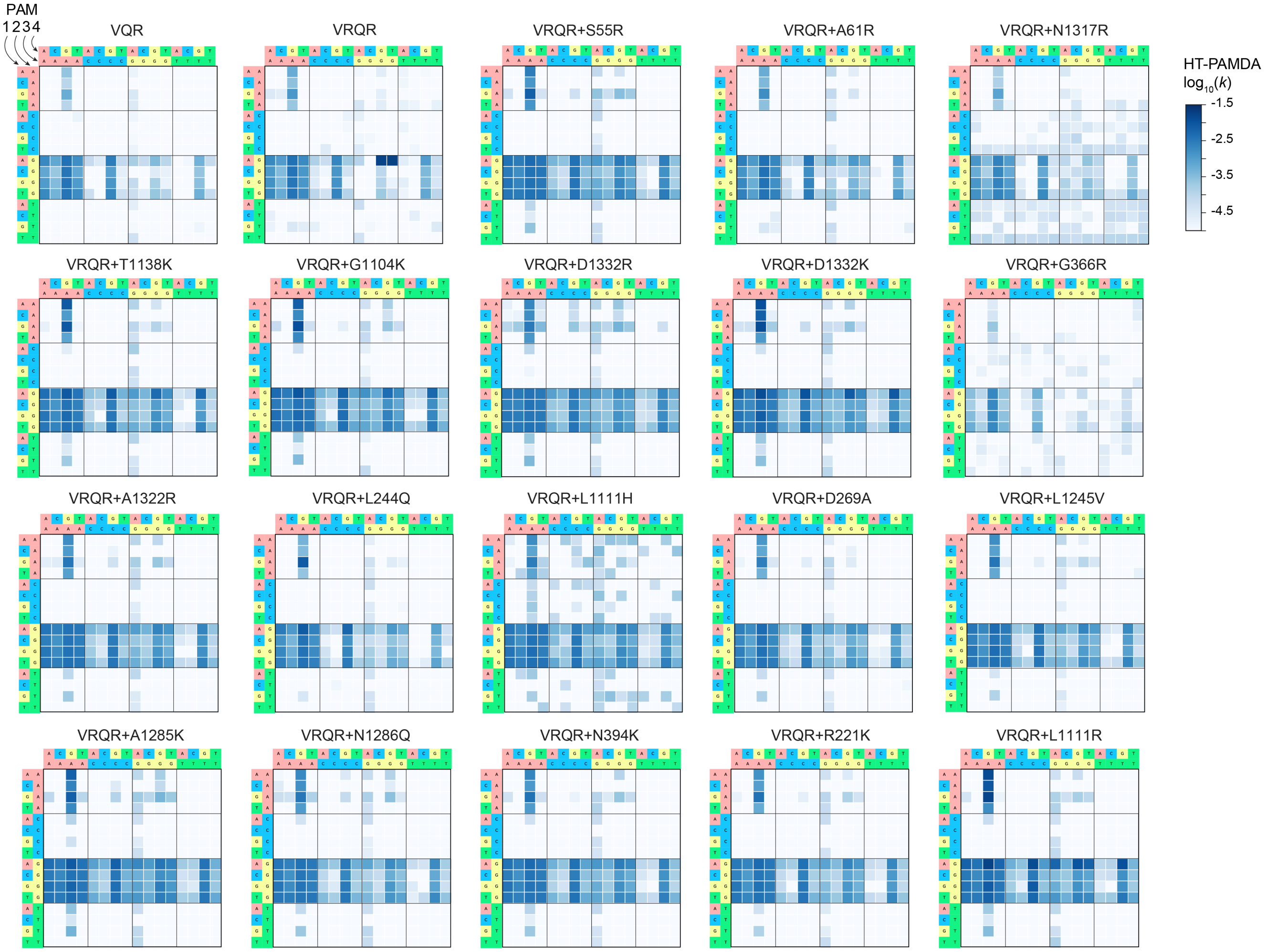


**Supplementary Figure 3. Characterization of engineered SpCas9-VRQR variant enzymes.** Heatmap representations of the PAM profiles of SpCas9-VRQR[11,12] and engineered derivatives bearing mutations developed in this study by structure guided engineering, or those described in previous studies[2,13–15]. PAM profiles were determined using the high-throughput PAM determination assay (HT-PAMDA)[2,16]. The $\log_{10}$ rate constants (*k*) are the mean of n = 2 biological replicate HT-PAMDA experiments using two distinct spacer sequences.

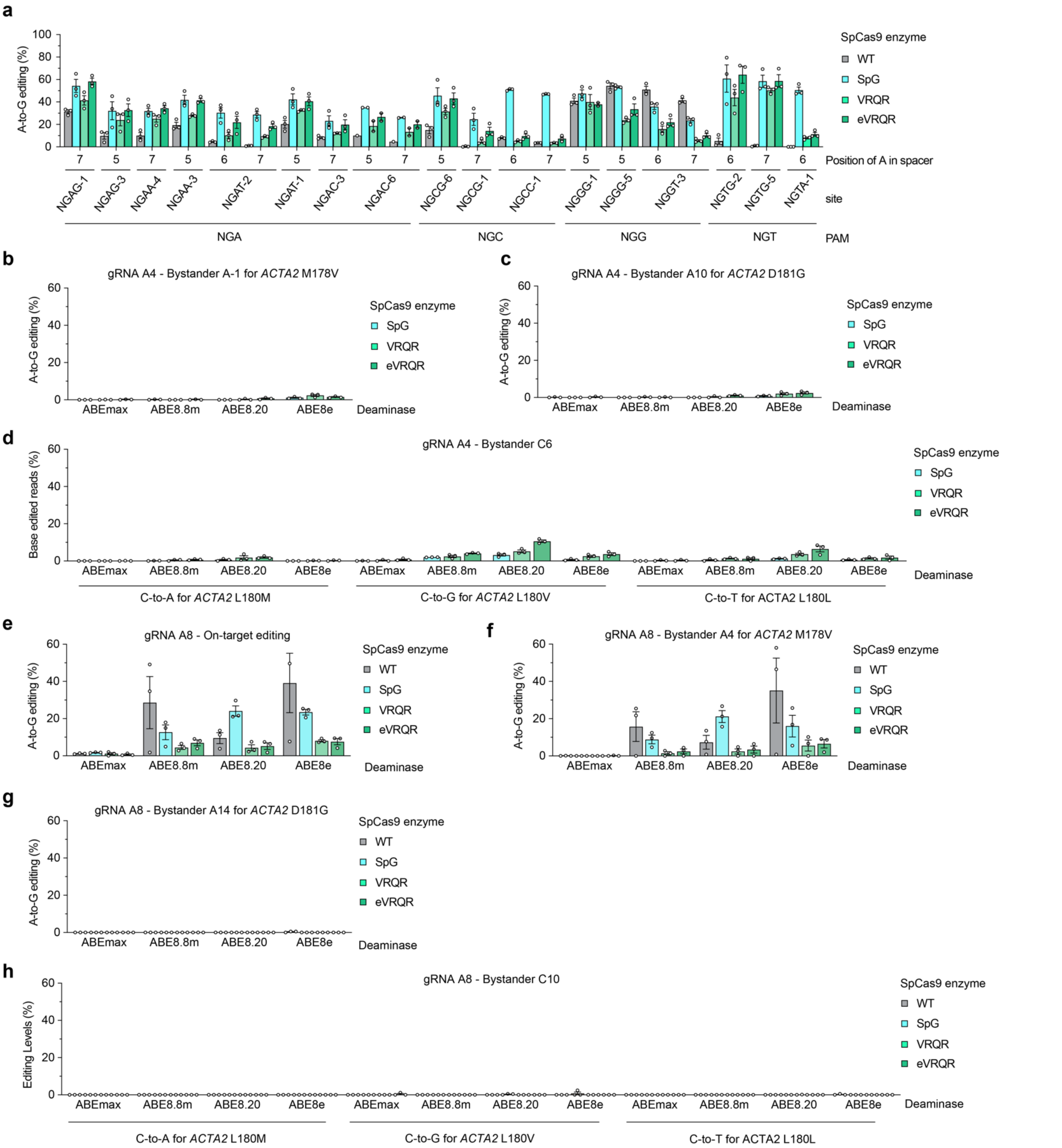


**Supplementary Figure 4. On-target and bystander base editing in experiments with SpCas9 PAM variant enzymes. a**, On-target editing across several endogenous target sites in HEK 293T cells when using various adenine base editors (ABEs) comprised of TadA8e fused to wild-type SpCas9 (WT), SpG[2], VRQR[11,12], or eVRQR. **b,c,** Bystander A-to-G editing that causes an *ACTA2* M178V via A-to-G editing at position A-1 (**panel b**) or *ACTA2* D181G via A-to-G editing at position A10 (**panel c**) when using *ACTA2* R179H gRNA A4 with various ABEs comprised of different TadA deaminase domains from ABEmax[17,18], ABE8.8m[9], ABE8.20m[9] or ABE8e[10] fused to different Cas9 enzymes including wild-type SpCas9 (WT), SpG, VRQR, or eVRQR. **d,**

Bystander C-to-T editing that causes a silent *ACTA2* L180L substitution, C-to-A editing that causes an L180M substitution, or C-to-G editing that causes an L180V substitution, at position C6 when using *ACTA2* R179H gRNA A4. On-target A-to-G editing to correct the *ACTA2 R179H* mutation using gRNA A4 is displayed in **Fig. 1f**. **e-h**, Base editing with *ACTA2* R179H gRNA A8 with various ABEs comprised of different TadA deaminase domains from ABEmax[17,18], ABE8.8m[9], ABE8.20m[9] or ABE8e[10] fused to different Cas9 enzymes including wild-type SpCas9 (WT), SpG[2], VRQR[11,12] or eVRQR, resulting in on-target A-to-G editing at A8 to correct the *ACTA2 R179H* mutation (**panel e**), bystander A-to-G editing at A4 that causes an *ACTA2* M178V (**panel f**), bystander A-to-G editing at position A14 that causes *ACTA2* D181G (**panel g**), or bystander C-to-T editing at position C10 that causes a silent *ACTA2* L180L substitution, C-to-A editing that causes an L180M substitution, or C-to-G editing that causes an L180V substitution (**panel h**). Data in all panels from experiments in HEK 293T cells harboring the *ACTA2 R179H* mutation; editing assessed by targeted sequencing; mean, s.e.m., and individual datapoints shown for n = 3 independent biological replicates.

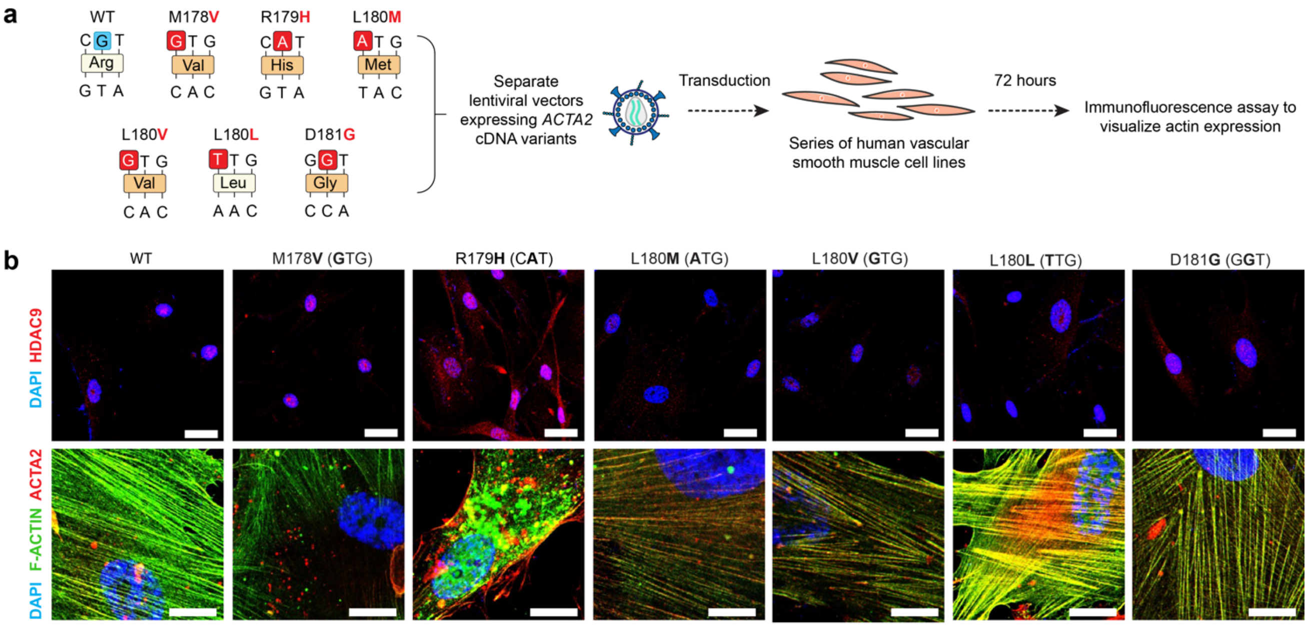


**Supplementary Figure 5. Phenotypic consequences of overexpressing of *ACTA2* variants in primary smooth muscle cells. a,** Schematic of experiments to overexpress mutant *ACTA2* cDNAs in human vascular smooth muscle cells. Various *ACTA2* cDNAs encoding mutations caused by bystander base edits were cloned into lentiviral plasmids. Lentiviral particles were produced, and then used to transduce human vascular smooth muscle cells. Cells were imaged 72 hours after lentiviral transduction. **b**, Representative Immunofluorescence images of primary human smooth muscle cells transduced with lentivirus to overexpress cDNAs encoding various *ACTA2* mutations, including those potentially induced by base editor mediated bystander edits at the on-target site. Putative base edits that would cause each mutation are displayed in the schematic of Fig. 2a. The top panels are representative images of cells stained with DAPI (blue) and HDAC9 (red) antibodies; the bottom panels are representative images of cells stained with DAPI (blue), F-ACTIN (green), and ACTA2 (red) antibodies; scale bars: 80 μm or 30 μm for the top and bottom panels, respectively.

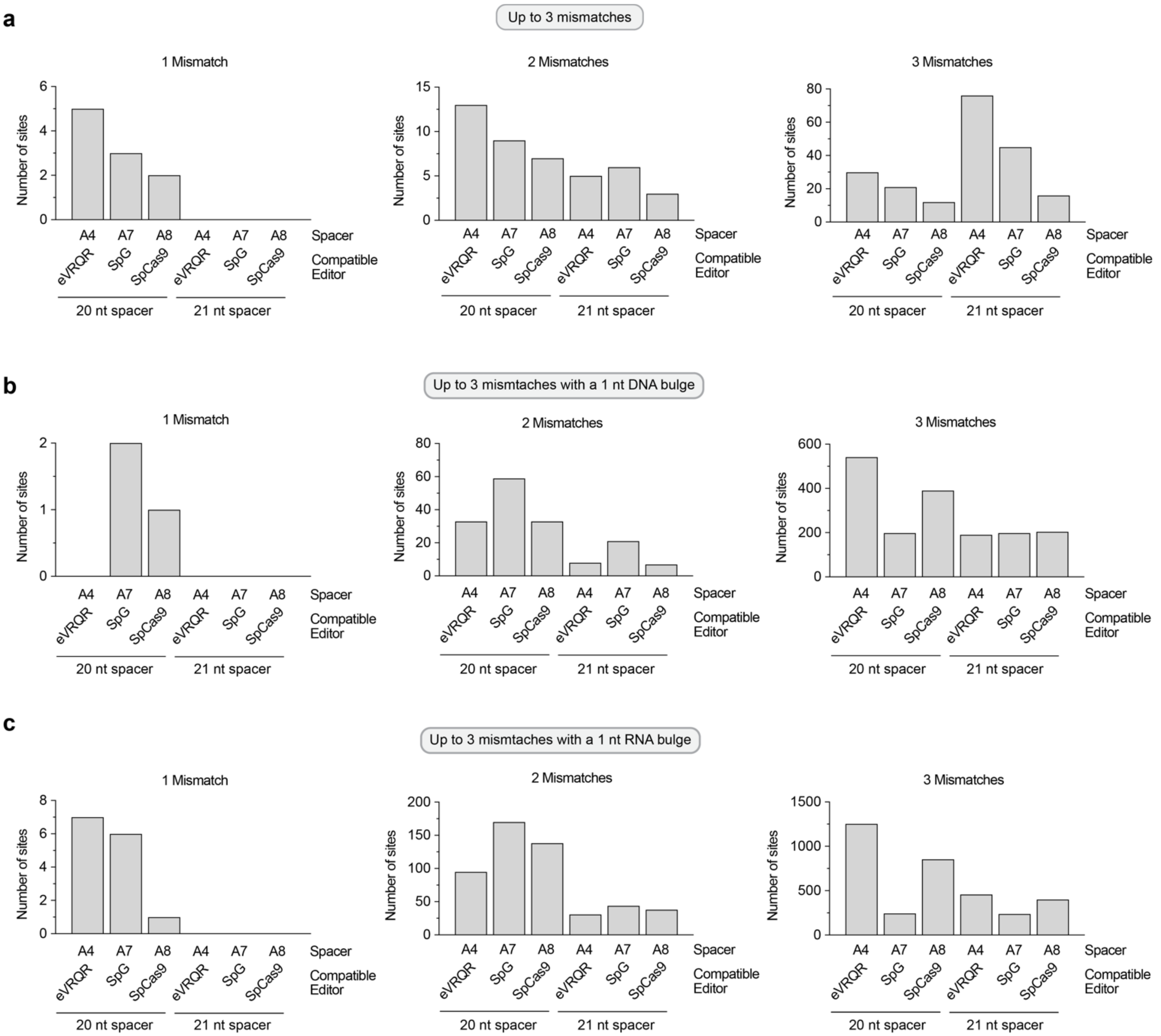


**Supplementary Figure 6. *In silico* annotation of putative off-target sites. a-c**, Putative off-target sites were identified computationally using CasOFFinder[19] for gRNA A4 with NGAN and NGNG PAMs, gRNA A7 with NGNN PAM, and gRNA A8 with NGGN, NAGN and NGAN PAM. Off-target sites were nominated when considering up to 3 mismatches (**panel a**), 3 mismatches with a 1 nt DNA bulge (**panel b**), or 3 mismatches with a 1 nt RNA bulge (**panel c**).

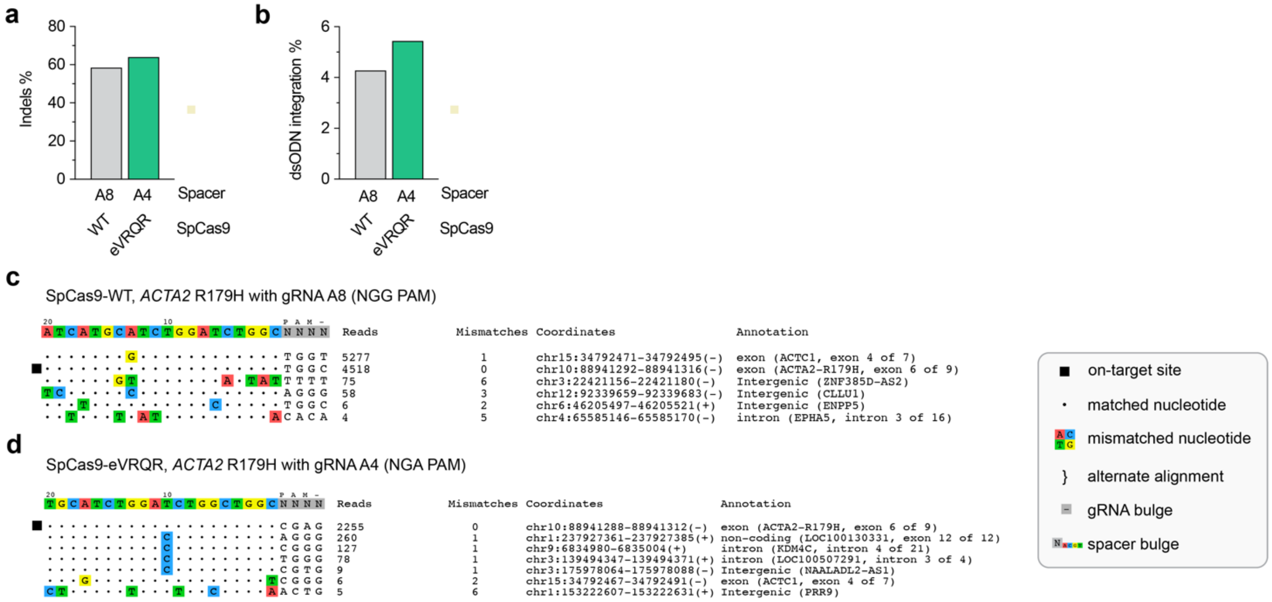


**Supplementary Figure 7. Off-target site nomination using the nuclease-based GUIDE-seq2 assay. a,b,** Results from GUIDE-seq2 experiments including analysis of insertion or deletion mutations (indels; **panel a**) and quantification of reads containing the GUIDE-seq2 dsODN tag (**panel b**) in *ACTA2 R179H* HEK 293T cells when using SpCas9 nuclease with *ACTA2* R179H gRNA A8 or eVRQR nuclease with *ACTA2* R179H gRNA A4. **c,d**, Rank-ordered visualization of genomic on- and off-target sites identified by GUIDE-seq2 experiments performed in *ACTA2* R179H HEK 293T cells transfected with SpCas9 nuclease and *ACTA2* R179H gRNA A8 (**panel c**), or eVRQR nuclease and *ACTA2* R179H gRNA A4 (**panel d**). Mismatched positions in the spacers of the off-target sites are highlighted in color; GUIDE-seq2 read counts from consolidated unique molecular events for each variant are shown to the right of the sequence plots. Datasets represent a single sequencing result from 3 independent biological replicates that were pooled prior to sequencing.

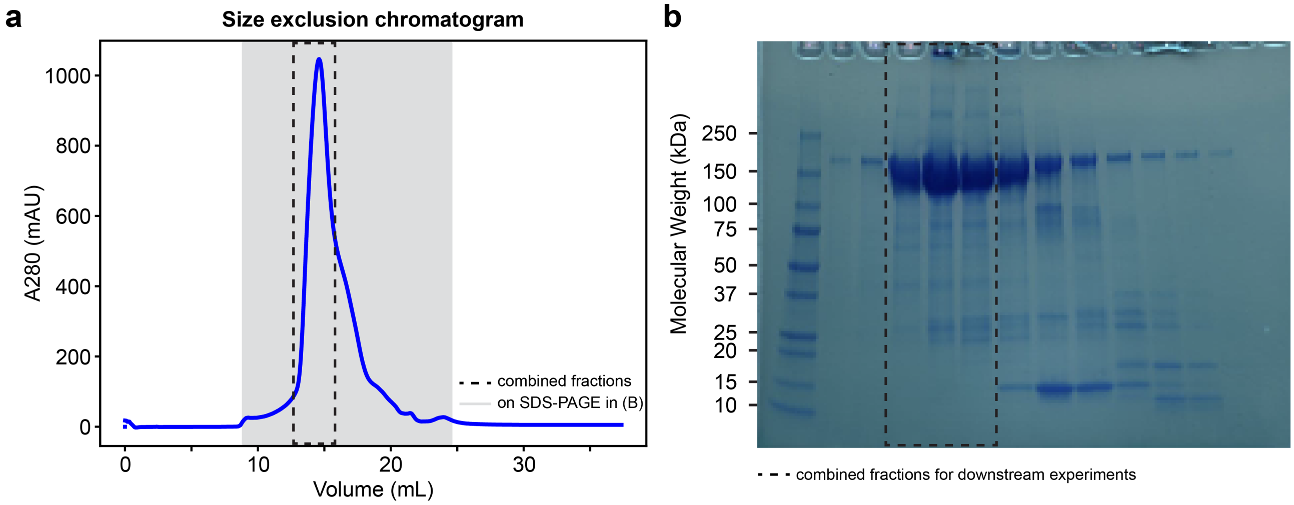


**Supplementary Figure 8. Purification of recombinant ABE8e-eVRQR protein.** (**a**,**b**) Size exclusion chromatography trace (**panel a**) and SDS-PAGE gel (**panel b**) of the elution fractions of from the purification of ABE8e-eVRQR protein. The eluate from the highlighted lanes in **panel b** (between the dashed lines) was collected and stored for experiments.

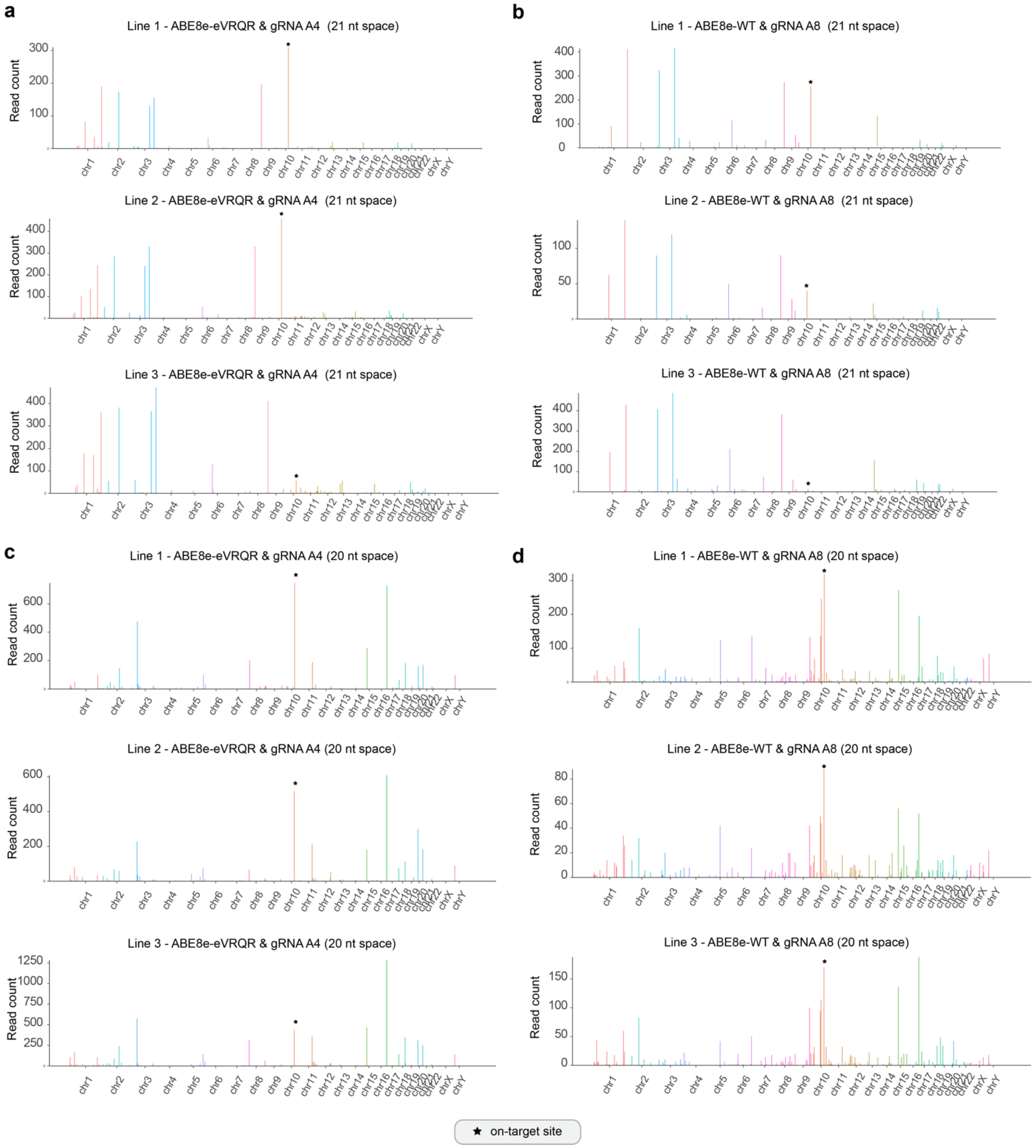


**Supplementary Figure 9. Nomination of base editor-induced off-target sites via CHANGE-seq-BE assay with human genomic DNA.** The CHANGE-seq-BE assay[20] was performed to detect putative off-target sites across the human genome with ABE8e-eVRQR paired with *ACTA2* R179H gRNA A4, or with ABE8e-WT paired with gRNA A8. (**a-d**) Manhattan plots of CHANGE-seq-BE-detected on- and off-target sites for ABE8e-eVRQR and gRNA A4 with a 21 nt spacer (+1 5'G) (**panel a**), ABE8e-WT and gRNA A8 with a 21 nt spacer (+1 5'G) (**panel b**), ABE8e-eVRQR and gRNA A4 with a 20 nt spacer (**panel c**), and ABE8e-WT and gRNA A8 with a 20 nt spacer (**panel d**). The off-target sites are ordered by chromosomal position with bar heights proportional to CHANGE-seq-BE read counts. Experiments were performed using genomic DNA from *ACTA2* R179H MSMDS patient-derived fibroblast lines 1-3. The on-target site is indicated using a black star.

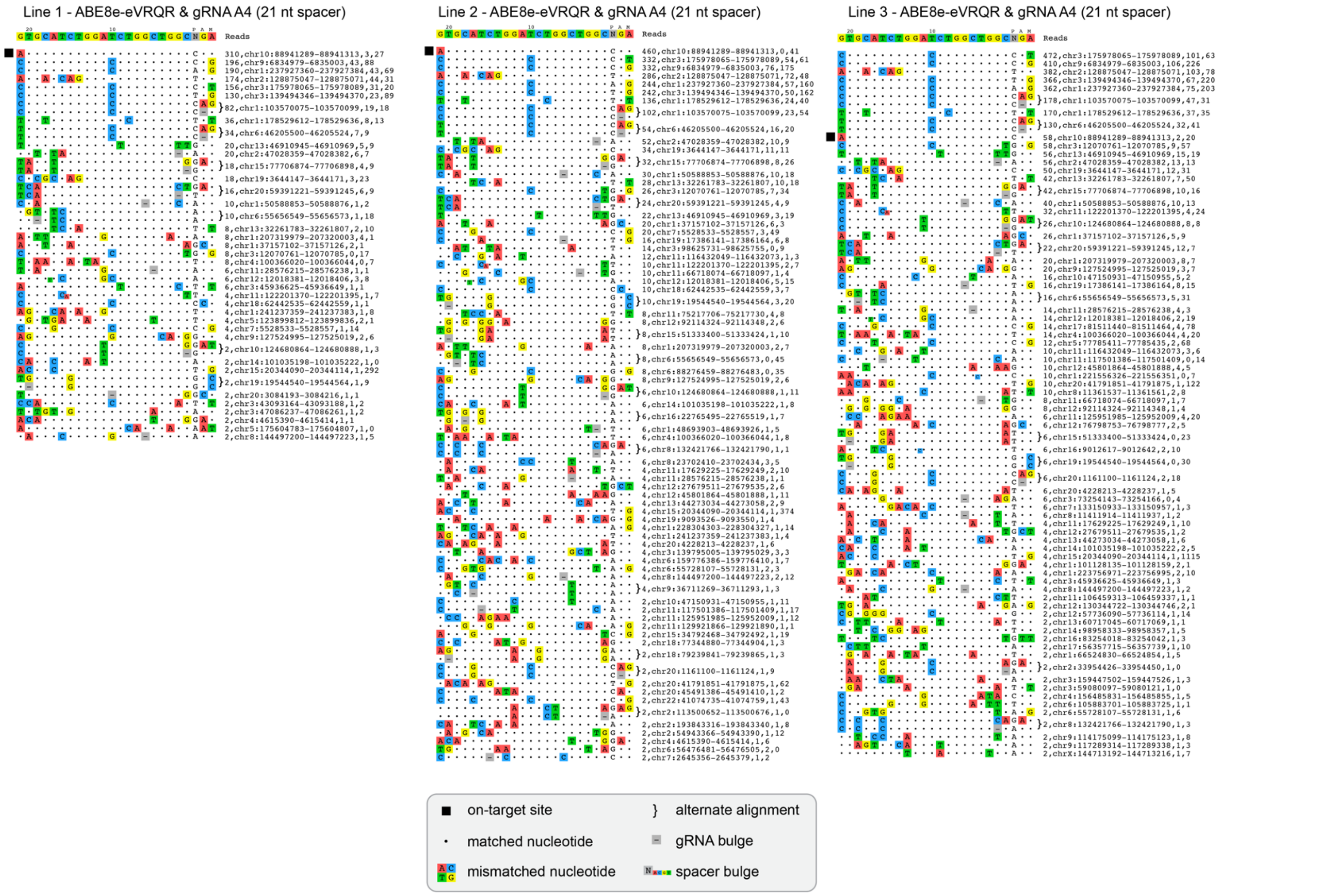


**Supplementary Figure 10. CHANGE-seq-BE detected human genome off-target sites for ABE8e-eVRQR paired with gRNA A4 with a 21-nucleotide spacer (+1 5'G)**. Rank-ordered visualization of on- and off-target genomic sites identified by CHANGE-seq-BE using genomic DNA from three different *ACTA2* R179H MSMDS patient fibroblast lines. The on-target site is indicated using a black box; alternate alignments are shown for sites that are potentially targeted via 1 nt DNA or gRNA spacer bulges.

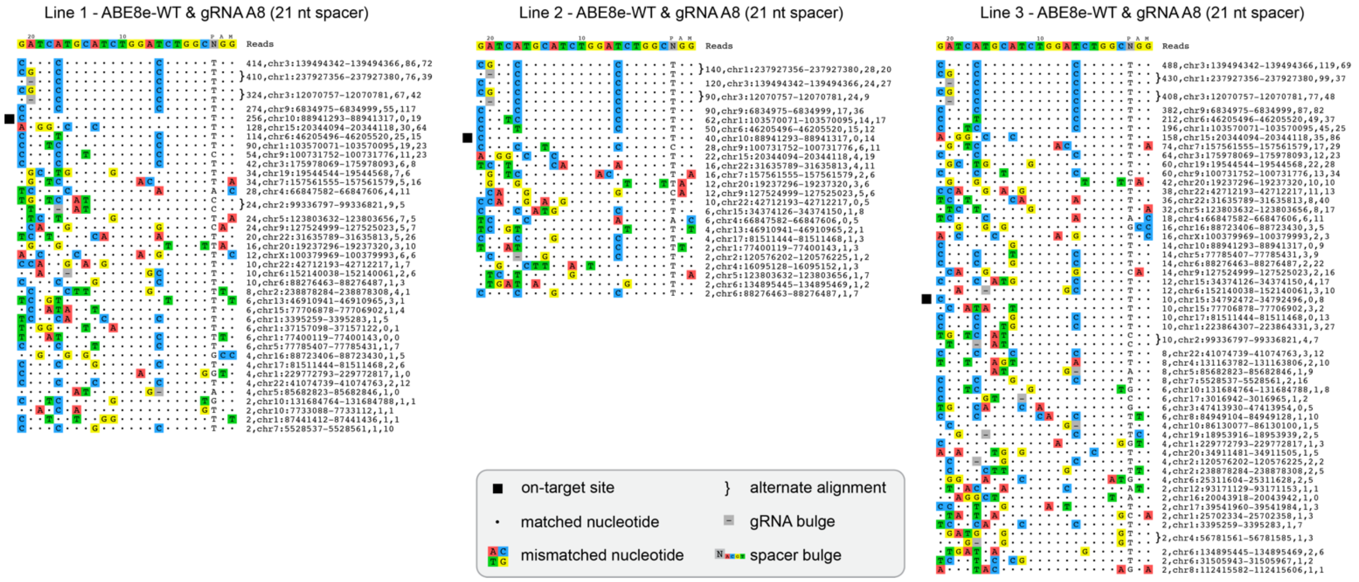


**Supplementary Figure 11. CHANGE-seq-BE detected human genome off-target sites for ABE8e-WT paired with gRNA A8 with a 21-nucleotide spacer (+1 5'G)**. Rank-ordered visualization of on- and off-target genomic sites identified by CHANGE-seq-BE using genomic DNA from three different *ACTA2* R179H MSMDS patient fibroblast lines. The on-target site is indicated using a black box; alternate alignments are shown for sites that are potentially targeted via 1 nt DNA or gRNA spacer bulges.

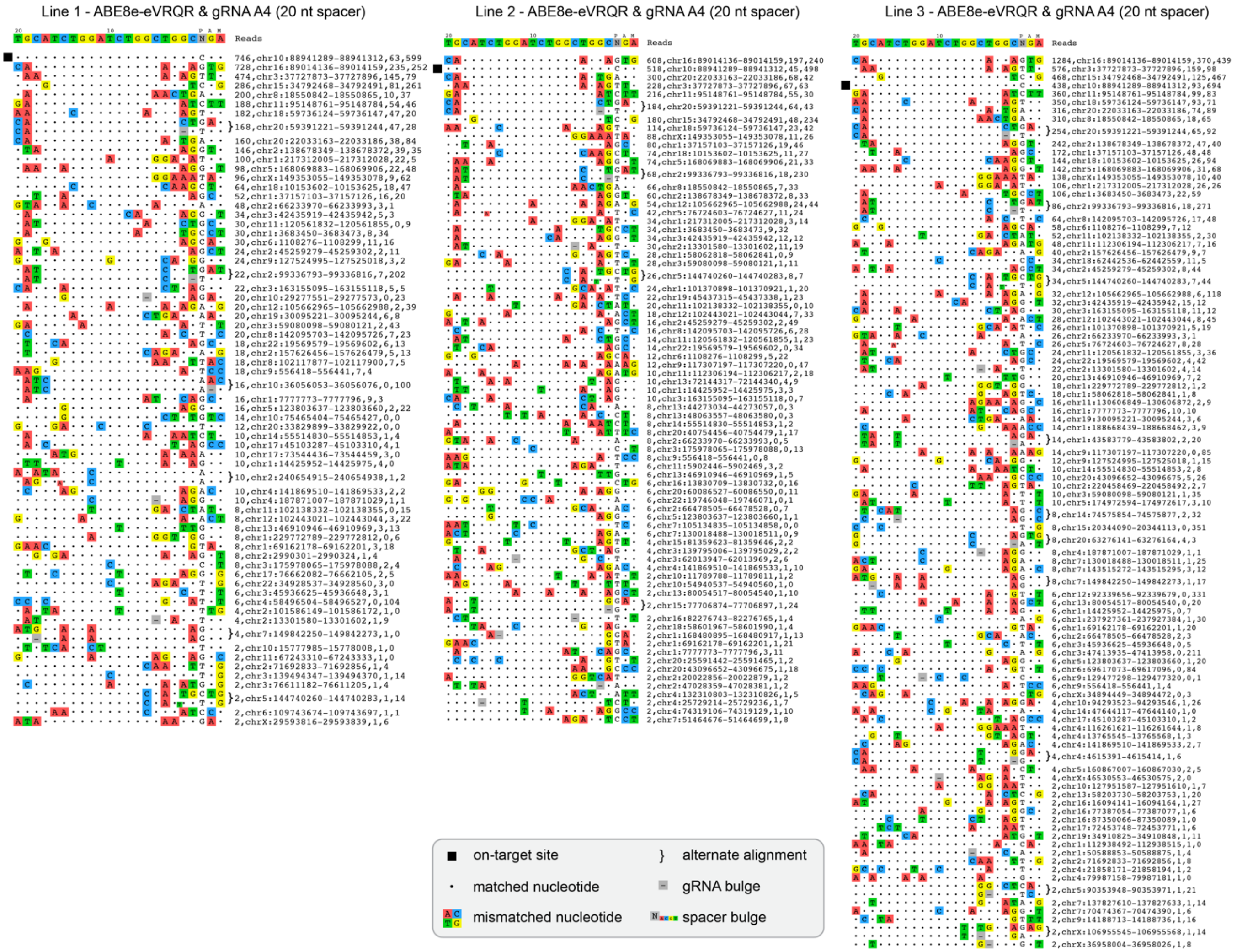


**Supplementary Figure 12. CHANGE-seq-BE detected human genome off-target sites for ABE8e-eVRQR paired with gRNA A4 with a 20-nucleotide spacer**. Rank-ordered visualization of on- and off-target genomic sites identified by CHANGE-seq-BE using genomic DNA from three different *ACTA2* R179H MSMDS patient fibroblast lines. The on-target site is indicated using a black box; alternate alignments are shown for sites that are potentially targeted via 1 nt DNA or gRNA spacer bulges.

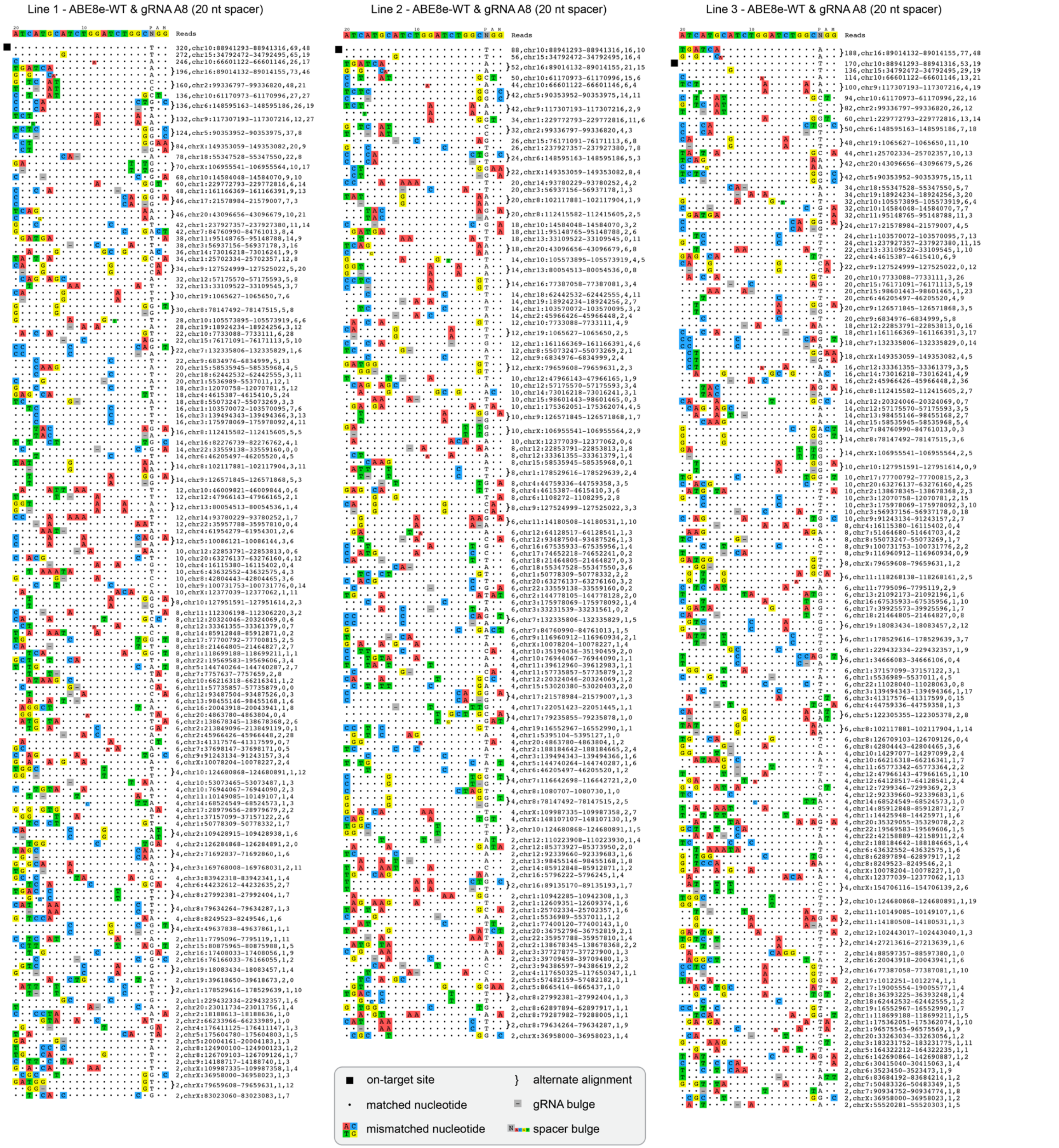


**Supplementary Figure 13. CHANGE-seq-BE detected human genome off-target sites for ABE8e-WT paired with gRNA A8 with a 20-nucleotide spacer**. Rank-ordered visualization of on- and off-target genomic sites identified by CHANGE-seq-BE using genomic DNA from three different *ACTA2* R179H MSMDS patient fibroblast lines. The on-target site is indicated using a black box; alternate alignments are shown for sites that are potentially targeted via 1 nt DNA or gRNA spacer bulges.

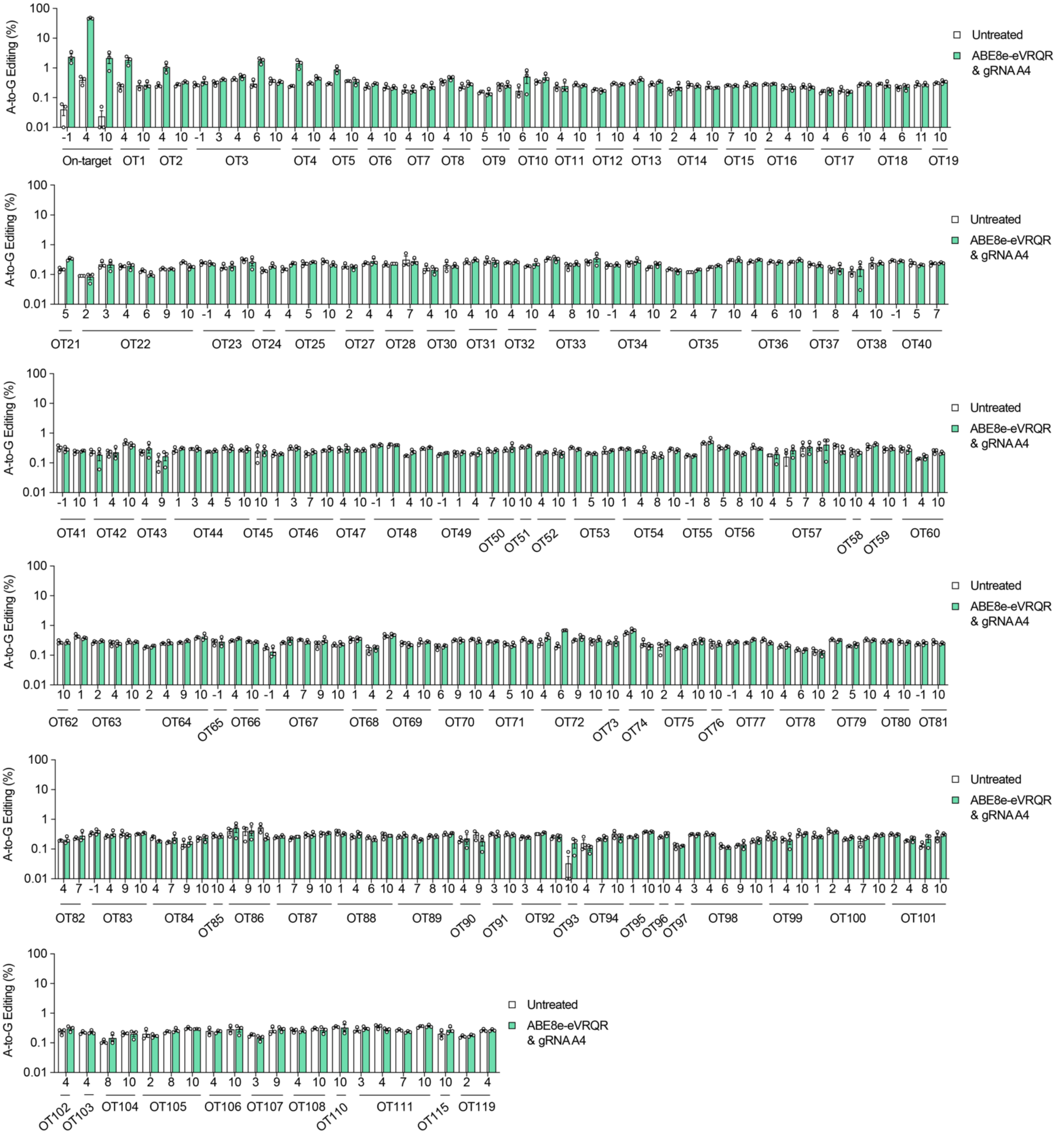


**Supplementary Figure 14. Validation of on- and off-target base editing with ABE8e-eVRQR and *ACTA2* R179H gRNA A4.** The on-target site and 121 off-target sites were sequenced via rhAmpSeq pooled multiplex amplicon sequencing (**Supplementary Table 1**), using genomic DNA extracted from independent replicates of untreated samples or transfections using ABE8e-eVRQR and gRNA A4 in homozygous HEK 293T *ACTA2* R179H cells. Data from the rhAmpSeq output was analyzed using CRISPResso2; the base editing efficiencies were plotted for all adenine bases across a wide edit window (defined as bases 1 through 12 in the target site spacer, counting from the PAM distal end of the spacer); off-target sites were selected based on all sites nominated by CHANGE-seq-BE or GUIDE-seq2, and sites with up to 2 mismatches with up to 1 bulge nominated by CasOFFinder; the genomic regions for off-target sites 20, 29, 39, 61, 109, 112-114, 116-118, 120, and 121 failed to amplify or sequence.

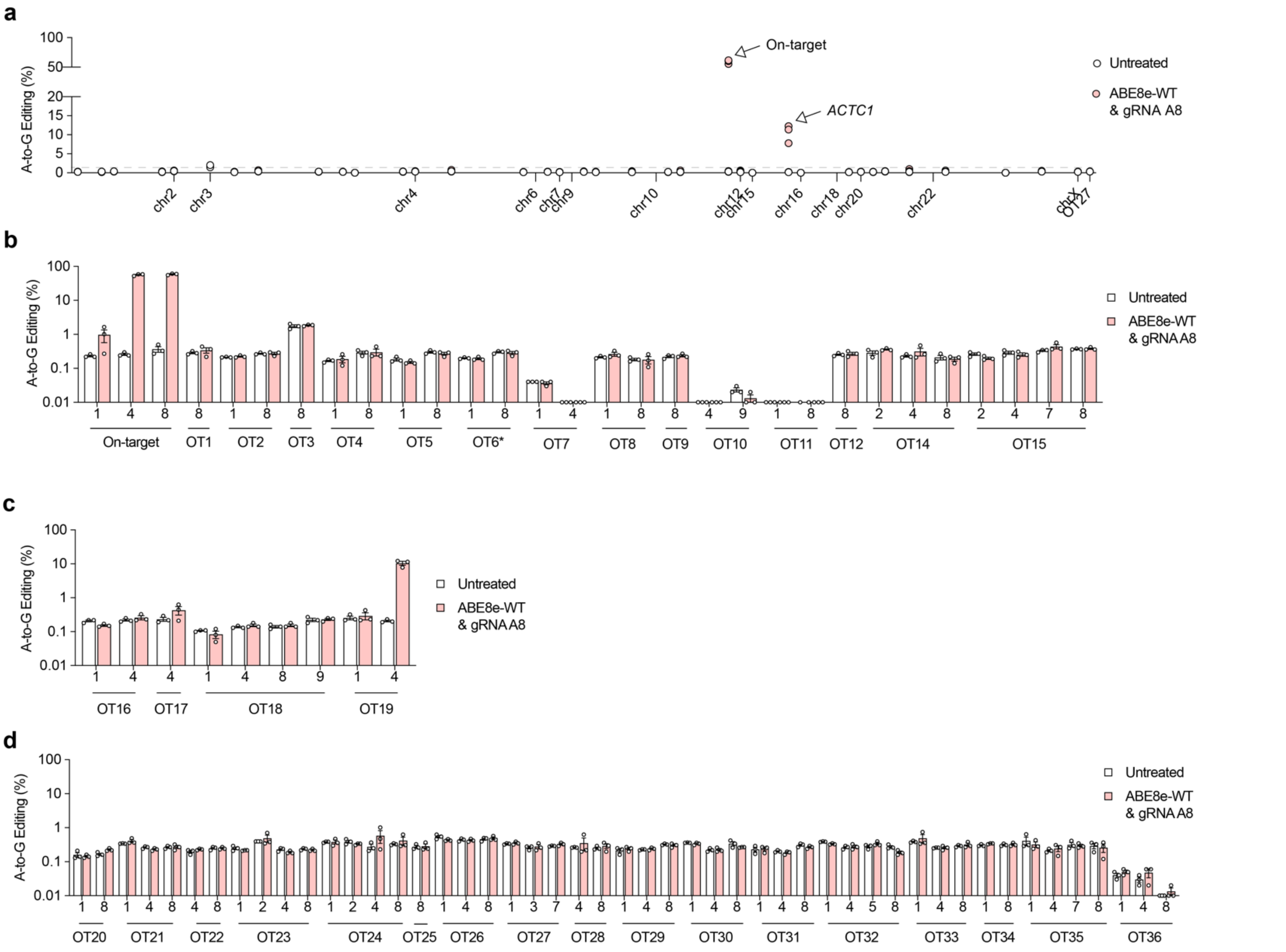


**Supplementary Figure 15. Validation of on- and off-target base editing with ABE8e-WT and *ACTA2* R179H gRNA A8. a**, Summary of on- and off-target base editing in homozygous HEK 293T *ACTA2* R179H cells that were untreated (naïve) or treated with ABE8e-WT and *ACTA2* R179H gRNA A8. Base editing efficiencies are plotted for only the most edited base for each target site, which is typically an adenine in the middle of the optimal base editor edit window. The *ACTA2* on-target site and a prevalent off-target within the *ACTC1* gene (causing an M178V mutation) are indicated with arrows. The on- and off-target sites are ordered based on chromosomal positions. **b-d,** Analysis of on- and off-target base editing via rhAmpSeq using genomic DNA from HEK 293T *ACTA2* R179H cells that were untreated (naïve) or treated with ABE8e-WT and *ACTA2* R179H gRNA A8; sequencing was performed at the on-target site and 36 off-targets nominated by CHANGE-seq-BE (only those with more than 1% of CHANGE-seq-BE reads were selected for validation using the A8 gRNA; **panel b**), GUIDE-seq2 detected sites (**panel c**), or CasOFFinder annotated sites (**panel d**), with data analysis performed via CRISPResso2[21] for n = 3 independent biological replicates. The genomic region for off-target site 13 failed to amplify or sequence.

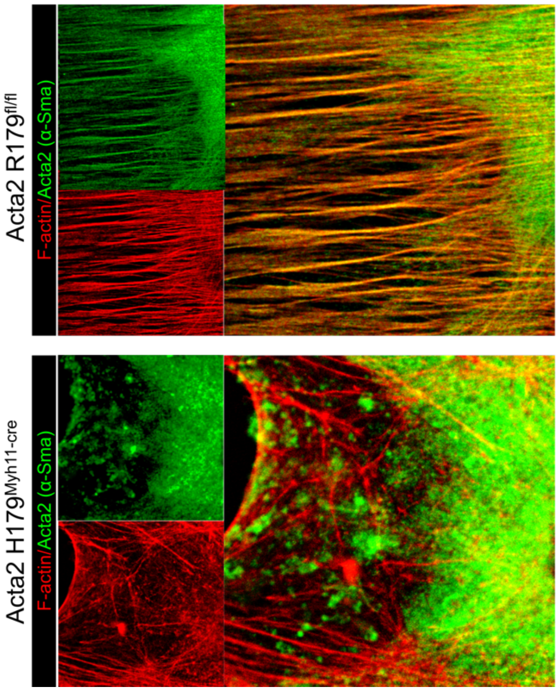


**Supplementary Figure 16. Impact of the *Acta2* R179H mutation on the cellular cytoskeleton.** Representative immunofluorescence images of aortic vascular smooth muscle cells (VSMCs) from control (Acta2 R179fl/fl) and MSMDS (Myh11-Cre R179H/fl/+) mice (top and bottom panels, respectively). VSMCs were stained with rhodamine-labeled phalloidin (red) to visualize F-actin stress fibers, α-Smooth Muscle Actin (α-SMA, green) to assess smooth muscle-specific actin organization, and DAPI (blue) for nuclei. Cells were fixed and blocked with 10% donkey serum, incubated overnight at 4°C with primary antibodies (ACTA2 monoclonal, 1:200) and an F-actin probe (2 drops/mL). Following washes with PBS-tween, secondary antibodies (1:400) were applied for 1 hour at room temperature. After mounting with DAPI-containing medium, images were acquired using a Leica TCS SP8 confocal microscope. The merged panels are overlays of the F-actin and α-SMA channels, illustrating potential colocalization of actin fibers with smooth muscle markers. In the control group (Acta2 R179^fl/fl^), well-organized F-actin stress fibers colocalize with α-SMA along the filaments. In the mutant group (Myh11-Cre^R179H/fl/+), there are fewer stress fibers, with disrupted colocalization of α-SMA and actin filaments, showing impaired cytoskeletal structure.

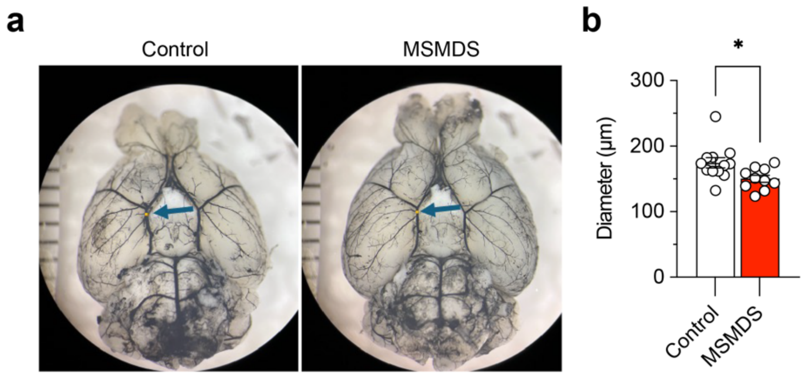


**Supplementary Figure 17. Distal internal carotid artery (ICA) diameter measurements in Acta2fl/+ (control) and Acta2fl+/Myh11Cre+ (MSMDS) mice.** Transcardiac perfusion was performed with black ink and brains were collected. **a,** Representative photographs of the circle of Willis. The bars and blue arrow points indicate the distal ICA. **b**, The diameter of the distal internal carotid arteries (ICA) from both left and right hemispheres was measured using ImageJ (NIH, Bethesda, MD, USA), and used to calculate the average of both sides. The diameters were significantly different between control and MSMDS mice (t-test). mean, s.e.m., and individual datapoints shown for n = 10-12 mice.

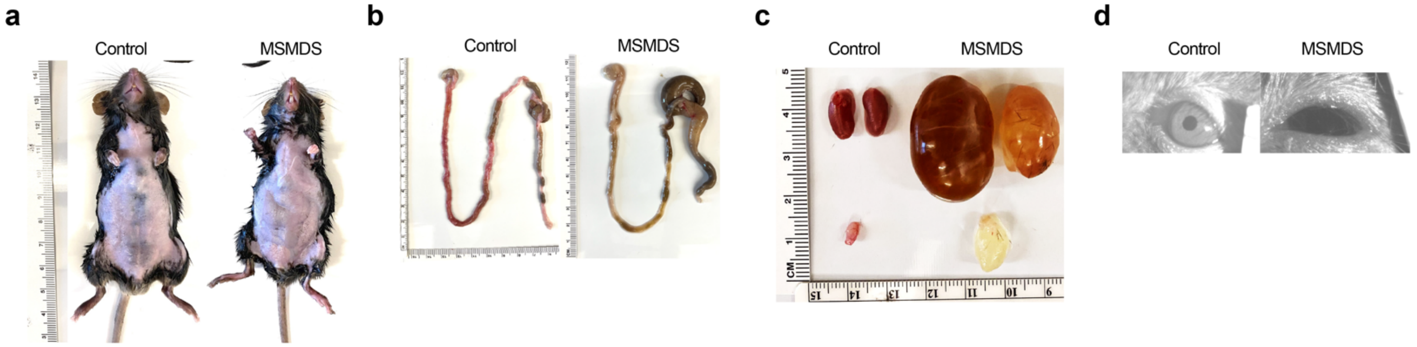


**Supplementary Figure 18. Macro analysis of Acta2fl/+ (Control) and Acta2fl+/Myh11Cre+ (MSMDS) mice.** **a-d**, Representative images comparing anatomy of 7-week-old control and MSMDS mice, including distended abdomen (**panel a**), fecal impaction (**panel b**), hydronephrotic kidneys (top) and distended bladder (bottom) (**panel c**), and pupil dilatation (**panel d**).

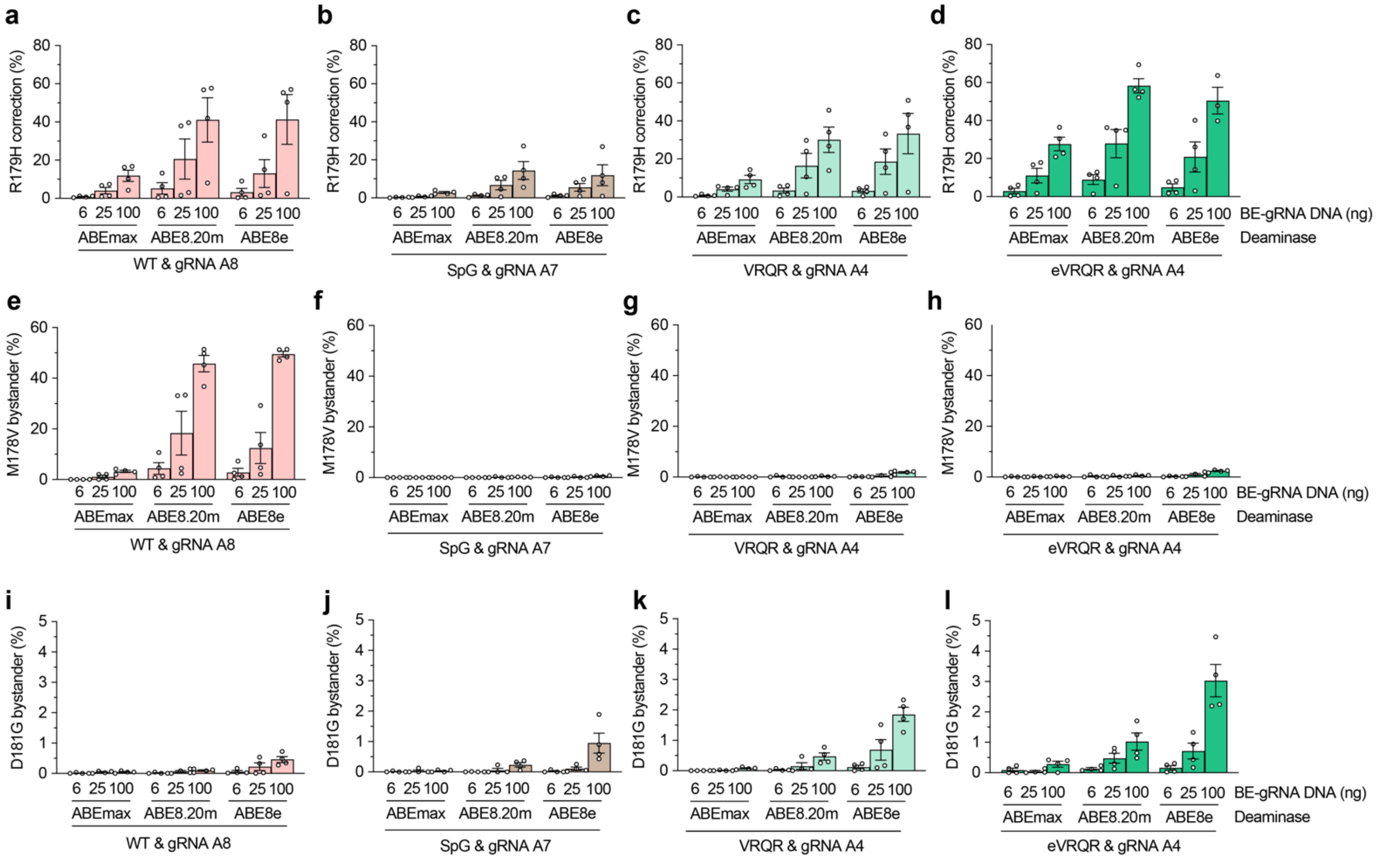


**Supplementary Figure 19. Correction of *ACTA2* R179H via base editing when titrating the dose of dual-AAV plasmids.** Heterozygous *ACTA2* R179H HEK 293T cells were transfected with dual AAV intein-split expression plasmids encoding the ABE and gRNA, titrated at different DNA amounts (including stuffer DNA to maintain the total amount of transfected DNA at 100 ng per 96-well of cells). The N- and C-terminal AAV plasmids were transfected at a 1:1 ratio by weight. **a-d**, On-target A-to-G editing of the *ACTA2* R179H target adenine when using different engineered TadA deaminases[10,18,22,23] fused to WT SpCas9 and paired with gRNA A8 (**panel a**), fused to SpG[2] and paired with gRNA A7 (**panel b**), fused to SpCas9-VRQR[11,12] and paired with gRNA A4 (**panel c**), and fused to eVRQR and paired with gRNA A4 (**panel d**). **e-h,** Bystander A-to-G editing causing *ACTA2* M178V (base A-1 with gRNA A4; A3 with gRNA A7; A4 with gRNA A8) when using different engineered TadA deaminases fused to SpCas9 and paired with gRNA A8 (**panel e**), fused to SpG and paired with gRNA A7 (**panel f**), fused to SpCas9-VRQR and paired with gRNA A4 (**panel g**), and fused to eVRQR and paired with gRNA A4 (**panel h**). (**i-l**) Bystander A-to-G editing causing *ACTA2* D181G (base A10 with gRNA A4; A13 with gRNA A7; A14 with gRNA A8) when using different deaminases fused to WT SpCas9 and paired with gRNA A8 (**panel i**), fused to SpG and paired with gRNA A7 (**panel j**), fused to SpCas9-VRQR and paired with gRNA A4 (**panel k**), and fused to eVRQR and paired with gRNA A4 (**panel l**). Editing assessed by targeted sequencing; mean, s.e.m., and individual datapoints shown for n = 3 independent biological replicates**.**

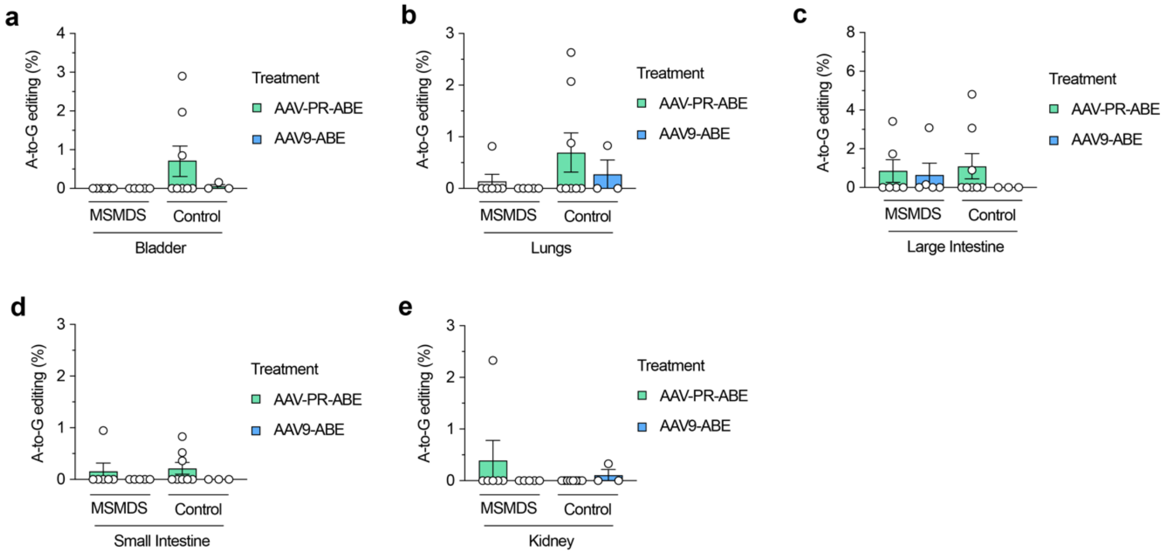


**Supplementary Figure 20. *In vivo* on-target A-to-G correction of *Acta2* R179H**. **a-e**, Tissues from MSMDS (Myh11-Cre:Acta2fl/+) or control (*Acta2fl*/+) mice treated with dual AAV vectors encoding ABE8e-eVRQR and *Acta2* R179H gRNA A4 were assessed for on-target A-to-G editing at position A4 of the target site in the bladder, lungs, large intestine, small intestine, and kidney (**panels a-e**, respectively). Tissues were extracted after 8 weeks following IV retro-orbital injections of AAV-PR-ABE or AAV9-ABE vectors into P3 control or MSMDS mice and subjected to targeted sequencing. Mean, s.e.m., and individual datapoints shown in all panels for n = 6 mice.

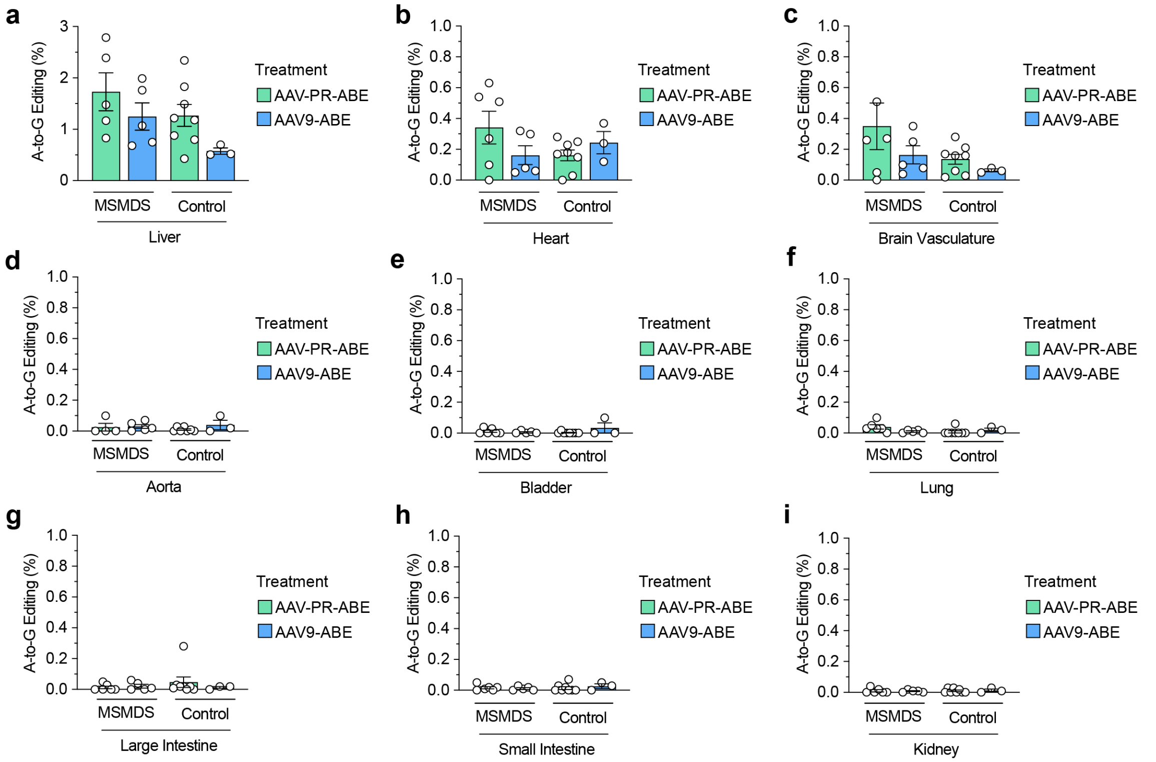


**Supplementary Figure 21. *In vivo* bystander A-to-G editing causing an *Acta2* M178V mutation**. **a-i**, Tissues from MSMDS (Myh11-Cre:Acta2fl/+) or control (*Acta2fl*/+) mice treated with dual AAV vectors encoding ABE8e-eVRQR and *Acta2* R179H gRNA A4 were assessed for bystander A-to-G editing at position A-1 of the target site in the liver, heart, brain vasculature, aorta, bladder, lungs, large intestine, small intestine, and kidney (**panels a-i**, respectively). Tissues were extracted after 8 weeks following IV retro-orbital injections of AAV-PR-ABE or AAV9-ABE vectors into P3 control or MSMDS mice and subjected to targeted sequencing. Mean, s.e.m., and individual datapoints shown in all panels for n = 6 mice.

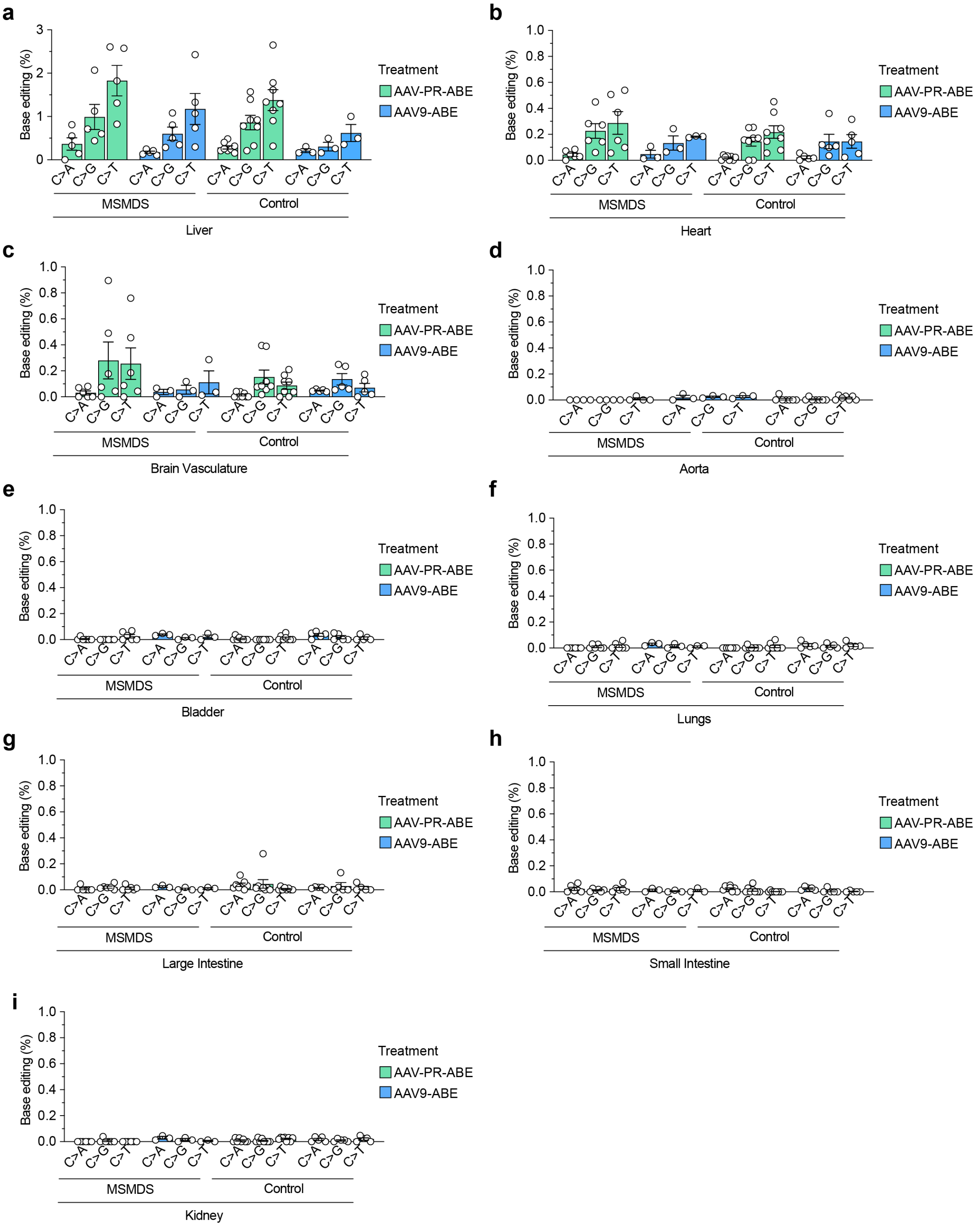


**Supplementary Figure 22. *In vivo* cytosine bystander editing causing various *Acta2* L180 mutations**. **a-i**, Tissues from MSMDS (Myh11-Cre:Acta2fl/+) or control (*Acta2fl*/+) mice treated with dual AAV vectors encoding ABE8e-eVRQR and *Acta2* R179H gRNA A4 were assessed for bystander C-to-A, C-to-G, or C-to-T editing at position C6 of the target site (causing *ACTA2* L180M, L180V, and a silent L180L mutation, respectively) in the liver, heart, brain vasculature, aorta, bladder, lungs, large intestine, small intestine, and kidney (**panels a-i**, respectively). Tissues were extracted after 8 weeks following IV retro-orbital injections of AAV-PR-ABE or AAV9-ABE vectors into P3 control or MSMDS mice and subjected to targeted sequencing. Mean, s.e.m., and individual datapoints shown in all panels for n = 6 mice.

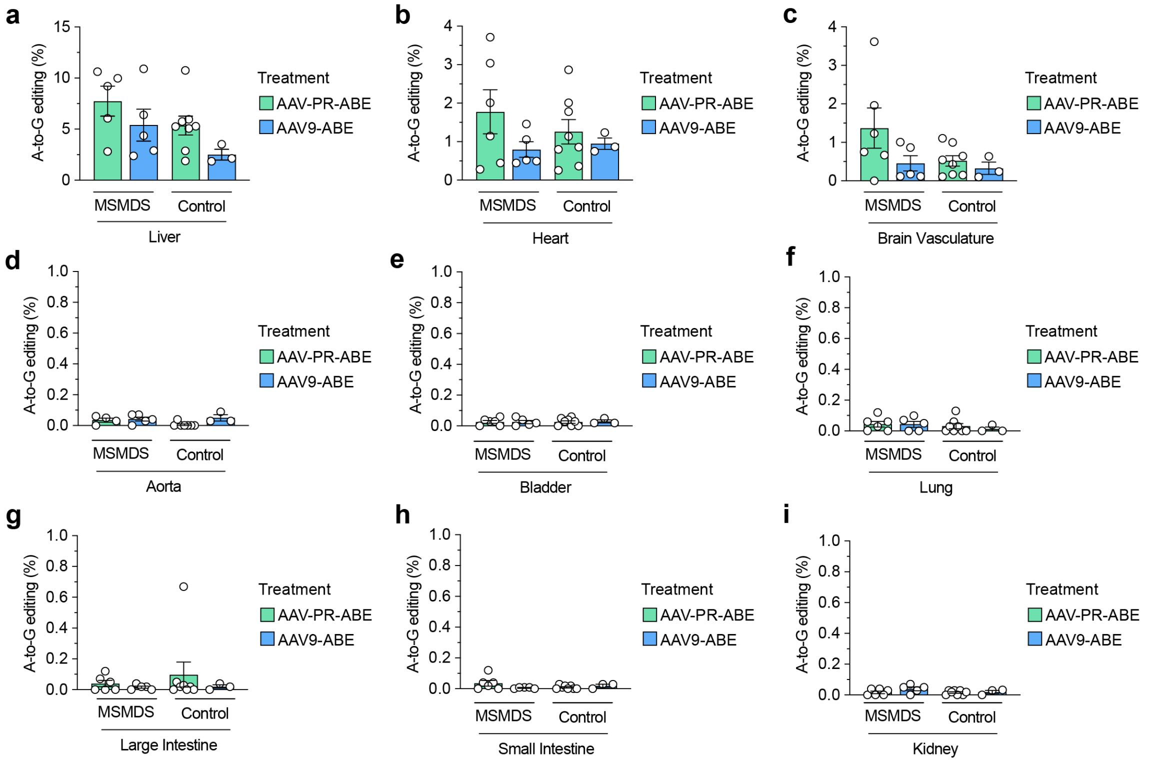


**Supplementary Figure 23. *In vivo* bystander A-to-G editing causing an *Acta2* D181G mutation**. **a-i**, Tissues from MSMDS (Myh11-Cre:Acta2fl/+) or control (*Acta2fl*/+) mice treated with dual AAV vectors encoding ABE8e-eVRQR and *Acta2* R179H gRNA A4 were assessed for bystander A-to-G editing at position A10 of the target site in the liver, heart, brain vasculature, aorta, bladder, lungs, large intestine, small intestine, and kidney (**panels a-i**, respectively). Tissues were extracted after 8 weeks following IV retro-orbital injections of AAV-PR-ABE or AAV9-ABE vectors into P3 control or MSMDS mice and subjected to targeted sequencing. Mean, s.e.m., and individual datapoints shown in all panels for n = 6 mice.

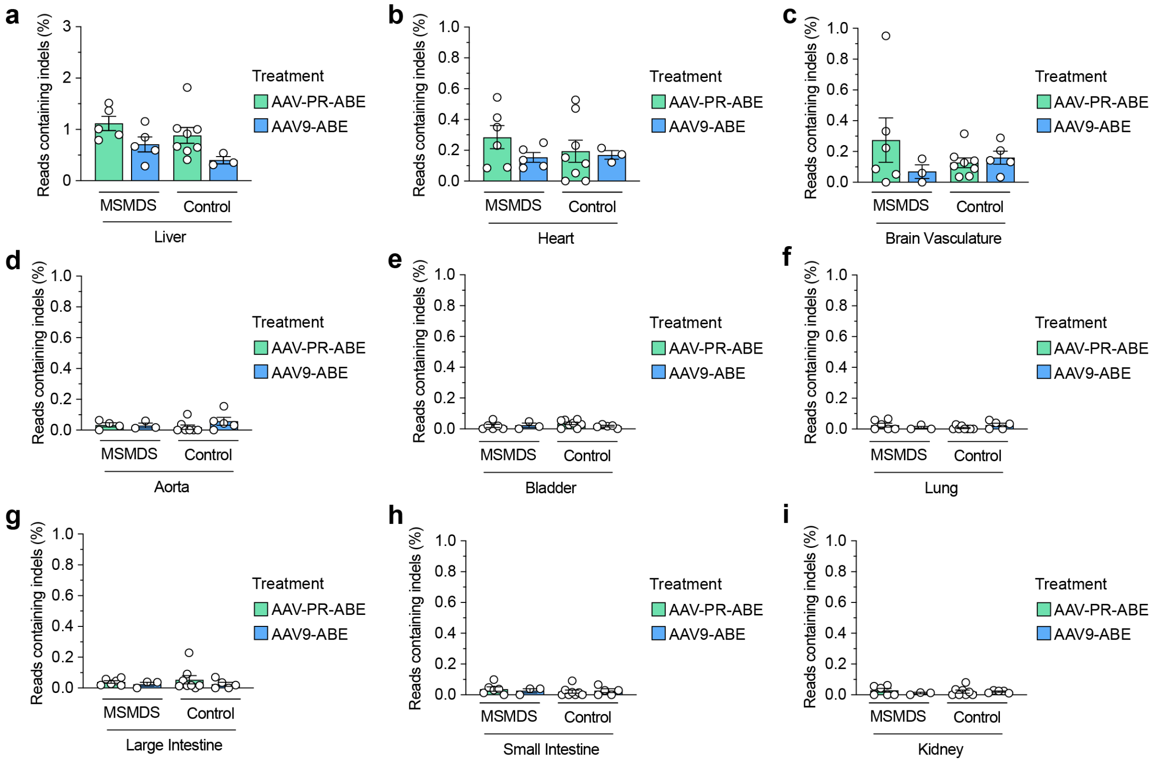


**Supplementary Figure 24. *In vivo* insertion or deletion mutations (indels) at the *ACTA2* R179H target site**. **a-i**, Tissues from MSMDS (Myh11-Cre:Acta2fl/+) or control (*Acta2fl*/+) mice treated with dual AAV vectors encoding ABE8e-eVRQR and *Acta2* R179H gRNA A4 were assessed for indels at the target site in the liver, heart, brain vasculature, aorta, bladder, lungs, large intestine, small intestine, and kidney (**panels a-i**, respectively). Tissues were extracted after 8 weeks following IV retro-orbital injections of AAV-PR-ABE or AAV9-ABE into P3 control or MSMDS mice and subjected to targeted sequencing. Mean, s.e.m., and individual datapoints shown in all panels for n = 6 mice.

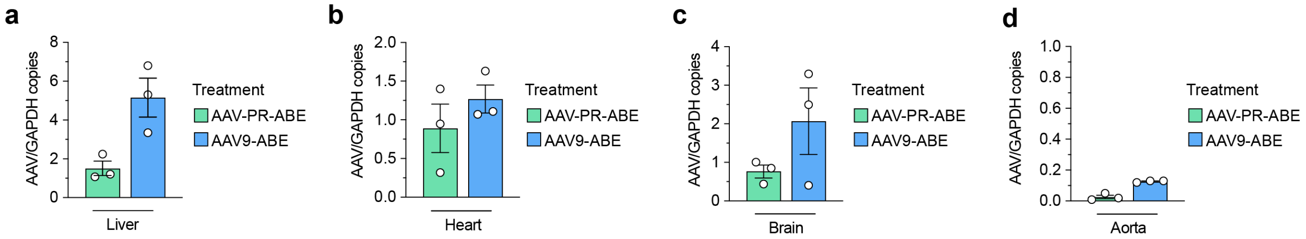


**Supplementary Figure 25. Quantification of AAV-ABE genome copies in MSMDS mice. a-d,** AAV vector genome copies in Myh11-Cre:Acta2fl/+ MSMDS mice quantified 8 weeks following AAV-mediated delivery of ABE8e-eVRQR and *Acta2* R179H gRNA A4 via AAV9 or AAV-PR capsids via retro-orbital injections into P3 MSMDS mice, assessed by ddPCR and normalized by intracellular genomic GAPDH in liver, heart, brain and aorta (**panels a-d**, respectively). Mean, s.e.m., and individual datapoints shown in all panels for n = 3 mice.

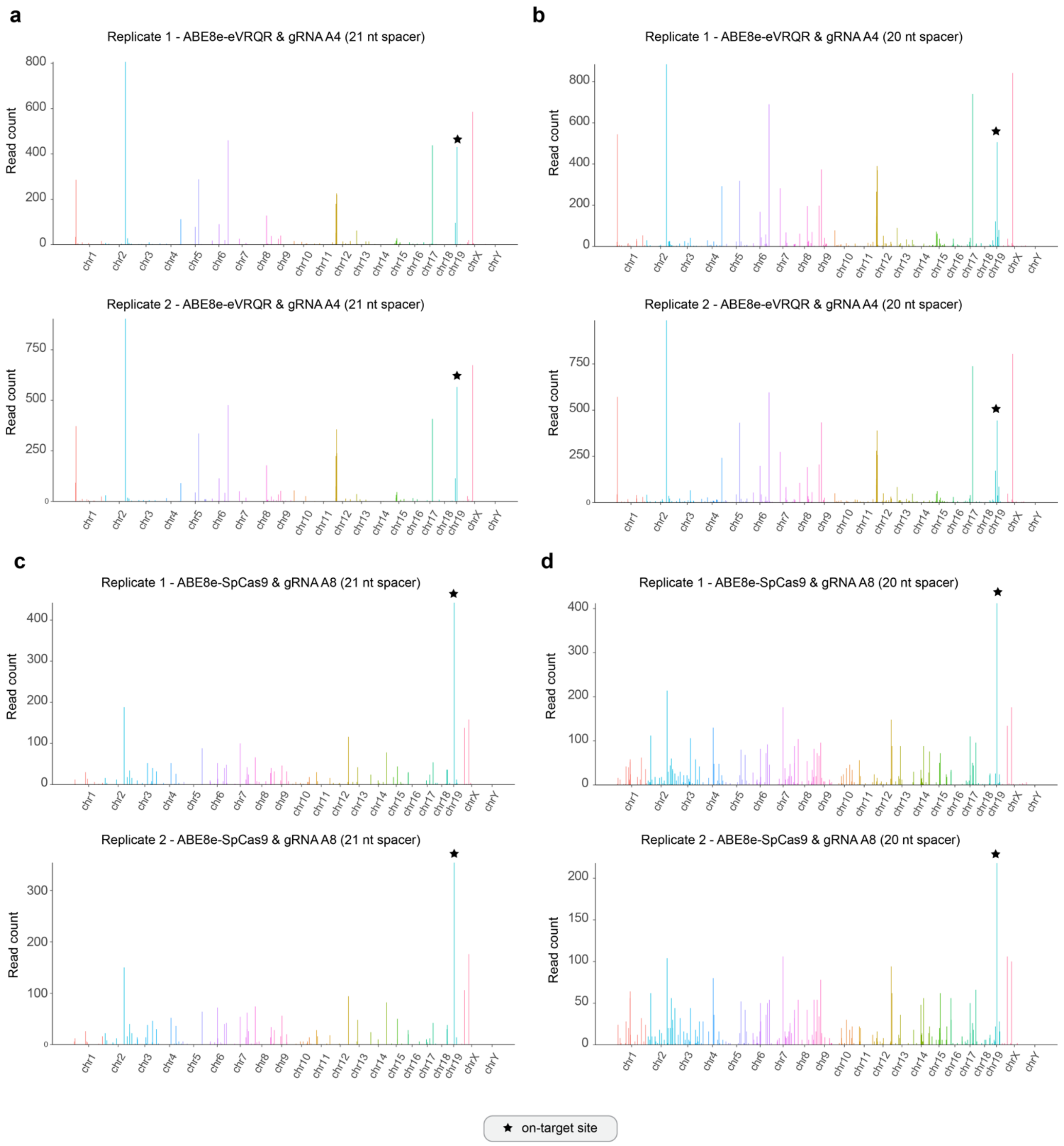


**Supplementary Figure 26. Nomination of ABE-induced off-target sites in the mouse genome.** The CHANGE-seq-BE assay was performed to nominate putative off-target sites in the mouse genome with ABE8e-eVRQR paired with *Acta2* R179H gRNA A4, and with ABE8e-WT paired with gRNA A8. (**a-d**) Manhattan plots of CHANGE-seq-BE-detected on- and off-target sites for ABE8e-eVRQR and gRNA A4 with a 21 nt spacer (+1 5'G) (**panel a**), ABE8e-eVRQR and gRNA A4 with a 20 nt spacer (**panel c**), ABE8e-WT and gRNA A8 with a 21 nt spacer (+1 5'G) (**panel c**), and ABE8e-WT and gRNA A8 with a 20 nt spacer (**panel d**). Off-target sites are ordered by chromosomal position with bar heights proportional to CHANGE-seq-BE read counts. Biological duplicate experiments were performed using genomic DNA from Myh11-cre:Acta2fl/+ MSMDS mice (*Acta2* R179H). The on-target site is indicated using a black star.

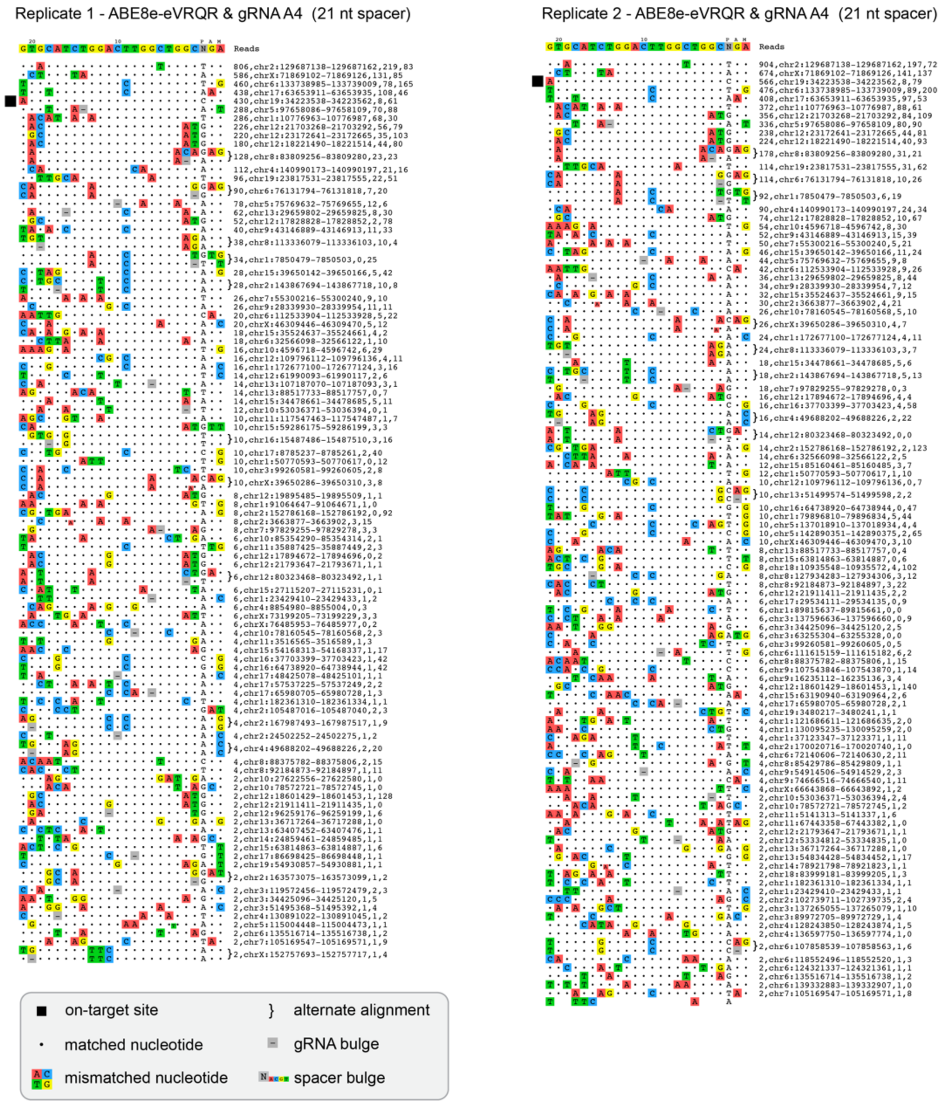


**Supplementary Figure 27. CHANGE-seq-BE detected sites in the mouse genome for ABE8e-eVRQR paired with gRNA A4 with a 21-nucleotide spacer (+1 5'G)**. Rank-ordered visualization of on- and off-target genomic sites identified by CHANGE-seq-BE using genomic DNA from Myh11-cre:Acta2fl/+ MSMDS mice. The on-target site is indicated using a black box; alternate alignments are shown for sites that are potentially targeted via 1 nt DNA or gRNA spacer bulges; genomic locations are listed.

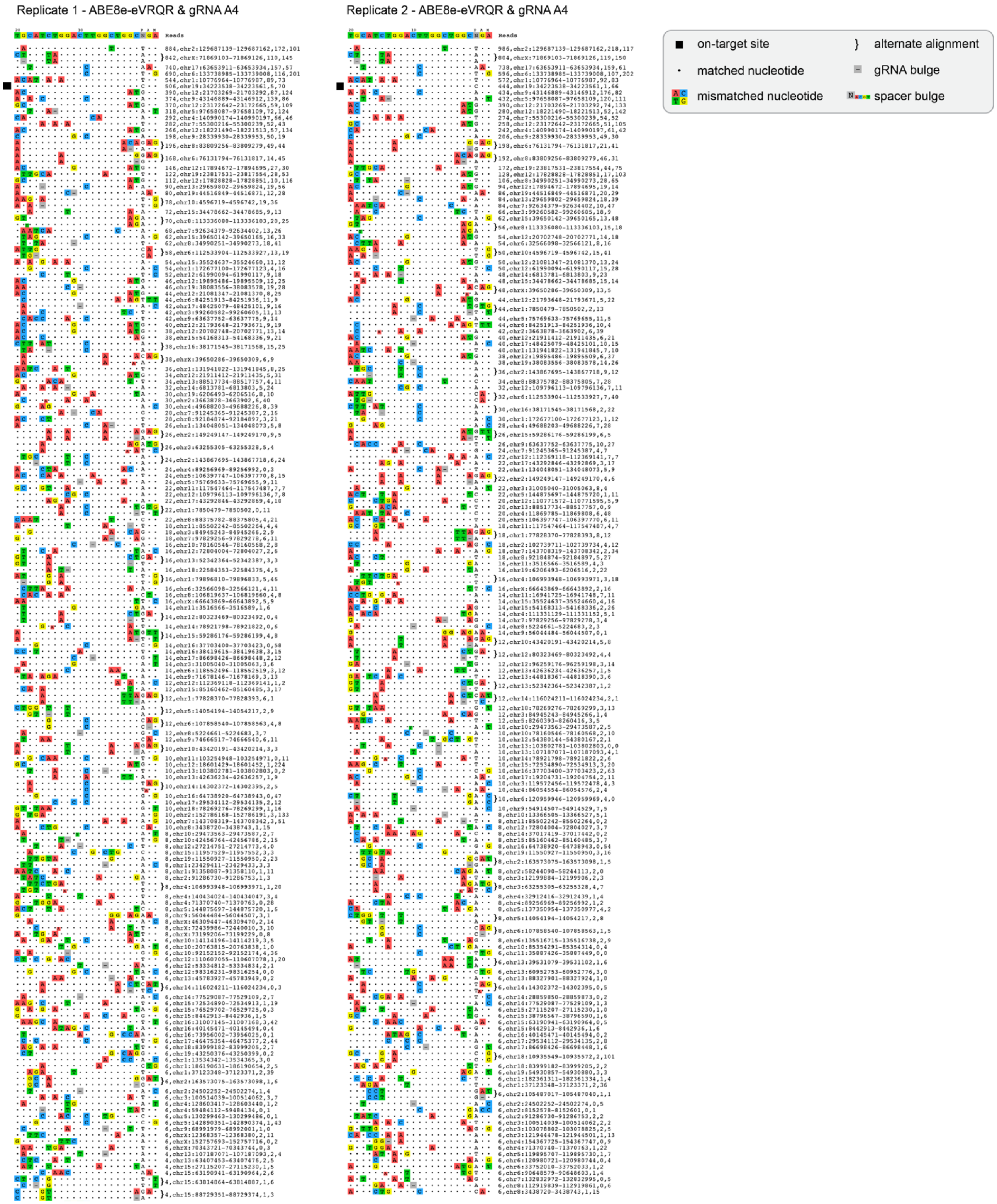


**Supplementary Figure 28. CHANGE-seq-BE detected sites in the mouse genome for ABE8e-eVRQR paired with gRNA A4 with a 20-nucleotide spacer**. Rank-ordered visualization of on- and off-target genomic sites identified by CHANGE-seq-BE using genomic DNA from Myh11-cre:Acta2fl/+ MSMDS mice. The on-target site is indicated using a black box; alternate alignments are shown for sites that are potentially targeted via 1 nt DNA or gRNA spacer bulges; genomic locations are listed. Lower efficiency off-target sites not shown for brevity; the full set of off-target sites are available in **Supplementary Table 1**.

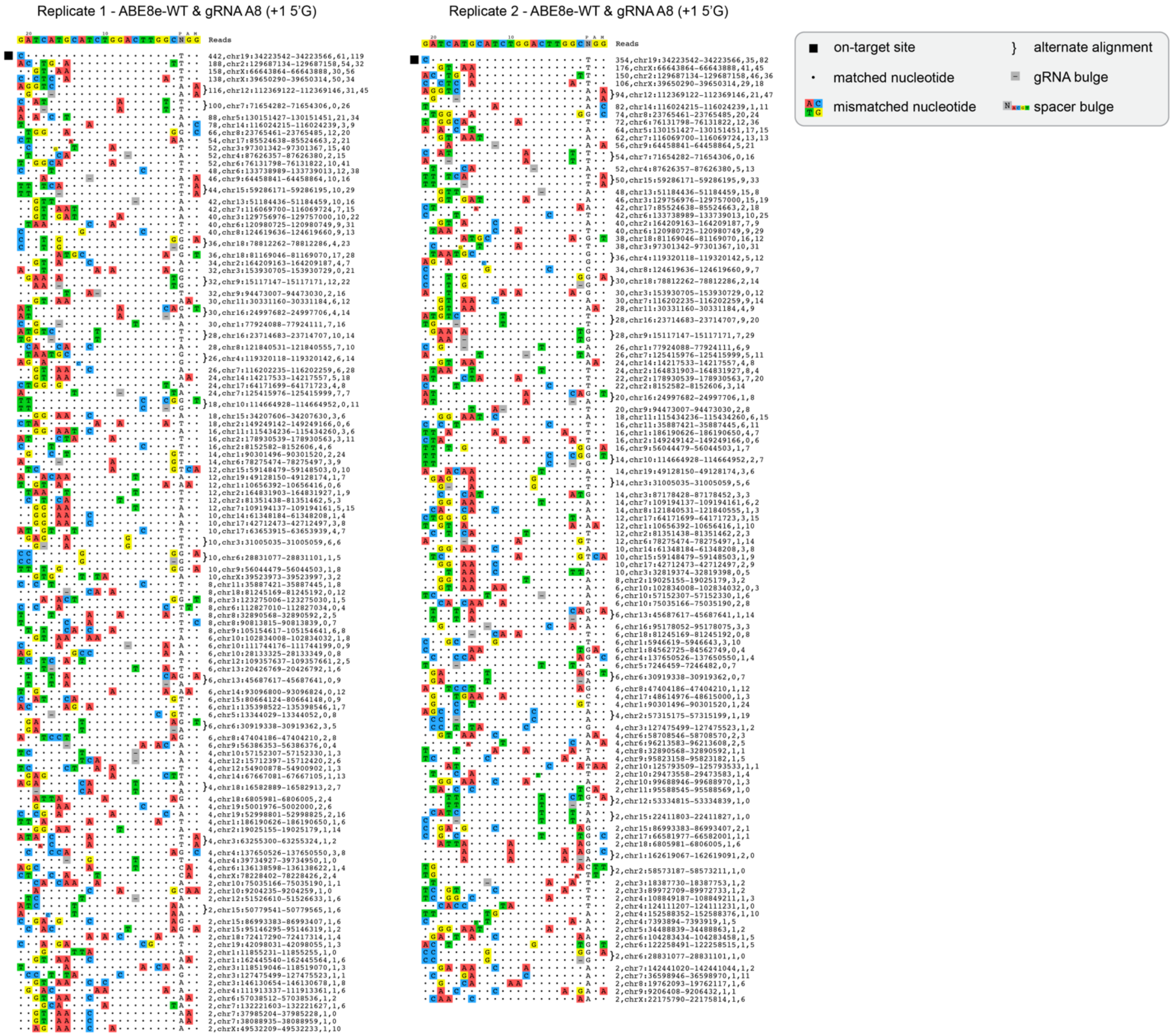


**Supplementary Figure 29. CHANGE-seq-BE detected sites in the mouse genome for ABE8e-WT paired with gRNA A8 with a 21-nucleotide spacer (+1 5'G)**. Rank-ordered visualization of on- and off-target genomic sites identified by CHANGE-seq-BE using genomic DNA from Myh11-cre:Acta2fl/+ MSMDS mice. The on-target site is indicated using a black box; alternate alignments are shown for sites that are potentially targeted via 1 nt DNA or gRNA spacer bulges; genomic locations are listed.

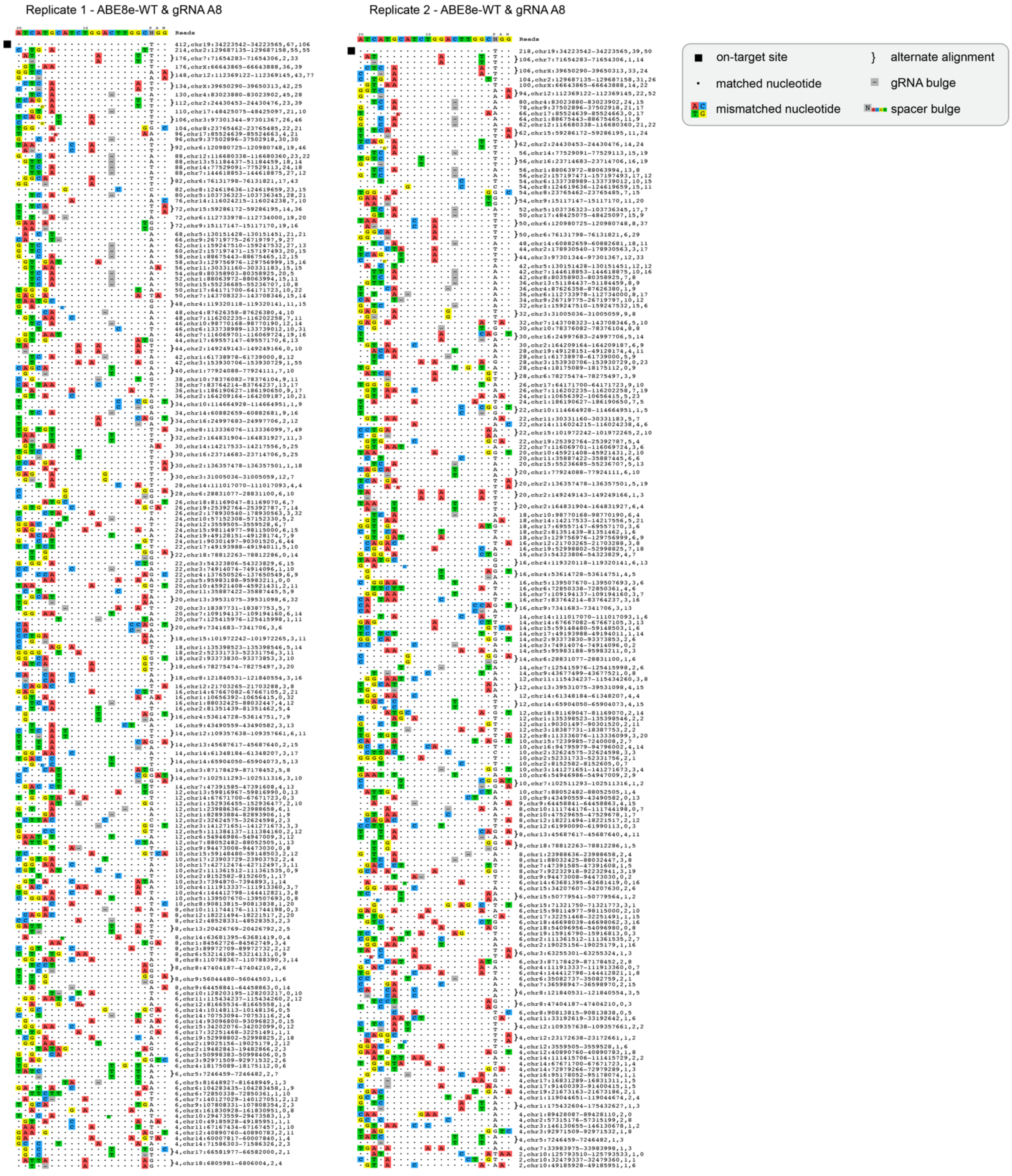


**Supplementary Figure 30. CHANGE-seq-BE detected sites in the mouse genome for ABE8e-WT paired with gRNA A8 with a 20-nucleotide spacer**. Rank-ordered visualization of on- and off-target genomic sites identified by CHANGE-seq-BE using genomic DNA from Myh11-cre:Acta2fl/+ MSMDS mice. The on-target site is indicated using a black box; alternate alignments are shown for sites that are potentially targeted via 1 nt DNA or gRNA spacer bulges; genomic locations are listed. Lower efficiency off-target sites not shown for brevity; the full set of off-target sites are available in **Supplementary Table 1**.

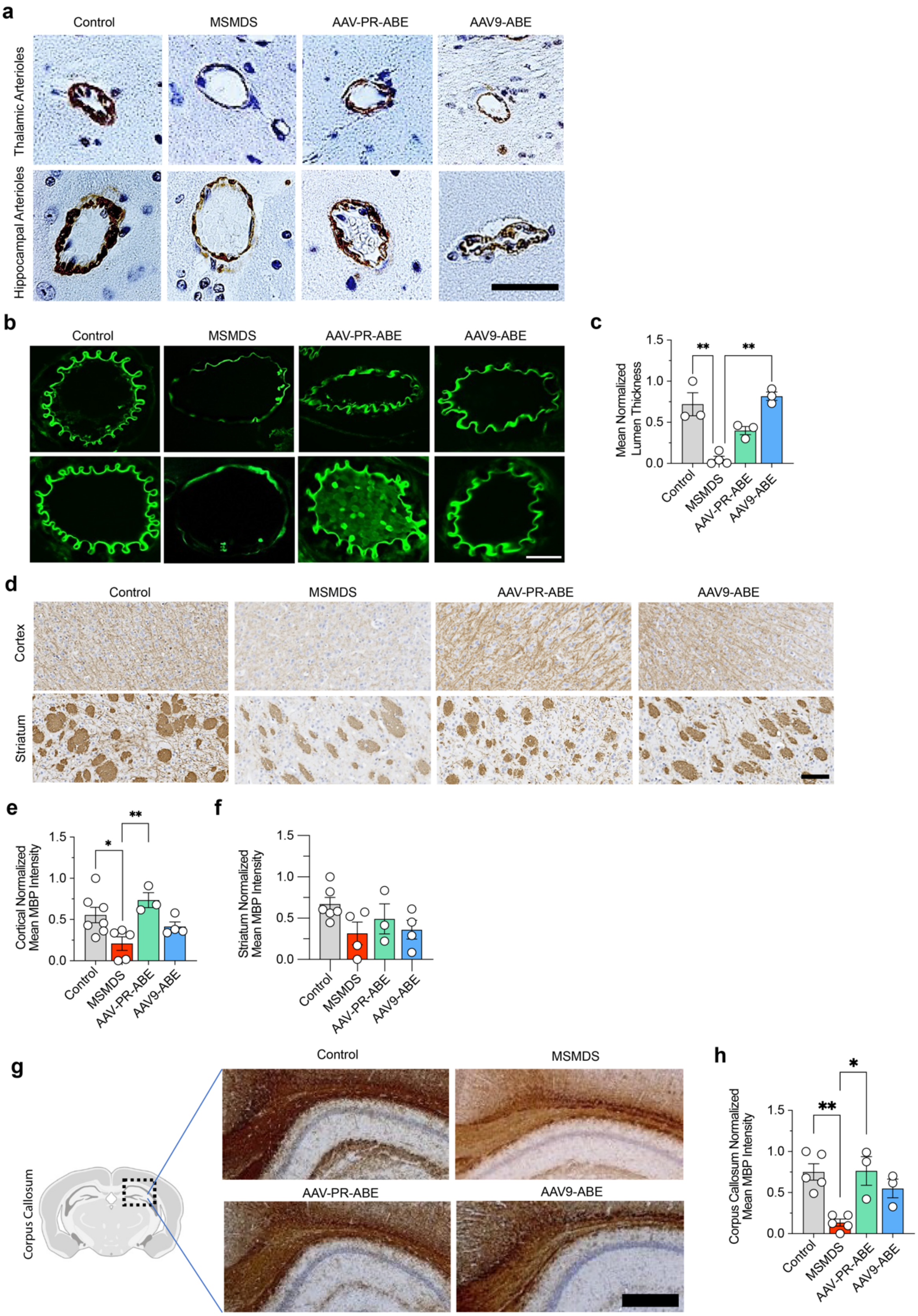


**Supplementary Figure 31. Alteration of vascular phenotypes following AAV-mediated delivery of base editors.** Analyses were performed with untreated control mice, untreated MSMDS mice, and treated MSMDS mice with AAV-PR or AAV9 viral vectors that encode ABE8e-eVRQR and gRNA-A4, with injections performed at P3. **a**, Representative images of arterioles in the thalamus (top) and hippocampus (bottom). Staining was performed using DAB-based immunohistochemistry for smooth muscle actin (SMA); scale bar: 20 µm. **b,c,** Representative autofluorescence images of elastin (**panel b**) and quantification of lumen thickness in arterioles (**panel c**), illustrating structural differences in vessel walls. Scale bar in **panel b**, 20 µm. **d**, Representative images with DAB-based MBP staining in the cortex (top panels) and striatum (bottom panels); scale bar: 50 µm. **e,f,** Quantification of MBP intensity in the cortex (**panel e**) and striatum (**panel f**) from images in **panel d**. **g,** Representative images of sagittal sections stained with DAB for MBP; scale bar: 100 µm. **h**, Quantification of MBP levels in the corpus callosum from images in **panel g**. One-way ANOVA followed by Fisher's exact test revealed significant difference in **panels c**,**e**,**f,h**; *P <0.05, **P <0.01; mean, s.e.m., and individual datapoints shown in panels shown in **panels b**,**e**,**f,h**.

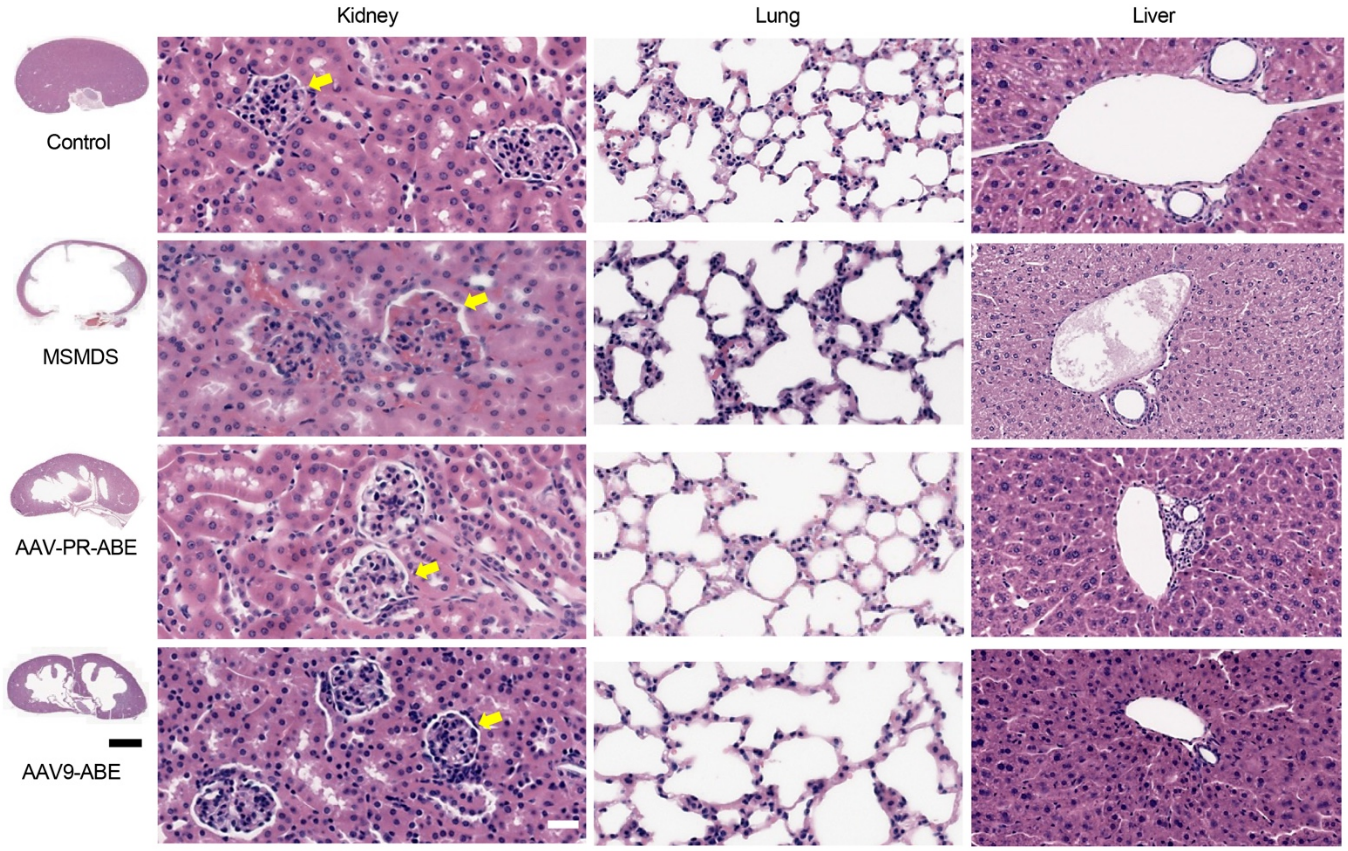


**Supplementary Figure 32. Histological changes following AAV-mediated delivery of base editors.** Representative histological images of kidney, lung, and liver via hematoxylin and eosin (H&E) staining of sections from untreated control mice, untreated MSMDS mice, and MSMDS mice treated with AAV-PR or AAV9 vectors encoding ABE8e-eVRQR and gRNA-A4, delivered at P3. Kidney: Large kidney image scale bar: 2 mm; small scale bar: 40 μm. Yellow arrows indicate glomeruli. Lung scale bar: 20 μm. Liver: Sectioned liver tissue showing vascular structures, scale bar: 25 μm.

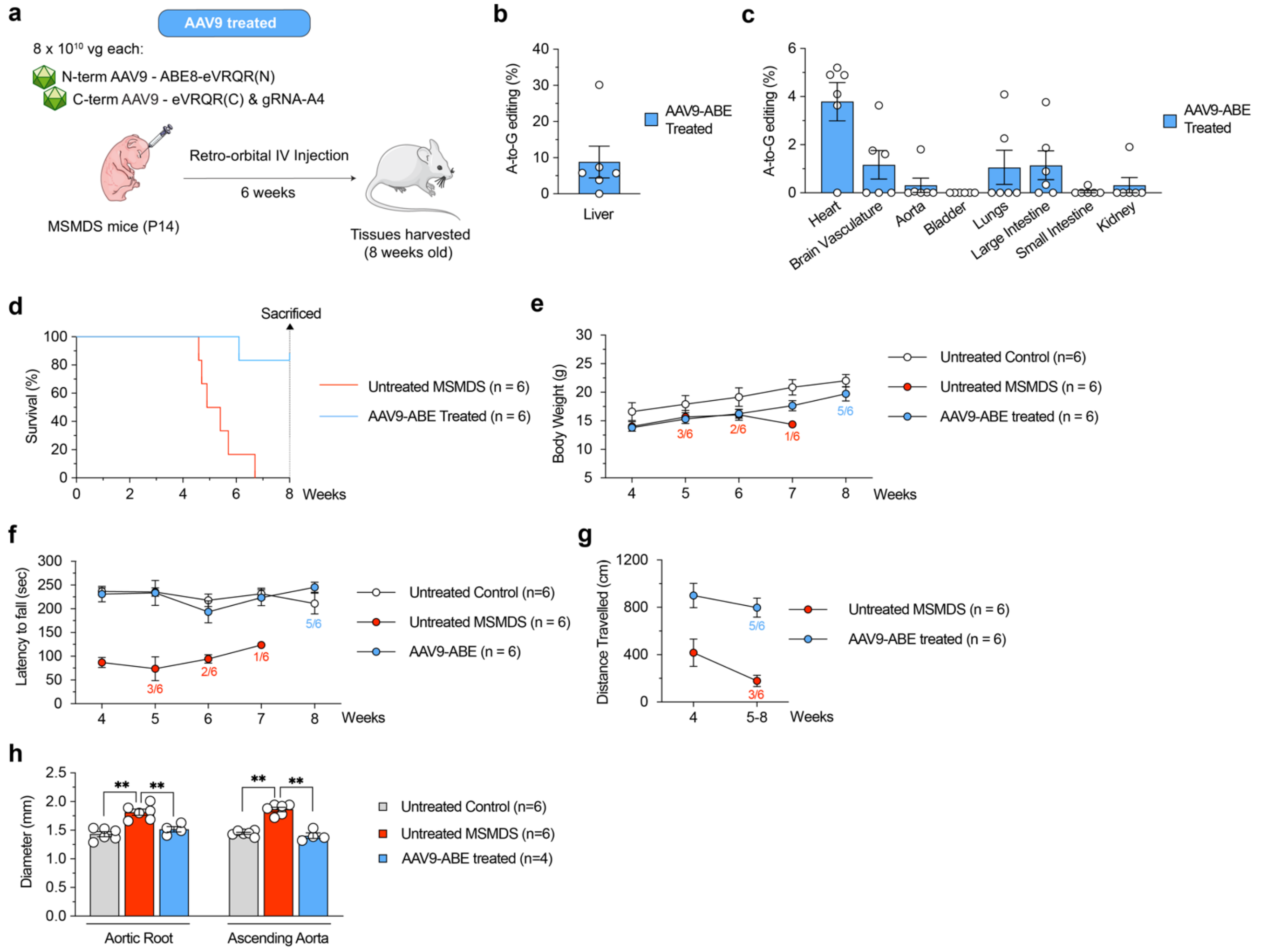


**Supplementary Figure 33. AAV9-mediated delivery of customized base editors at P14 for *in vivo ACTA2* editing**. **a**, Schematic of P14 intravenous (IV) injections in MSMDS (Myh11-Cre:Acta2fl/+) and control (*Acta2fl*/+) mice with dual AAV-PR and AAV9 vectors that express intein-split ABE8e-eVRQR and gRNA A4. Multiple tissues were harvested 6 weeks after P14 injections (from 8-week-old mice). **b,c**, On-target A-to-G editing of *ACTA2* R179H in the liver (**panel b**) and other tissues (**panel c**) from MSMDS mice sacrificed at 8 weeks of age, six weeks after AAV9-ABE treatment at P14. **d**, Survival of a cohort of untreated and AAV9-ABE treated MSMDS mice followed for 8 weeks. **e-h**, Systemic phenotypic characterization of untreated control (*Acta2fl*/+), untreated MSMDS, and AAV9-ABE treated MSMDS mice at various timepoints up to 8 weeks of age following P14 injections, including: body mass (**panel e**), rotarod performance (**panel f**), distance traveled in open field testing (**panel g**), and aortic diameter assessed by ultrasound (**panel h**). Log-rank test revealed a significant difference ($P < 0.001$) between treated and untreated mice survival in **panel d**. Repeated measures ANOVA revealed significant differences ($P < 0.01$) between treated and untreated MSMDS mice in **panels f,g**. One-way ANOVA followed by Fisher's exact test revealed significant difference in **panel h**. **P <0.01. Mean and s.e.m. shown in **panels b-h**. Sample size indicated in **panels d-g**, and individual datapoints shown in **panels b,c,h**.

#### Supplementary References


1. Anzalone, A. V. *et al.* Search-and-replace genome editing without double-strand breaks or donor DNA. *Nature* **576**, 149–157 (2019).
2. Walton, R. T., Christie, K. A., Whittaker, M. N. & Kleinstiver, B. P. Unconstrained genome targeting with near-PAMless engineered CRISPR-Cas9 variants. *Science* **368**, 290–296 (2020).
3. Kweon, J. *et al.* Engineered Prime Editors with PAM flexibility. *Molecular Therapy* (2021) doi:10.1016/j.ymthe.2021.02.022.
4. Ramirez, S. H. *et al.* An Engineered Adeno-Associated Virus Capsid Mediates Efficient Transduction of Pericytes and Smooth Muscle Cells of the Brain Vasculature. *Human Gene Therapy* **34**, 682–696 (2023).
5. Lang, J. F., Toulmin, S. A., Brida, K. L., Eisenlohr, L. C. & Davidson, B. L. Standard screening methods underreport AAV-mediated transduction and gene editing. *Nat Commun* **10**, 3415 (2019).
6. Aksenov, S. *et al.* Current and Next Steps Toward Prediction of Human Dose for Gene Therapy Using Translational Dose-Response Studies. *Clin Pharmacol Ther* **110**, 1176–1179 (2021).
7. Zou, P. Interspecies normalization of dose-response relationship for adeno-associated virus-mediated haemophilia gene therapy-Application to human dose prediction. *Br J Clin Pharmacol* **89**, 1393–1401 (2023).
8. Salabarria, S. M. *et al.* Thrombotic microangiopathy following systemic AAV administration is dependent on anti-capsid antibodies. *J Clin Invest* **134**, e173510 (2024).
9. Gaudelli, N. M. *et al.* Directed evolution of adenine base editors with increased activity and therapeutic application. *Nat Biotechnol* **38**, 892–900 (2020).
10. Richter, M. F. *et al.* Phage-assisted evolution of an adenine base editor with improved Cas domain compatibility and activity. *Nat Biotechnol* **38**, 883–891 (2020).
11. Kleinstiver, B. P. *et al.* High-fidelity CRISPR-Cas9 nucleases with no detectable genome-wide off-target effects. *Nature* **529**, 490–495 (2016).
12. Kleinstiver, B. P. *et al.* Engineered CRISPR-Cas9 nucleases with altered PAM specificities. *Nature* **523**, 481–485 (2015).
13. Anders, C., Bargsten, K. & Jinek, M. Structural Plasticity of PAM Recognition by Engineered Variants of the RNA-Guided Endonuclease Cas9. *Molecular Cell* **61**, 895–902 (2016).
14. Spencer, J. M. & Zhang, X. Deep mutational scanning of S. pyogenes Cas9 reveals important functional domains. *Sci Rep* **7**, 16836 (2017).
15. Hibshman, G. N. *et al.* Unraveling the mechanisms of PAMless DNA interrogation by SpRY-Cas9. *Nat Commun* **15**, 3663 (2024).
16. Walton, R. T., Hsu, J. Y., Joung, J. K. & Kleinstiver, B. P. Scalable characterization of the PAM requirements of CRISPR-Cas enzymes using HT-PAMDA. *Nat Protoc* **16**, 1511–1547 (2021).
17. Gaudelli, N. M. *et al.* Programmable base editing of A•T to G•C in genomic DNA without DNA cleavage. *Nature* **551**, 464–471 (2017).
18. Koblan, L. W. *et al.* Improving cytidine and adenine base editors by expression optimization and ancestral reconstruction. *Nat Biotechnol* **36**, 843–846 (2018).
19. Cas-OFFinder: a fast and versatile algorithm that searches for potential off-target sites of Cas9 RNA-guided endonucleases - PubMed. https://pubmed.ncbi.nlm.nih.gov/24463181/.

20. Lazzarotto, C. *et al.* CHANGE-seq-BE enables simultaneously sensitive and unbiased in vitro profiling of base editor genome-wide activity. 2024.03.28.586621 Preprint at https://doi.org/10.1101/2024.03.28.586621 (2024).

21. Clement, K. *et al.* CRISPResso2 provides accurate and rapid genome editing sequence analysis. *Nature Biotechnology* **37**, 224–226 (2019).

22. Gaudelli, N. M. *et al.* Programmable base editing of A•T to G•C in genomic DNA without DNA cleavage. *Nature* **551**, 464–471 (2017).

23. Gaudelli, N. M. *et al.* Directed evolution of adenine base editors with increased activity and therapeutic application. *Nat Biotechnol* **38**, 892–900 (2020).
